# Reorganization of innate immune cell lipid profiles by bioinspired meroterpenoids to limit inflammation

**DOI:** 10.1101/2024.05.24.595516

**Authors:** Lorenz Waltl, Klaus Speck, Raphael Wildermuth, Franz-Lucas Haut, Stephan Permann, Danilo D’Avino, Ida Cerqua, Anita Siller, Harald Schennach, Antonietta Rossi, Thomas Magauer, Andreas Koeberle

## Abstract

Lipidomics-guided screening of unexplored chemical space in natural products provides access to small molecules capable of modifying cellular lipid profiles on a global scale. Here, we show that the meroterpenoid cyclosmenospongine from *Spongia sp*. shapes the lipid profile of immune cells, favoring anti-inflammatory and pro-resolving over pro-inflammatory lipid mediators. Structural variation revealed derivatives that inhibit leukotriene biosynthesis to varying extents while differentially upregulating pro-resolving lipid mediators, epoxyeicosatrienoic acids, endocannabinoids, and sphingosine-1-phosphate, along with other mediators, both in resting and activated innate immune cells *in vitro* and in self-resolving murine peritonitis *in vivo*. Mechanistically, meroterpenoids target 5-lipoxygenase or 5-lipoxygenase-activating protein, promote the translocation of 15-lipoxygenase-1 to cytoplasmatic sites, and inhibit monoacylglycerol lipase. They also redirect arachidonic acid (AA) from neutral lipids to specific phospholipids, while increasing the total concentration of free AA. Furthermore, meroterpenoids reprogram lipid metabolism in immune cells, decreasing the levels of neutral lipids, triacylglycerols, and cholesteryl esters. This shift correlates with a reduced capacity for leukotriene biosynthesis and is mimicked by the inhibition of sterol-O-acyltransferase and diacylglycerol acyltransferase-1/2. In conclusion, specific meroterpenoids exert anti-inflammatory effects by intervening in lipid mediator biosynthesis, prompting structure-controlled switches in lipid mediator classes, among others, through an unexpected link between lipogenesis and inflammation.

## 1 Introduction

Inflammation is a tightly regulated physiological response to tissue damage and infection aimed at restoring homeostasis.^[1]^ Failure to dampen the pro-inflammatory reaction in the resolution phase can lead to low-grade persistent inflammation that contributes to a wide range of chronic disorders, including metabolic, autoimmune, neurodegenerative, cardiovascular, and malignant diseases.^[2]^ The initiation, propagation, and termination of inflammation are orchestrated by a plethora of mediators, including cytokines and lipids. The latter are a highly diverse class of biomolecules comprising thousands of different molecules,^[3]^ among them, lipid mediators with pro-inflammatory, anti-inflammatory, pro-resolving, or immunomodulatory activity, which act as ligands for membrane or intracellular receptors.^[2a, 4]^ In addition, links between cellular metabolism, immune function, and persistent inflammation^[5]^ are emerging, blurring the line between structural lipids and signaling molecules, as exemplified by the recent discovery of a lipokine-like, stress-protective phosphatidylinositol (PI).^[6]^

Immune and non-immune cells produce lipid mediators from polyunsaturated fatty acids (PUFAs), which are released from membrane phospholipids by members of the phospholipase A_2_ (PLA_2_) superfamily.^[7]^ Free PUFAs are then converted into pro-inflammatory (prostaglandins and leukotrienes [LTs]), anti-inflammatory (fatty acid epoxides), or specialized pro-resolving lipid mediators (SPMs) by di- and monooxygenases in concert with additional biosynthetic enzymes.^[2a, 2b, 4e-g, 4i]^ Cyclooxygenase (COX) isoenzymes are the primary targets of non-steroidal anti-inflammatory drugs (NSAIDs).^[4i]^ They convert the ω6-fatty acid arachidonic acid (AA (20:4)) to prostaglandin H_2_ (PGH_2_), from which prostaglandin E_2_ (PGE_2_) and other prostanoids are generated. PGE_2_ drives inflammation, fever, and pain, but it also has homeostatic and anti-inflammatory activity and initiates the switch to resolution of inflammation, depending on its (spatial) concentration and kinetics.^[4i, 8]^

5-Lipoxygenase (5-LOX) translocates to the perinuclear membrane, accepts AA (20:4) from 5-lipoxygenase-activating protein (FLAP), and initiates the production of LTs to attract and activate immune cells, increase vascular permeability, and induce bronchial constriction,^[2b, 4g, 4i, 9]^ but also synthesizes lipoxins (LXs) in combination with 12-or 15-LOX.^[10]^ The isoenzyme 15-LOX-1 undergoes a Ca^2+^-dependent translocation to poorly defined cytoplasmic compartments in activated M2 macrophages^[11]^ and has been recognized as a key regulated enzyme for SPM biosynthesis.^[12]^ It accepts ω3-fatty acids and, together with lipoxygenases of different regiospecificity or (partially unknown) oxidoreductases, produces SPMs, such as LXs, protectins (PDs), resolvins (Rvs), and maresins (MaR),^[2a, 2b, 4e, 4f, 11-12]^ which have been proposed to actively promote the resolution of inflammation, possibly via G-protein-coupled receptors.^[2a, 13]^ SPMs suppress leukocyte infiltration and activation, in part by interfering with nuclear factor (NF)-κB signaling, which drives the expression of pro-inflammatory cytokines and chemokines.^[14] [13]^ They also promote phagocytosis, bacterial clearance, efferocytosis, and tissue repair.^[2a, 12]^ Whether SPMs reach effective concentrations *in vivo* is controversially discussed.^[15]^

The diversity of lipid mediators is further increased by the 2C and 2J families of cytochrome P_450_ monooxygenases (CYP_450_) enzymes, which are primarily responsible for the epoxidation of AA (20:4) and other PUFAs.^[16]^ Among many other functions, these epoxy-fatty acids interfere with immune cell activation, cytokine release, and COX-2 expression.^[16b]^ PUFAs can also be conjugated to produce intracellular and extracellular ligands, such as endocannabinoids, which together with other (lipid) factors, such as sphingosine-1-phosphate (S1P) orchestrate immune cell activation, lymphocyte proliferation, chemotaxis, cytokine production,^[17]^ regulate nociception^[17c, 18]^ and are closely associated with cellular metabolism and metabolic diseases. ^[4d, 19]^

Current anti-inflammatory therapies are largely based on glucocorticoids and NSAIDs. While the former suppress the production of cytokines and lipid mediators in an undifferentiated manner, the latter target COX isoenzymes and are afflicted with multiple and in some cases severe adverse effects such as ulceration, bleeding, and cardiotoxicity, caused by an impaired biosynthesis of homeostatic prostanoids and shunting of AA (20:4) to LT biosynthesis.^[4i, 20]^ New directions in anti-inflammatory drug development aim to selectively target disease-related mediators^[21]^ and induce resolution of inflammation,^[2c, 12]^ which is considered particularly valuable for the treatment of chronic inflammatory diseases. To date, only a few compounds have been identified that are capable of enhancing SPM biosynthesis in activated immune cells (up to 10-fold for MaR1 by 8-methylsocotrin-4’-ol)^[22]^ and even fewer have been shown to induce SPM biosynthesis in non-activated cells (up to 70-fold for RvD5 by cannabidiol).^[22a, 22b, 22f, 23]^ The control of beneficial lipid mediator profiles on a larger scale is still in its infancy, but can build on recent developments of dual and multiple inhibitors of lipid mediator metabolism.^[8b, 21, 24]^

In this study, we employed lipidome-wide screening to discern unexplored natural products within underrepresented chemical space and identified the meroterpenoid cyclosmenospongine (**1**; **Table 1**) to induce a shift in the lipid profile of innate immune cells toward inflammation suppression and resolution. In 2002, the *trans*-decalin **1** was isolated from the Australian sponge *Spongia sp*. (order Dictyoceratida, family Spongiidae). Subsequent studies indicated its efficacy against methicillin-resistant *Staphylococcus aureus* strains^[25]^ and showed limited inhibition of ascites cancer cell proliferation.^[25-26]^ Through modular total synthesis, we obtained access to structural derivatives of **1**,^[25]^ capable of rearranging the lipid mediator profile within innate immune cells. These derivatives are effective in a murine model of self-resolving inflammation. They inhibit LT biosynthesis to varying degrees, interfere with cytokine expression, and induce the formation of SPMs in activated M2 macrophages (up to 3.3-fold for MaR2) and resting M2 macrophages (up to 190-fold for RvD5), while also upregulating anti-inflammatory epoxyeicosatrienoic acids (EETs) and immunomodulatory lipid mediators, such as PGE_2_, endocannabinoids, and S1P. Mechanistically, structural derivatives inhibit 5-LOX or antagonize FLAP, release PUFAs from distinct lipid pools, induce subcellular re-localization of 15-LOX-1, and inhibit monoacylglycerol lipase, an endocannabinoid-degrading enzyme.^[4d]^ The derivatives also reprogram cellular metabolism by depleting neutral lipids, thus establishing a previously proposed link between lipid metabolism and pro-inflammatory lipid mediator biosynthesis in human innate immune cells. Notably, even minor structural alterations can selectively influence the preferential formation of specific SPM subclasses, underscoring the directional manipulability of lipid mediator profiles.

**Table 1.**
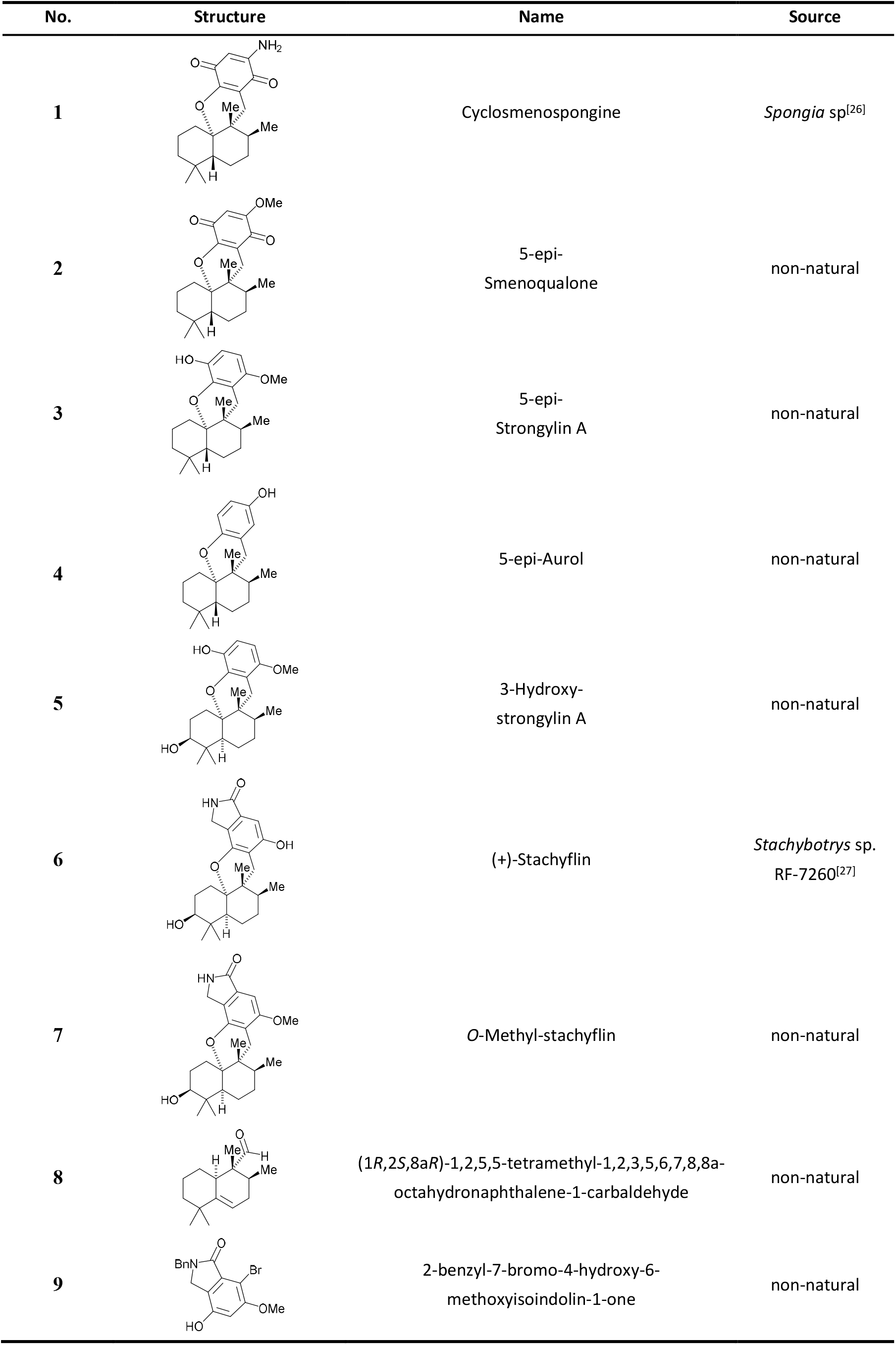
Structure of cyclosmenospongine (1) and related natural or bioinspired compounds.

| No. | Structure | Name | Source |
| --- | --- | --- | --- |
| 1 |  | Cyclosmenospongine | *Spongia* sp<sup>[26]</sup> |
| 2 |  | 5-epi-Smenoqualone | non-natural |
| 3 |  | 5-epi-Strongylin A | non-natural |
| 4 |  | 5-epi-Aurol | non-natural |
| 5 |  | 3-Hydroxy-strongylin A | non-natural |
| 6 |  | (+)-Stachyflin | *Stachybotrys* sp.<br>RF-7260<sup>[27]</sup> |
| 7 |  | O-Methyl-stachyflin | non-natural |
| 8 |  | (1*R*,2*S*,8*aR*)-1,2,5,5-tetramethyl-1,2,3,5,6,7,8,8a-octahydronaphthalene-1-carbaldehyde | non-natural |
| 9 |  | 2-benzyl-7-bromo-4-hydroxy-6-methoxyisoindolin-1-one | non-natural |

## 2 Results

### 2.1 Compound 1 induces a lipid mediator class switch from LTs to SPMs

To identify small molecules that suppress pro-inflammatory while inducing pro-resolving lipid mediator production, we initially screened natural products in A23187-activated human peripheral blood mononuclear cells (PBMCs), which consist mainly of monocytes but also contain dendritic cells, lymphocytes, and natural killer cells.^[28]^ None of the compounds investigated here exhibited cytotoxic effects within 48 h, as assessed by measuring mitochondrial dehydrogenase activity, membrane integrity, and viable cell numbers (**Figure S1A-D**, Supporting Information). The HIT compound **1** (Table 1) efficiently suppressed the formation of 5-LOX products (LTB_4_, LTB_4_-isomers, 20-hydroxy-LTB_4_ [20-OH-LTB_4_], 5-hydroxyeicosatetraenoic acid [5-HETE], 5-hydroxyeicosapentaenoic acid [5-HEPE] and 5*S*,6*R*-dihydroxy-HETE [5*S*,6*R*-diHETE]) while increasing the production of 12- and 15-LOX products (**Figure 1A** and **Figure S1E**, Supporting Information), including precursors of specialized pro-resolving mediators (SPMs), i.e., 15-HETE and 14-hydroxydocosahexaenoic acid (14-HDHA) (**Figure 1B** and **Figure S1F**, Supporting Information).

**Figure 1.**
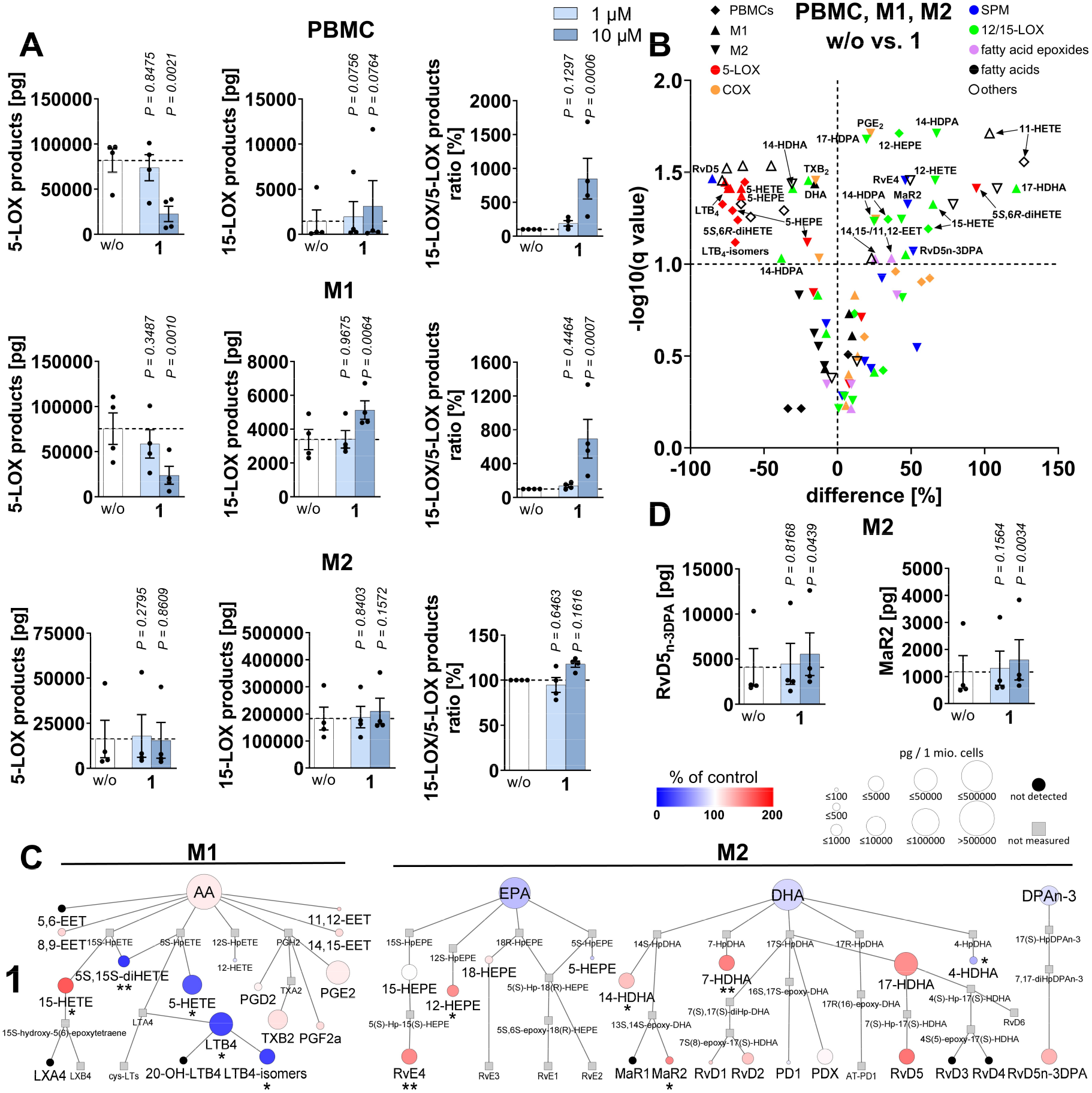
Compound **1** switches the lipid mediator profile of activated innate immune cells towards inflammation suppression and resolution. A,B) PBMCs were preincubated (10 min) with vehicle (DMSO, 0.1%) or **1** and then challenged with A23187 (10 min). A-D) M1 or M2 macrophages were preincubated (15 min) with vehicle (DMSO, 0.1%) or **1** (10 µM if not indicated otherwise) and stimulated with SACM (180 min). A) 5- and 15-LOX products [per 5×10^6^ PBMCs or 2×10^6^ M1/M2]. B) Volcano plot showing the mean percentage difference relative to vehicle control and the negative log_10_(q value) calculated vs. vehicle control; two-tailed multiple paired Student *t* tests with correction for multiple comparisons (false discovery rate 10%). C) Pathway diagram showing percentage changes in lipid mediator formation upon treatment with compound **1** (10 µM) relative to vehicle control. D) SPMs upregulated by **1** [per 2×10^6^ M2]. Mean (B,C) or mean + SEM and single data (A,D) from n = 3-4 (B,C), n = 4 (A,D) independent experiments. *P* values given vs. vehicle control (A-D) or \**P* < 0.05, \*\**P* < 0.01 (C); repeated measures one-way ANOVA of log data + Dunnett *post hoc* tests (A,D) or two-tailed paired Student *t* test of log data (C).

Since PBMCs have a low capacity to produce 5-LOX-derived LTs and 15-LOX-derived SPMs as compared to more specialized immune cells such as macrophages,^[11, 22b-d]^ we focused further studies on activated M1 and M2 macrophages. To this end, we polarized human monocyte-derived macrophages and challenged them with *Staphylococcus aureus* (LS1)-conditioned medium (SACM) as a physiological stimulus.^[29]^ M1 macrophages participate in the initiation of inflammation and efficiently produce LTs and COX-derived prostanoids, whereas M2 macrophages strongly express 15-LOX-1 and generate a broad spectrum of SPMs,^[11]^ thereby promoting the resolution of inflammation.^[12-13]^ As expected from our results in PBMCs, compound **1** induced a switch from 5-LOX to 15-LOX product formation in M1 macrophages (Figure 1A,**C**; Figure S1F and **Figure S2**, Supporting Information) and to 12-LOX and less 15-LOX product formation in M2 macrophages (Figure 1A-C; Figure S1F,**G** and Figure S2, Supporting Information). As a consequence, SPM levels were increased by trend in M2 macrophages (*P* = 0.0810) (Figure S1G, Supporting Information) and reached significance for MaR2, RvE4 and RvD5_n-3DPA_ (Figure 1B-D and Figure S1F, Supporting Information). Activation of phospholipases A_2_ can be ruled out as a primary mechanism promoting 12- and 15-LOX product formation, as levels of free PUFAs were not or barely upregulated (Figure 1C and **Figure S1H**, Supporting Information).

The increase in 15-LOX product formation in PBMCs was accompanied by enhanced biosynthesis of COX products such as PGE_2_ (Figure 1B and Figure S1E, Supporting Information), which in addition to inducing inflammation and pain also has anti-inflammatory activities and initiates the switch to inflammation resolution. It appears that AA (20:4) is partially diverted from 5-LOX product formation to other lipid mediator biosynthetic pathways, including prostaglandin biosynthesis.^[4i]^ Effects on COX products were less pronounced or absent in macrophages (Figure 1B,C; Figure S1F and Figure S2, Supporting Information).

Together, compound **1** establishes a pro-resolving lipid mediator profile in innate immune cells by inducing the biosynthesis of SPMs including 12-LOX- and 15-LOX-derived precursors, while elevating the availability of immunomodulatory prostanoids and suppressing the formation of pro-inflammatory LTs and other 5-LOX products.

### 2.2 Structural requirements for promoting SPM production

Starting from compound **1** we explored the potential of eight structural derivatives (Table 1) to induce a lipid mediator class switch in activated PBMCs, M1 and M2 macrophages. These compounds carry modifications in the *p*-quinone moiety (**2–4**), contain a *cis*-decalin scaffold (**5**-**7**), or represent fragments from total synthesis (**8, 9**). Note that the pentacyclic meroterpenoid stachyflin (**6**) is a secondary metabolite from the mold fungus Stachybotrys *sp*. RF-7260 and has been reported to have antiviral activity.^[25, 27]^

The NH_2_ group in **1** is essential for potent inhibition of 5-LOX product formation in PBMCs and M1 macrophages (**3**). Replacement with another H-bond acceptor, i.e., a methoxy (**2**) or alcohol group (**4**), is tolerated, even in the absence of other functional groups on the upper ring for PBMCs but not for M1 macrophages (**4**) (**Figure 2A,B**; Figure S2 and **Figure S3**, Supporting Information). Conversion of *trans*-into *cis*-decalin along with the introduction of an alcohol group at the 3-position (**5**) is also accepted, as is annulation of the benzene ring with γ-lactam (**6, 7**). Interestingly, the fragment **8** with a desaturated decalin retains 5-LOX-inhibitory activity in PBMCs (not in M1), whereas the building block for the upper ring system (**9**) is barely active (Figure 2A,B; Figure S3 and **Figure S4**, Supporting Information). 5-LOX product formation is less pronounced in M2 than in M1 macrophages ^[11]^ and is also less potently inhibited by the meroterpenoids **1–7** in this phenotype (Figure S2 and **Figure S5**, Supporting Information).

**Figure 2.**
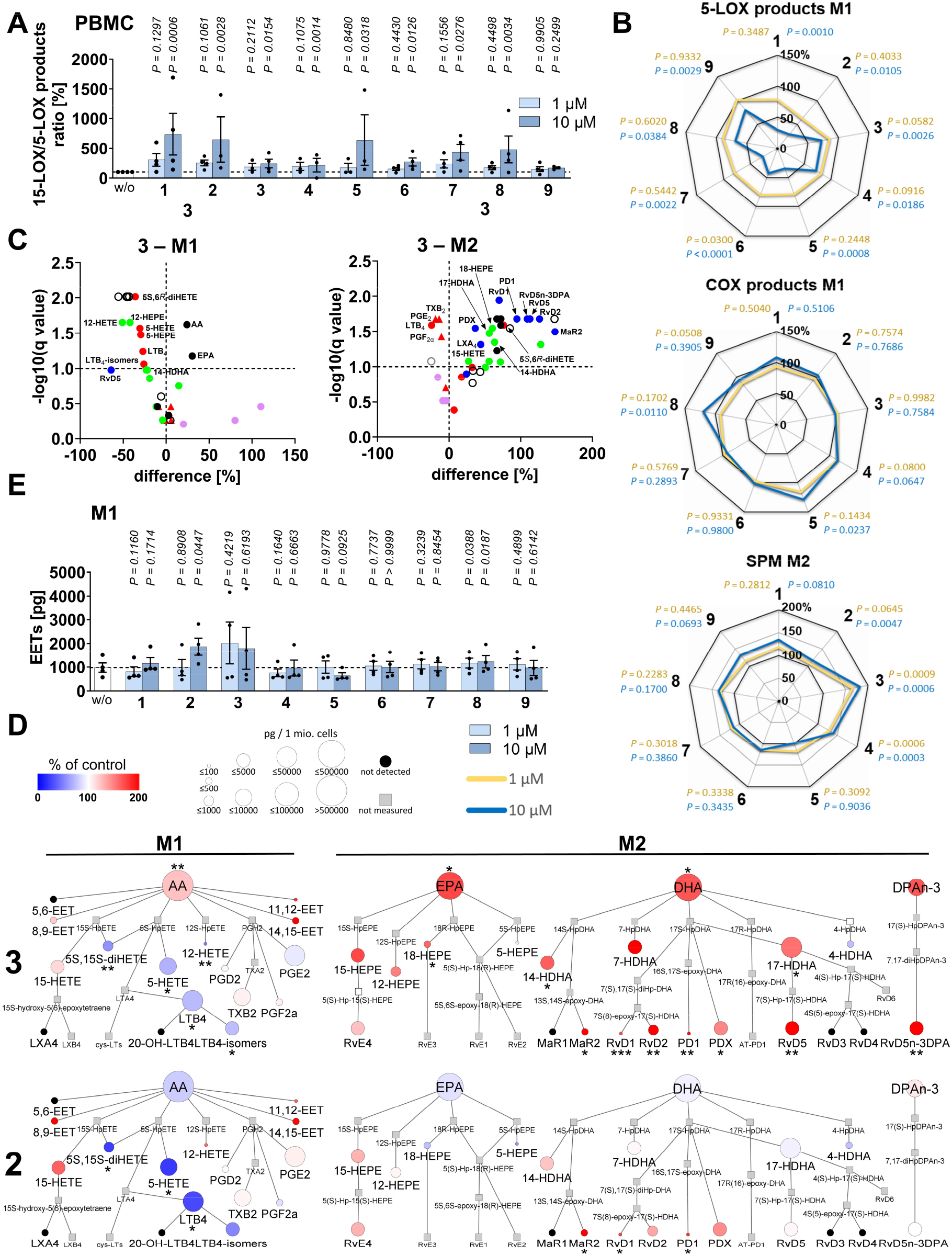
Structural meroterpenoid derivatives induce anti-inflammatory and pro-resolving changes in the lipid mediator profile of activated immune cells. A) 15-LOX/5-LOX ratio of PBMCs following 10 min preincubation with vehicle (DMSO, 0.1%) or test compounds and addition of calcium-ionophore (A23187, 10 min) [per 5×10^6^ PBMCs]. Data for vehicle control and **1** are identical to Figure 1A. B-E) M1 or M2 macrophages were preincubated (15 min) with vehicle (DMSO, 0.1%) or test compounds (10 µM if not indicated otherwise) and activated with SACM (180 min). B) 5-LOX products, COX products and SPMs. The vehicle control and data for **1** is identical to Figure 1B,C (SPMs). C) Volcano plot showing the mean percentage difference relative to vehicle control and the negative log_10_(q value) calculated vs. vehicle control; two-tailed multiple paired Student *t* tests with correction for multiple comparisons (false discovery rate 10%). D) Pathway diagrams illustrating percentage changes relative to vehicle control in the lipid mediator profile of M1 and M2 macrophages treated with **2** (10 µM) or **3** (10 µM). E) EET levels [per 2×10^6^ M1]. Mean (B,C,D) or mean + SEM and single data (A,E) from n = 3-4 independent experiments. *P* values given vs. vehicle control (A-E) or \**P* < 0.05, \*\**P* < 0.01, \*\*\**P* < 0.001; repeated measures one-way ANOVA of log data or mixed-effects model (REML) of log data + Dunnett *post hoc* tests (A,B,E) and two-tailed paired Student *t* test of log data (D).

The ratio of 15-LOX to 5-LOX products increases in PBMCs treated with **1** or derivatives (except for the truncated **9**) (Figure 2A), with *trans*-decalins (**1–4**) preferentially upregulating 15-LOX-derived products in addition to or instead of suppressing 5-LOX product formation (Figure S1E and S3, Supporting Information). Similar results were obtained in M2 macrophages, where the increase in 15-LOX and 12-LOX product formation was translated into enhanced SPM biosynthesis (**3** > **4** > **1, 2**) (Figure 2B,**C**; Figure S5 and **Figure S6A**, Supporting Information). Interestingly, structural variants allow partial discrimination between SPM subclasses (Figure S2 and Figure S6A, Supporting Information), possibly due to differences in subcellular distribution and thereby access to biosynthetic enzymes.^[30]^ Thus, compared to other compounds in this series, **2** is particularly efficient at increasing PD1 and MaR2 levels, **3** strongly enhances the production of all major SPMs (including Rvs, PD1, MaR2, and LXA_4_), and the otherwise less active *cis*-decalin **5** specifically induces PD1 biosynthesis (Figure S6A, Supporting Information). Mechanistically, the enhanced SPM biosynthesis by **3** appears to be (at least partially) dependent on the release of PUFAs and the formation of 12-/15-LOX-derived SPM precursors, such as 17-HDHA (D-series Rv and PD precursor), 18-HEPE (E-series Rv precursor), 14-HDHA (MaR precursor), and 15-HETE (LX precursor). The stimulatory effect of **2**, which is mainly directed to PD1, RvD1, and MaR2, is instead not accompanied by a substantially higher availability of SPM precursors or free PUFAs (**Figure 2D**; Figure S2, Figure S5 and Figure S6A, Supporting Information). These findings imply that the stimulated release of PUFAs by (specific) meroterpenoids contributes to the efficient induction of SPM biosynthesis in innate immune cells but is not the main driving mechanism. The moderate stimulatory activity of **1** on the biosynthesis of COX-derived PGE_2_ and/or other prostanoids was also evident for **2–9**, although the efficiency to induce the formation of COX products varied greatly between activated innate immune cells and prostanoids (Figure 2B; Figure S1E,F and Figure S4,5, Supporting Information). Levels of individual prostanoids (PGE_2_, PGF_2α_, and/or TxB_2_) or total COX products were significantly increased in PBMCs for **2, 3, 5–8** (Figure S3, Supporting Information), in M1 macrophages for **4, 5**, and **8** (Figure 2B and Figure S4, Supporting Information), and in M2 macrophages for **5, 7**, and **8** (Figure S5, Supporting Information). For other compounds/settings, prostanoid production was either not substantially affected or even decreased (Figure S1E and Figure S2, Supporting Information), especially in M2 macrophages treated with **3** (Figure 2C).

The ability of meroterpenoids to increase anti-inflammatory EET levels was strongly enhanced (specifically in M1 macrophages) by replacing the amine in **1** with a methoxy group (**2**), whereas other derivatives were hardly active or, in the case of **3**, gave mixed results in independent datasets based on immune cells from different platelet donors (**Figure 2E**, Figure S4 and **Figure S6B**, Supporting Information).

Conclusively, the structural modification of **1** led to derivatives that outperform the parental compound (**1**) in inducing a favorable lipid mediator class switch from LTs to SPMs (**2**-**4**) and partially also to EETs (**2**) in polarized macrophages. While compound **2** preferentially increases PD1, MaR2 and EET levels, compound **3** potently induces the biosynthesis of SPMs across subclasses (Figure 2D).

### 2.4 Meroterpenoids actively trigger SPM biosynthesis in resting macrophages

The small β-pore-forming toxin α-hemolysin from SACM induces 15-LOX product and SPM formation in M2 macrophages dependent on Metallopeptidase Domain 10 (ADAM10).^[29]^ We asked whether meroterpenoids sensitize cells to SACM or activate SPM biosynthesis by independent mechanisms. To answer this question, we challenged non-activated M2 macrophages with **2** and **3** and monitored the lipid mediator profile. Both **2** and **3** potently induced SPM production already at 1 µM, with a trend towards enhanced 12/15-LOX and COX product formation (**Figure 3A** and **Figure S7A**, Supporting Information). At higher concentrations (10 µM), both derivatives increased the availability of free PUFAs, whose release likely contributes to the overall biosynthesis of lipid mediators, but still allows a preference for 12/15-LOX products and especially SPMs (**Figure 3B** and **Figure S7B**, Supporting Information). The total amount of SPMs consequently equals or even exceeds the low basal levels of 5-LOX products, thereby potentially reaching physiologically relevant concentrations (Figure S7B, Supporting Information).

**Figure 3.**
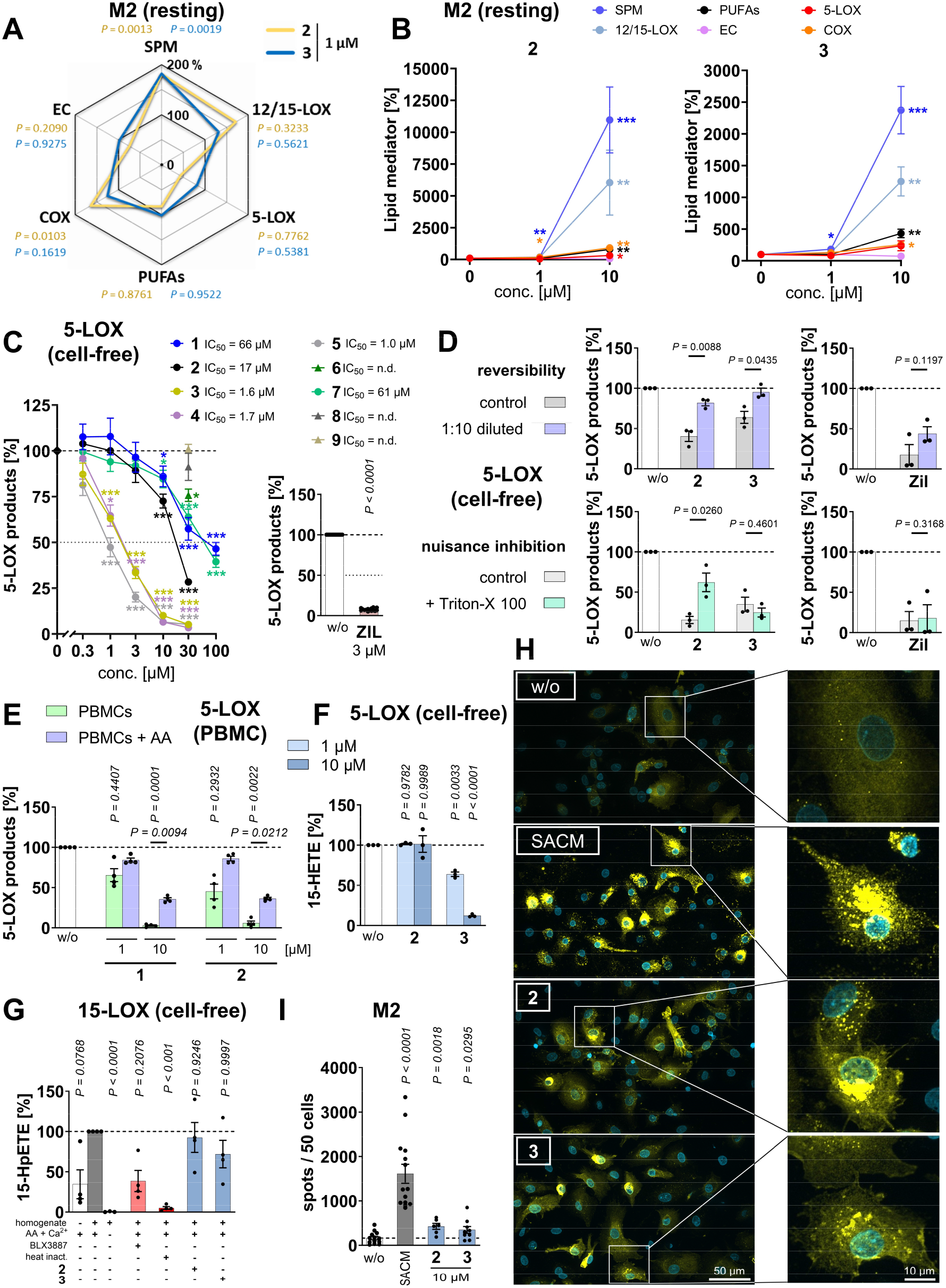
Specific meroterpenoids differentially suppress 5-LOX product formation while inducing 15-LOX-1 translocation associated with increased SPM formation in resting M2 macrophages. A,B) Percentage changes in the levels of PUFAs and lipid mediator classes produced by resting M2 macrophages treated with vehicle (DMSO, 0.1%), **2** or **3** for 195 min. The lipid mediators belonging to each subclass are defined in Figure S7A, Supporting Information. C,F) Human recombinant 5-LOX was pretreated (10 min) with vehicle (DMSO, 0.1%), zileuton (‘ZIL’) or test compounds, and product formation was initiated with AA (20:4) and CaCl_2_ (10 min). C) Concentration-dependent effect on the formation of total 5-LOX products. D) Effect of Triton X-100 (0.01%) on 5-LOX inhibition by vehicle (DMSO, 0.1%), **2** (30 µM) or **3** (3 µM). Reversibility of 5-LOX inhibition by **2** (30 µM) and **3** (3 µM). Samples were pre-treated (10 min) with vehicle, **2** or **3**, diluted 10-fold and incubated for another 2 min prior to addition of AA (20:4). E) PBMCs were preincubated (10 min) with vehicle (DMSO, 0.1%), **1** or **2**, and 5-LOX product formation (LTB_4_, LTB_4_-isomers, 20-OH-LTB_4_, 5*S*,6*R*-diHETE and 5-HEPE) was induced by calcium ionophore (A23187) in the presence or absence of AA (20:4). F) Conversion of AA (20:4) to 15-HETE by 5-LOX. G) 15-HpETE formation (indicative of 15-LOX activity) in M2 homogenates that were preincubated (15 min) with vehicle (DMSO, 0.1%) or test compounds (10 µM) and then treated with AA (20:4) and CaCl_2_ (15 min). H,I) M2 macrophages treated with vehicle (DMSO, 0.1%), SACM (0.5%), **2** (10 µM) or **3** (10 µM) for 90 min. H) Immunofluorescence images are stained for 15-LOX-1 (yellow, exposure time 300 ms) and nuclei (DAPI, light blue, exposure time 20 ms). Images are representative of > 100 single cells analyzed in n = 3 independent experiments. I) Quantification of spots resulting from 15-LOX-1 translocation, normalized to cell number. Mean (A) or mean + SEM (B,C) and single data (D-G,I) from n = 3 (C,D,F,H,I), n = 3-4 (G), n = 4 (E), n = 3-5 (A,B), n = 15 (C ZIL) independent experiments. \**P* < 0.05, \*\**P* < 0.01, \*\*\**P* < 0.001 or *P* values given vs. vehicle control or as indicated by bars; repeated measures one-way ANOVA of log data (A,C **2-5**,E,F), two-way ANOVA of log data (D) or mixed-effects model (REML) of log data (A,C **1** and **7**,G) + Dunnett *post hoc* tests and two-tailed paired Student *t* test (I) of log data (B,C **6, 8–9**, ZIL, D ZIL, E for pairwise comparisons).

### 2.3 Meroterpenoids use different strategies to inhibit 5-LOX product formation

The meroterpenoids **3–5** directly and, as shown for **3**, reversibly inhibit human 5-LOX (**Figure 3C,D**) and are even more effective in cell-free as compared to cell-based assays (Figure 2B and Figure 3C; Figure S3, Supporting Information). Non-specific interference by the formation of detergent-sensitive colloid-like aggregates^[31]^ could be excluded for **3** since Triton X-100 did not interfere with the 5-LOX inhibition (Figure 3D). All other derivatives, including the natural products **1** and **6**, instead suppressed 5-LOX product formation in innate immune cells more potently (Figure 2B and Figure 3C; Figure S3, Supporting Information), pointing to another high-affinity target within the 5-LOX pathway. The addition of Triton X-100 even implied that the moderate inhibition of cell-free 5-LOX by **2** may be an artifact (Figure 3D). Given the overall decrease in 5-LOX products, which is not limited to LTB_4_, we excluded LTA_4_ hydrolase as a target (Figure S1E and Figure S2, Supporting Information). Our results are consistent with FLAP antagonism, as exogenous AA (20:4) reduced the ability of **1** and **2** to decrease 5-LOX product formation in PBMC (**Figure 3E**). FLAP is an integral nuclear membrane protein that transfers AA (20:4) from cPLA_2α_ to 5-LOX at the nuclear membrane and is essential for 5-LOX product formation unless AA (20:4) is supplemented.^[32]^ We conclude that compound **3** is a direct and reversible inhibitor of 5-LOX, whereas **2** seems to be a FLAP antagonist.

### 2.4 Induction of SPM biosynthesis is associated with 15-LOX-1 translocation

Only a few small molecules are known to stimulate SPM biosynthesis in non-activated macrophages,^[22a, 22b, 22f, 23a]^ one of which, AKBA (3-O-acetyl-11-keto-β-boswellic acid), shifts the regiospecificity of cell-free 5-LOX toward 12/15-lipoxygenation and induces SPM biosynthesis in HEK293 cells stably transfected with 5-LOX and FLAP.^[33]^ We excluded such an activity for **2** and **3**, which not only failed to enhance the conversion of AA (20:4) to 12-HETE and 15-HETE by human recombinant 5-LOX (**Figure 3F** and **Figure S8A**, Supporting Information), but in case of **3** even significantly reduced this side reaction. We then investigated in a cell-free assay whether **2** and **3** directly activate human 15-LOX, a key enzyme in SPM biosynthesis ^[11, 22a]^, which was also not the case (**Figure 3G** and **Figure S8B**, Supporting Information).

SACM, as well as previously reported small molecules that actively trigger SPM biosynthesis, redistribute 15-LOX-1 to particulate sites in the cytoplasm.^[22a, 22b, 22h, 23a, 29]^ In this sense, **2** and **3** efficiently stimulate 15-LOX-1 translocation in M2 macrophages (**Figure 3H**) and increase the number of 15-LOX-1-enriched cytoplasmic particles (**Figure 3I**), which is considered to initiate SPM precursor formation.^[11, 34]^ It should be noted that SACM is more effective than the meroterpenoids, both in inducing 15-LOX-1 translocation (Figure 3H,I) and in triggering SPM biosynthesis, but also shows cytotoxicity upon prolonged exposure.^[29]^ In conclusion, **2** and **3** seemingly enhance SPM production in resting macrophages by coordinating the subcellular localization of 15-LOX-1 (Figure 3H,I), possibly in synergism with stimulating PUFA release at higher concentrations (Figure 3B).

### 2.5 Concomitant suppression of cytokine production

Cytokines play an important role in coordinating the inflammatory response in crosstalk with lipid mediators.^[35]^ We investigated the effect of meroterpenoids on the production of five pro-inflammatory (TNF-α, IL-1β, IL-6, IL-12 (p70), IL-23), and two anti-inflammatory cytokines (IL-10 and IL-1ra), as well as two chemokines (MCP-1 and IL-8) in lipopolysaccharide (LPS)-stimulated PBMCs. Overall, meroterpenoids (**1–7**) moderately decreased the cytokine and chemokine expression, except for MCP1, whose levels tended to increase, especially after treatment with **1** (**Figure 4A** and **Figure S9**, Supporting Information). Mechanistically, compounds **3** and by trend **2** interfere with the activation of NF-κB, a key transcription factor that induces the expression of pro-inflammatory cytokines.^[36]^ More specifically, the two meroterpenoids suppressed the LPS-induced phosphorylation of inhibitor of κ light chain gene enhancer in B cells α (p-IκBα) at 1 h after a transient increase at 15 min, without yet inducing its degradation by the proteasome (**Figure 4B** and **Figure S10**, Supporting Information). Interestingly, derivatives **3**-**5** and **7** are particularly effective in reducing the levels of specific cytokines. For example, compounds **3**-**5** strongly and preferentially decreased the synthesis of IL-1β and either IL-12 (**3, 4**), IL-23 (**4**), or IL-1ra (**5**) (**Figure 4C** and Figure S9, Supporting Information), suggesting that additional cytokine-modulating mechanisms besides impaired NF-κB signaling are involved. In conclusion, meroterpenoids downregulate cytokine expression by interfering with LPS-triggered NF-κB signaling, but possibly also by other mechanisms. While these results are consistent with impaired immune cell activation, it remains elusive whether the reduced cytokine production is secondary to the preceding lipid mediator class switch.

**Figure 4.**
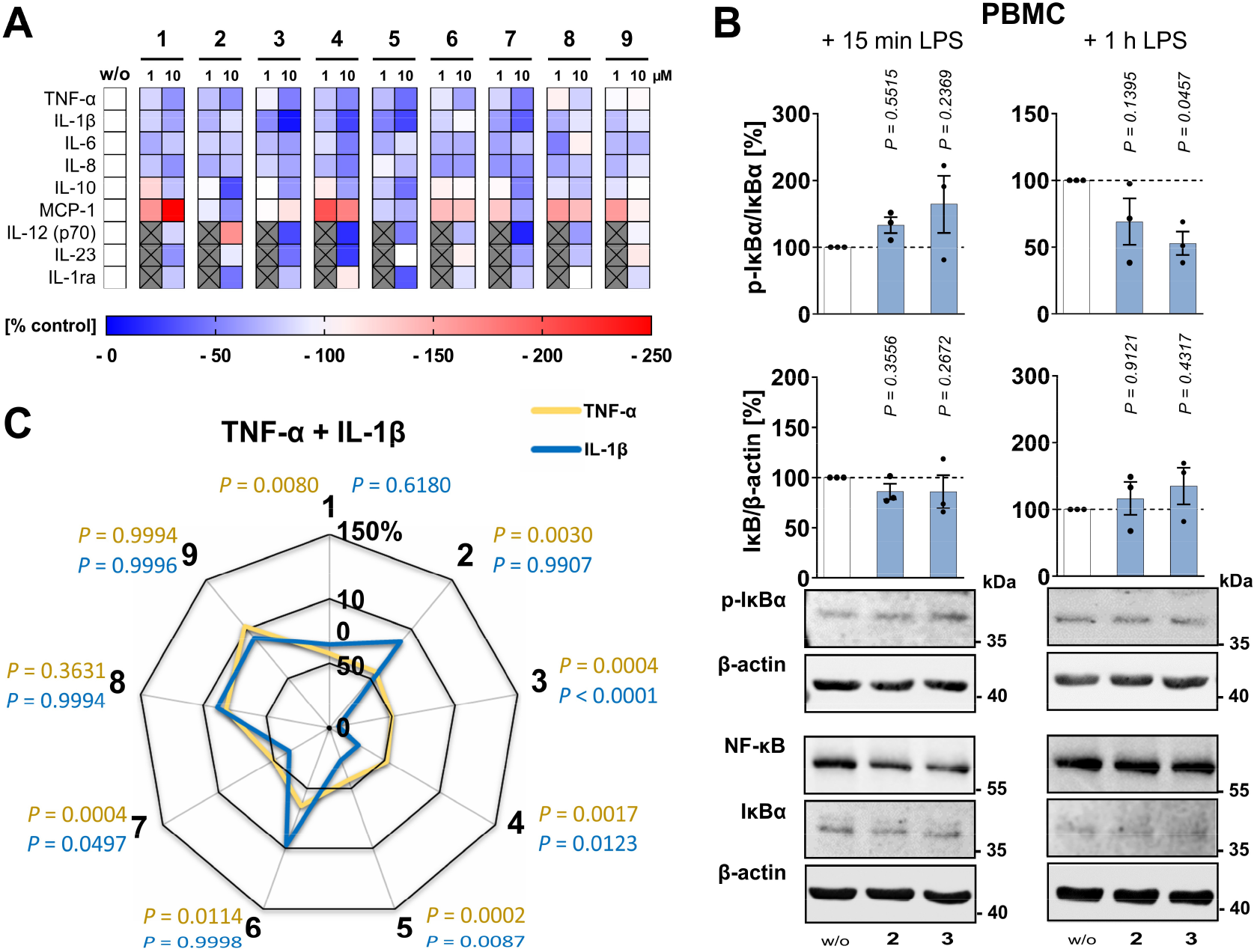
Meroterpenoids interfere with the production of inflammatory cytokines and chemokines. PBMCs were pretreated (30 min) with vehicle (DMSO, 0.1 %) or test compounds (10 µM if not indicated otherwise) and then stimulated with LPS for 4 h (TNF-α, IL-8), 18 h (IL-1β, IL-6, IL-10, MCP-1, IL-12 (p70), IL-23, IL-1ra) or as indicated. A) Heatmap showing changes in the cytokine and chemokine profile as percentage of vehicle control. B) Cellular IκBα phosphorylation, IκBα levels and NF-κB expression. Western blots are representative for n = 3 independent experiments. Uncropped blots are shown in **Figure S11A**,**B**, Supporting Information (A 15 min LPS, B 1 h LPS). C) Radar chart indicating percentage changes in TNF-α and IL-1β expression versus vehicle control. Mean (A,C) or mean + SEM and single data (B) from n = 3 (B,C), n = 3-4 (A) independent experiments. *P* values given vs. vehicle control (B,C); repeated measures one-way ANOVA (B) of log data (C IL-1β) or mixed-effects model (REML) of log data (C TNF-α) + Dunnett *post hoc* tests and two-tailed paired Student *t* test for pairwise comparisons as indicated by bars.

### 2.6 Changes in the lipid mediator profile in relieved murine peritonitis

Whether **2** and **3** induce a lipid mediator class switch from inflammation to resolution *in vivo* was investigated for zymosan-induced murine peritonitis (**Figure 5A**). Initiation, progression, and resolution of inflammation in this self-resolving model^[37]^ depend essentially, in addition to cytokines^[38]^, on lipid mediators, specifically i) LTs^[39]^ and prostaglandins^[40]^ which promote inflammation, ii) EETs^[41]^ and SPMs^[38a, 42]^ which limit or resolve inflammation, and iv) S1P^[4b, 43]^ and potentially endocannabinoids^[44]^ with immunomodulatory function. First, we determined the availability of **2** and **3** in plasma and peritoneal exudate upon i.p. injection during both acute (4 h) and resolving inflammation (18 h). Compound **2** is present in the peritoneal exudate and the systemic circulation at 4 h, but the levels then rapidly decrease, whereas **3** is maintained in the exudate and plasma even 18 h after zymosan injection (**Figure S12A**, Supporting Information).

**Figure 5.**
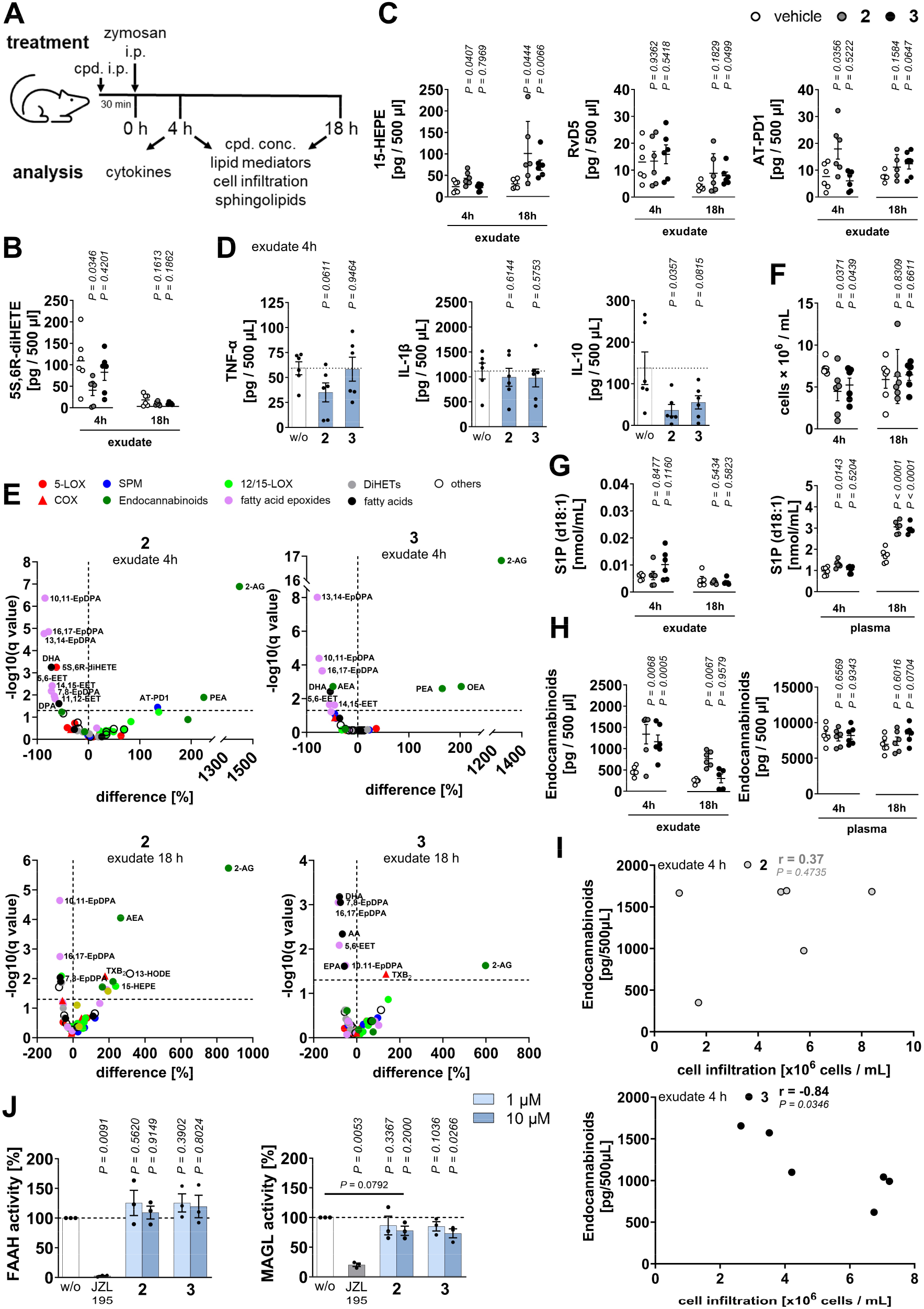
Alleviation of *in vivo* inflammation by **2** and **3** in murine peritonitis is associated with the reorganization of systemic and exudate lipid mediator profiles. A) Time course and readouts for zymosan-induced murine peritonitis. B-I) Mice received vehicle (2% DMSO in saline; 0.5 mL), **2** or **3** (10 mg/kg, i.p.) 30 min prior to zymosan injection and were sacrificed after 4 h or 18 h for the analysis of lipid mediators and cytokines in plasma and exudate (B-E, G-I) and the determination of cell infiltration into the peritoneal cavity (F). B) 5*S*,6*R*-diHETE. C) 18-HEPE, RvD5, and AT-PD1. D) TNF-α, IL-1β, and IL-10. E) Volcano plots showing the mean percentage difference relative to vehicle control and the negative log_10_(q value) calculated vs. vehicle control; two-tailed multiple paired Student *t* tests with correction for multiple comparisons (false discovery rate 5%). F) Immune cell infiltration. G) Sphingosine-1-phosphate (S1P) (d18:1). H) Total endocannabinoids. I) Pearson correlation between peritoneal cell infiltration and total endocannabinoid concentration calculated from the mean cell number and mean endocannabinoid concentration. J) Human recombinant FAAH and MAGL were pretreated (FAAH: 5 min; MAGL: 15 min) with vehicle (DMSO, 0.1%), JZL195 (FAAH: 1 µM; MAGL: 4.4 µM) or test compounds before the enzymatic activity was determined by the hydrolysis of substrates yielding fluorescent or colored products. Mean (E,I) or mean + SEM and single data (B-D,F-H,J) from n = 5-6 (B,C,E,G), n = 6 (D,F,H,I) mice or from n = 3 (J) independent experiments. *P* values given vs. vehicle control (B-H,J); two-tailed unpaired Student *t* test (B-D,F,H) of log data (G) or repeated measures one-way ANOVA of log data (J) and two-tailed paired Student *t* test of log data for pairwise comparisons as indicated by bars (J). The lipid mediators belonging to each subclass are defined in Figure S13, Supporting Information.

Compound **2** reduced the exudate concentration of 5-LOX products such as 5*S*,6*R*-diHETE. However, the effects were limited to certain species, and **3** failed in this respect (**Figure 5B**; **Figure S12B** and **Figure S13**, Supporting Information). This trend towards reduced 5-LOX product levels was more pronounced in plasma during the resolution phase (18 h) (**Figure S12C**, Supporting Information). On the other hand, specific 12/15-LOX products and SPMs were enriched, reaching significance for 15-HEPE (**2**, 4+18 h; **3**, 18 h), RvD5 (**3**, 18 h), and AT-PD1 (**2**, 4h; **3**, 18 h) preferentially in the exudate (**Figure 5C** and **Figure S12D**, Supporting Information). Compound **2** (but not **3**) also tended to reduce TNF-α (but not IL-1β) exudate levels (**Figure 5D**). Together, as expected from *in vitro* studies, **2** and **3** *in vivo* induce changes in the lipid mediator and cytokine profile that favor the termination and resolution of inflammation, albeit with reduced efficacy and seemingly accompanied by unfavorable adjustments that may limit the anti-inflammatory activity. In this sense, **2** and **3** depleted the peritoneal exudate of anti-inflammatory mediators such as fatty acid epoxides and IL-10 (**Figure 5D,E**).

Despite these mixed changes in the exudate and systemic lipid mediator profile in terms of anti-inflammatory efficacy, both **2** and **3** substantially suppressed immune cell infiltration into the peritoneal cavity during acute inflammation (4 h) (**Figure 5F**). Therefore, we speculated that there might be additional anti-inflammatory mechanisms and extended our investigations into bioactive sphingolipids and endocannabinoids.

Ceramides (Cer), ceramide-1-phosphate (C1P), and sphingosine-1-phosphate (d18:1) (S1P) have pleiotropic functions in the regulation of immunity and inflammation.^[4b, 43b, 45]^ By activating G-protein-coupled S1P receptors, S1P regulates Toll-like receptor signaling as well as immune cell trafficking and maturation, among many other pathways.^[4b, 46]^ Its serum concentration is decreased in zymosan-challenged mice^[43a]^ and negatively correlates with peritonitis in humans.^[47]^ Notably, the role of S1P receptors in inflammation is complex and includes both anti- and pro-inflammatory activities depending on the disease and experimental setting.^[4b, 43b, 48]^ Compounds **2** and **3** have pronounced effects on sphingolipid metabolism and substantially increase plasma S1P levels in mice during the resolution phase (18 h) and, in the case of **2**, also during the acute phase (4 h) (**Figure 5G**). Profiling of intermediates in sphingolipid metabolism implies that **2** and **3** impair S1P degradation rather than enhance S1P biosynthesis, as the levels of the precursor, sphingosine (So), are not elevated in plasma (but interestingly in the exudate) (**Figure S14**, Supporting Information). Our data suggest that **2** and **3** reverse the depletion of S1P in peritonitis, though it remains elusive, whether they increase S1P levels actively or indirectly by alleviating peritonitis. Endocannabinoids are potent immunomodulatory mediators that bind to cannabinoid (CB_1_ and/or CB_2_) and other receptors as extracellular ligands but also trigger intracellular signaling cascades.^[4d]^ Agonists of these receptors have been shown to limit peritoneal inflammation in mice by blocking neutrophil infiltration into the peritoneal cavity^[44c]^ and stimulating LXA_4_ biosynthesis^[44a, 44b]^. Compounds **2** and **3** strongly increase endocannabinoid levels (2-AG > PEA > OEA) during acute inflammation (4 h), specifically in the peritoneal exudate (Figure 5E and **Figure 5H**; **Figure S12E**,**F** and Figure S13, Supporting Information). While endocannabinoid levels are maintained up to 18 h after administration of **2**, they return to baseline in mice treated with **3** and in parallel evoke a trend towards higher plasma concentrations. We conclude that **2** and **3** induce comprehensive changes in the cytokine and lipid mediator profile during the course of resolving inflammation, with favorable adaptations dominating, and the accumulation of endocannabinoids being particularly prominent. Supporting the important role played by endocannabinoids, the number of peritoneal infiltrating cells negatively correlated with local endocannabinoid concentrations in **3** but not **2**-treated mice (**Figure 5I**).

### 2.7 Upregulation of endocannabinoids via inhibition of MAGL and other mechanisms

Endocannabinoids are divided into *N*-acylethanolamines (AEA, PEA and OEA), which are hydrolyzed and thereby inactivated by fatty acid amide hydrolase (FAAH), and monoacylglycerols (1-AG and 2-AG), which are degraded by monoacylglycerol lipase (MAGL).^[4d]^ Compound **3** and, to a lesser extent, **2** decreased cell-free MAGL activity (**Figure 5J**), although the effect was mild and, while possibly contributing to the upregulation of exudate endocannabinoids, cannot readily explain the magnitude of the effect. In addition, neither **2** nor **3** suppressed human recombinant FAAH activity (Figure 5J), suggesting that additional mechanisms, such as lipid dynamics providing phospholipid or diacylglycerol (DG) substrates for endocannabinoid biosynthesis, govern the strong increase in endocannabinoids.

### 2.8 The meroterpenoids 1, 3, 4 and 7 alter the lipidome of innate immune cells

The differential effects of meroterpenoids on the lipid mediator profile led us to speculate that there may be another level of regulation that is under metabolic control and related to the cellular lipid composition. To explore this hypothesis, we exposed PBMCs to meroterpenoids for 48 h and analyzed the cellular lipid composition, focusing on glycerophospholipids, sphingolipids and neutral lipids, in total 202 species. Principal component analysis revealed three clusters based on absolute differences in the lipid composition (**Figure 6A**): Cluster B, consisting of **1, 3**, and **4**, together with compound **7** (cluster C), is clearly separated from Cluster A, which comprises the vehicle control and meroterpenoids (**2, 5, 6, 8, 9**) that have weaker effects on the overall lipidome (**Figure 6B**). Most strikingly, compounds from cluster B induce a shift from neutral lipids, i.e., triacylglycerols (TGs) and/or cholesteryl esters (CE), to phospholipids, specifically (lyso)phosphatidylcholine ((L)PC), phosphatidylserine (PS), cardiolipin (CL), and dihydrosphingomyelin (dhSM) (Figure 6B and **Figure S15** and **16**, Supporting Information. The decrease in neutral lipids (despite forming the inner core of lipid droplets^[49]^) does not seem to be related to impaired lipid droplet biogenesis, as **3** did not reduce the immunofluorescence signal of the coating protein perilipin-2 in A23187-activated PBMCs (**Figure 8B,C** and **Figure S23**, Supporting Information). Absolute amounts of lipid subclasses are shown in **Figure S17** and **Fig. S18**.

**Figure 6.**
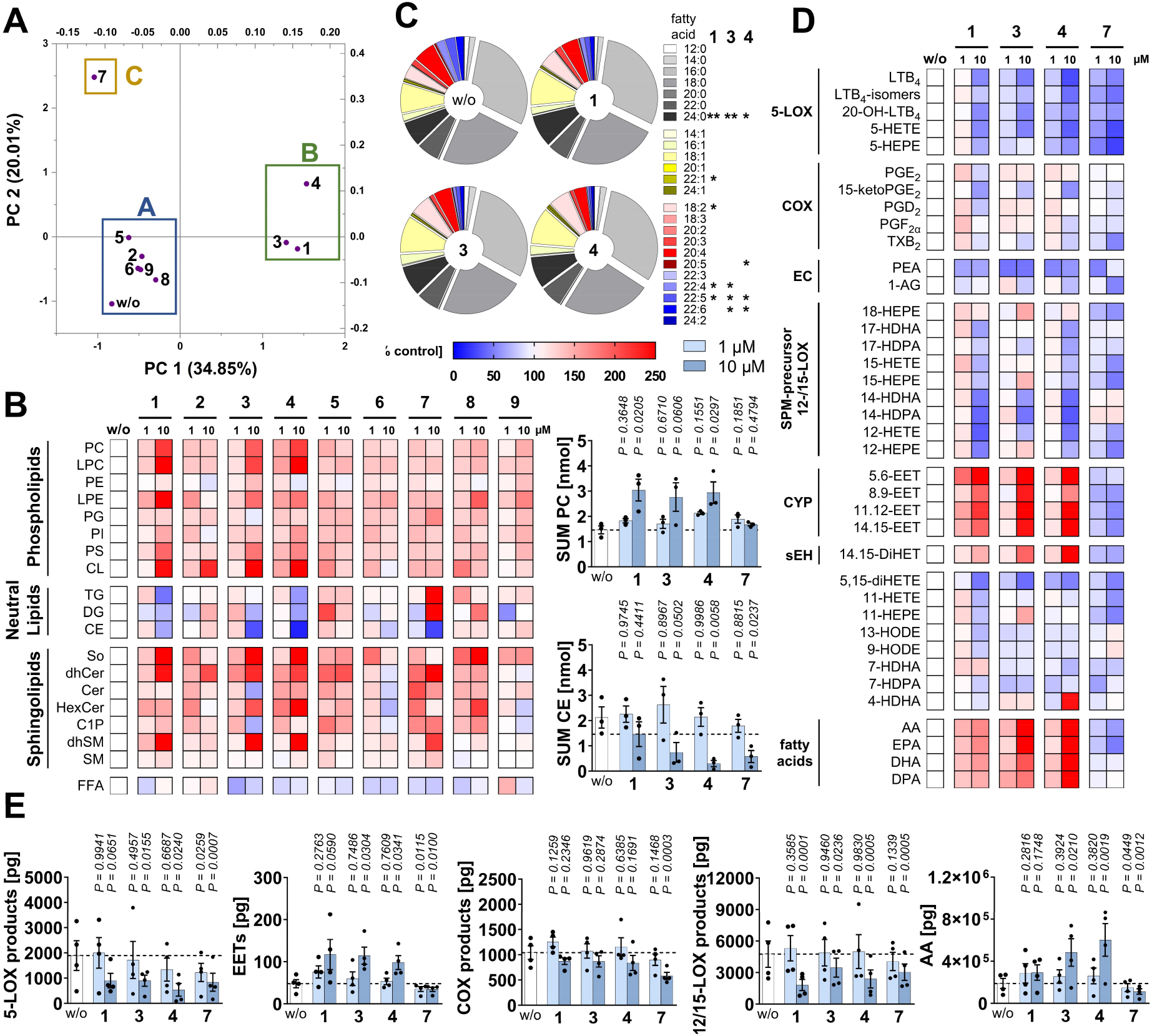
Prolonged exposure of PBMCs to meroterpenoids rearranges the cellular lipidome and establishes an anti-inflammatory lipid mediator profile. A-C) To assess the cellular lipidome, PBMCs were incubated with vehicle (DMSO, 0.1%) or test compounds (10 µM unless otherwise noted) for 48 h. A) Principal component analysis of the mean percentage changes in the amount of lipid species compared to vehicle control. B) Heatmap showing percentage changes in the concentration of individual lipid classes compared to vehicle control and bar charts showing total PC and CE concentrations [per 1×10^6^ PBMCs]. C) Pie charts illustrating changes in the fatty acid composition across all lipid species analyzed (202). Figures S15-16, Supporting Information detail the effects on individual lipid species. Lipid classes analyzed: phosphatidylcholine (PC), lysophosphatidylcholine (LPC), phosphatidylethanolamine (PE), lysophosphatidylethanolamine (LPE), phosphatidylglycerol (PG), phosphatidylinositol (PI), phosphatidylserine (PS), cardiolipin (CL), triacylglycerol (TG), diacylglycerol (DG), cholesteryl ester (CE), sphingosine (So), (dihydro)ceramide ((dh)Cer), hexosylceramide (HexCer), ceramide-1-phosphate (C1P), (dihydro)sphingomyelin ((dh)SM), free fatty acids (FFA). D-E) For lipid mediator analysis, PBMCs were pretreated (48 h) with vehicle (DMSO, 0.1%) or test compounds, washed and activated with calcium ionophore (A23187, 10 min). D) Heatmap showing percentage changes in the concentration of lipid mediators relative to vehicle control. E) Amount of 5-LOX products, EETs, COX products, 12/15-LOX products, and AA (20:4) [per 1×10^6^ cells]. Mean (A-D) or mean + SEM and single data (B,E) from n = 3 (A-C), n = 3-4 (D), n = 4 (E) independent experiments. *P* values given vs. vehicle control (B,C,E) with \**P* < 0.05, \*\**P* < 0.01 (C); two-tailed paired Student *t* test (C) or repeated measures one-way ANOVA of log data (B,E) + Dunnett *post hoc* tests.

In contrast to the compounds from cluster B, **7** suppressed CE, but increased TG levels and had only mild effects on glycerophospholipid levels (Figure 6B; Figure S15-18, Supporting information). The decrease in CE levels appears to be independent of cholesterol *de novo* biosynthesis, because cholesterol levels remained unchanged (Figure S17, Supporting Information). In addition to the changes in the total lipid subclass content, **1, 3, 4**, or **7** extensively altered the relative composition of esterified fatty acids (**Figure 6C; Figure S19-20**, Supporting Information). Significance is reached for docosapentaenoic acid (22:5) and docosahexaenoic acid (22:6) (and other very long-chain fatty acids), which serve as a reservoir for anti-inflammatory/pro-resolving lipid mediator biosynthesis^[3a]^ (including Rvs, PDs, and MaRs) and seem to be released, as suggested by the decrease in levels of the corresponding esterified fatty acids. Taken together, different meroterpenoids have pronounced effects on the abundance and fatty acid composition of major lipid classes in innate immune cells, with small structural changes, such as the methoxylation of **1** to **2**, determining the lipidome-modulating activity.

### 2.9 Long-term treatment of PBMC favors the production of EETs over LTs

To assess the impact of the lipidomic changes induced by **1, 3, 4**, and **7** on the lipid mediator profile, we investigated the capacity of reprogrammed PBMCs to produce lipid mediators. For this purpose, PBMCs were exposed to compounds for 48 h, washed, and then stimulated with the Ca^2+^-ionophore A23187 to induce massive lipid mediator production. As observed for short-term incubations, **1, 3**, and **4** (but not **7**) induced a lipid mediator class switch, however from 5-LOX products to EETs, without increasing the levels of 12-/15-LOX products, COX products, or endocannabinoids, whereas **7** only suppressed the formation of 5-LOX and COX products (**Figure 6D,E** and **Figure S21**, Supporting Information). SPMs were not detected under these stimulatory conditions. While AA (20:4) redirection towards EET biosynthesis cannot be completely excluded for **3** and **4**, compound **1** does not substantially elevate free AA (20:4) levels (Figure 6E).

### 2.10 Correlation between neutral lipid availability and lipid mediator production

Whether changes in the phospholipid or neutral lipid composition correlate with the switch from 5-LOX product to EETs biosynthesis was investigated in PBMCs that were incubated with **1, 3, 4**, or **7** for 48 h (prior to lipid composition analysis) and then treated with A23187 to induce lipid mediator formation in the absence of meroterpenoids. The network in **Figure 7A** shows the co-regulation of lipid species (visualized as nodes) and differentiates between lipid classes by color. The node diameter indicates the relative proportion within each lipid class and the shape indicates the fatty acid composition. Positively correlated lipid species are close together and connected by edges, while counter- or non-correlated lipids are separated from each other.

In **Figure 7B**, the color visualizes correlations (positive: 0.7 ≤ *r* ≤ 1; negative: -1 ≤ *r* ≤ -0.7) between cellular lipid species and the amounts of 5-LOX products or EETs generated. 5-LOX products (decreased levels; Figure 6E) correlate positively with CE (including CE 20:4) and partially with TG species (including TG(18:0_18:0_20:4)), whose levels were decreased by treatment with **1, 3**, or **4** (Figure S16, Supporting Information). The strongest negative correlations for EETs (increased levels; Figure 6E) were observed with TGs as well as phospholipids and Cer that are associated with the TG and CE clusters. Since especially TGs and CE contain substantial amounts of AA (20:4) (**Figure 7C**), it is tempting to speculate that the switch from 5-LOX product to EET formation upon treatment with **1, 3**, or **4** (Figure 6D,E) is at least partially related to lipid subclass-specific AA (20:4) supply. In addition, there are other positive and negative correlations between 5-LOX products or EETs and the cellular lipid composition (Figure 7B). For example, the 5-LOX product formation is negatively correlated with several PI and sphingolipid species, whereas EETs are positively correlated with a compact phospholipid cluster consisting of different PC, PE, PS, and CL species.

**Figure 7.**
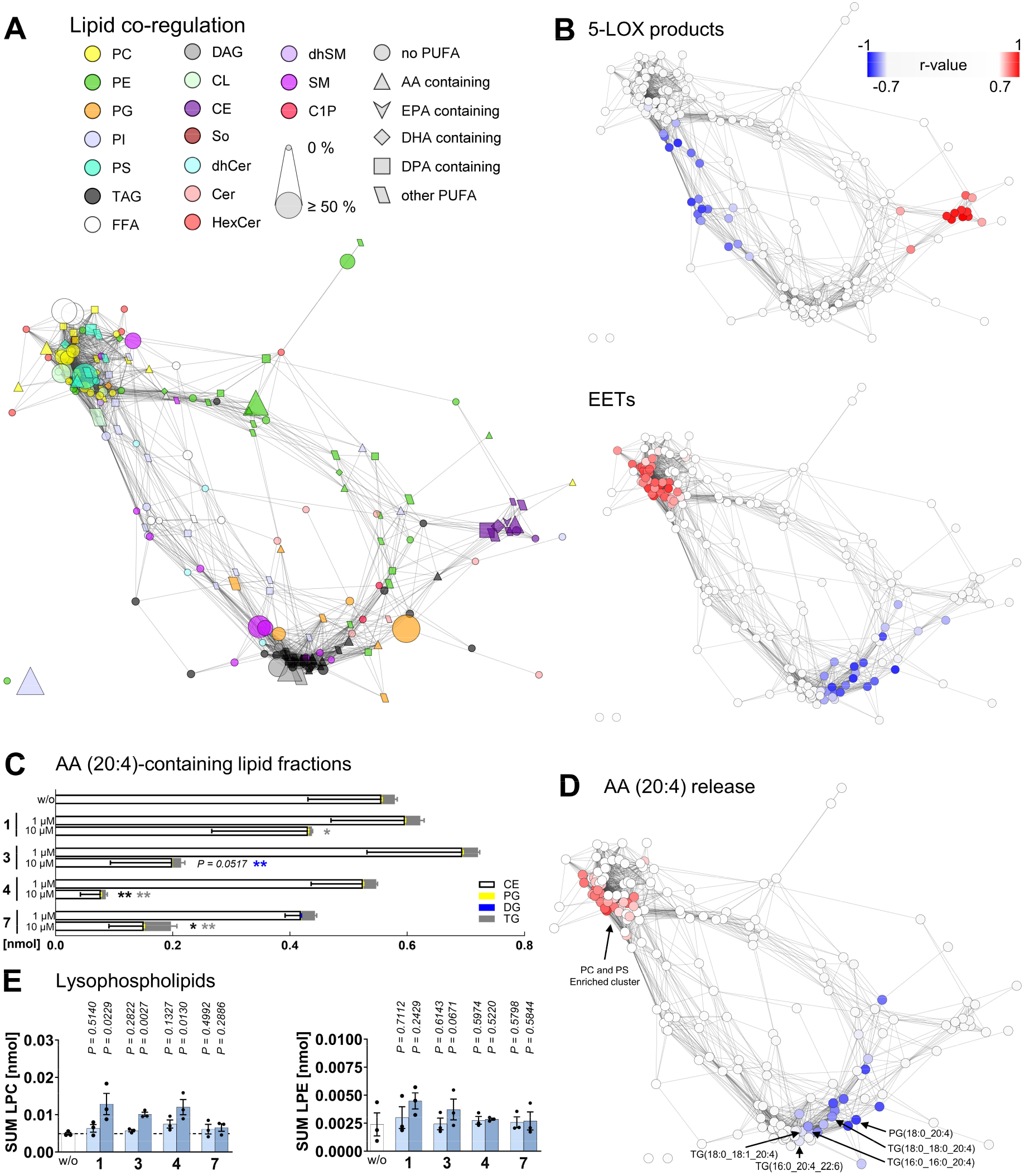
Depletion of neutral lipid fractions in PBMCs is associated with anti-inflammatory changes in lipid mediator production. A) Co-regulated lipid network based on Pearson correlations depicting positive lipid-lipid correlations (r ≥ 0.7). Each node represents a specific lipid (for detailed information on the lipid species, see Figure S15 and S16, Supporting Information) and defines the lipid class and fatty acid composition by color and shape, respectively. The network combines data of PBMCs incubated with **1, 3, 4** or **7** (1 and 10 µM each) for 48 h and was calculated from the mean percentage changes of lipid amounts relative to vehicle from n = 3 independent experiments. B) The network shown in A was overlaid with a color map indicating positive (r ≥ 0.7; red) and negative (r ≤ -0.7; blue) (Pearson) correlations of individual lipid species with lipid mediator subgroups, produced by PBMCs preincubated with **1, 3, 4**, or **7** (1 and 10 µM each) for 48 h after washing and addition of calcium-ionophore (A23187). The lipid mediators belonging to the subclasses are shown in Figure 6D. The correlations were calculated from the mean percentage changes of lipid and lipid mediator quantities relative to vehicle control of n = 3 (lipids) or n = 4 (lipid mediators) independent experiments. C) Amount of AA (20:4)-containing TG, DG, CE, and PG species [per 1 × 10^6^ PBMCs]. D) The network shown in Figure 7A was overlaid with a color map indicating positive (r ≥ 0.7) and negative (r ≤ -0.7) (Pearson) correlations of individual lipid species with AA (20:4). The experimental procedure is identical to B. E) Amount of lysophospholipids in PBMCs incubated with vehicle (DMSO, 0.1%), **1, 3, 4**, or **7** for 48 h [per 1 × 10^6^ cells]. Mean (A,B,D) or mean + SEM (C) and single data (E) from n = 3 (A,C,E), n = 3-4 (B, D) independent experiments. *P* values given vs. vehicle control (C,E) with \**P* < 0.05, \*\**P* < 0.01 (C); repeated measures one-way ANOVA of log data (C,E) + Dunnett *post hoc* tests.

The 5-LOX substrate AA is considered to be provided mainly by phospholipases A_2_, which cleave phospholipids into free fatty acids (FFA) and lysophospholipids^[7]^, but the lipolysis of TGs is also a rich source.^[50]^ Our data indicate that the levels of AA released from activated PBMCs correlate positively with the availability of AA (20:4)-containing PC and PS (**Figure 7D**) and negatively with that of 20:4-containing neutral lipids (TG and CE). Specifically, **3** and **4** and less **1** substantially increased the release of AA from challenged cells (Figure 6E), which *per se* contain more AA (20:4)-containing PC and PS and less AA (20:4)-containing TG and CE when exposed to these meroterpenoids (Figure 6B and Figure S15-18, Supporting Information). These data suggest that PBMCs reorganize AA (20:4) pools from neutral lipids to phospholipids in response to meroterpenoids. As expected from such phospholipid turnover, **1, 3** and **4** resulted in higher intracellular lysophospholipid levels (**Figure 7E**). We speculate that this rearrangement facilitates AA release by phospholipases in activated cells, thereby increasing the total concentration of free AA (20:4), while opposingly affecting AA sub-pools for either 5-LOX or EET biosynthesis. In conclusion, long-term incubation with **1, 3** and **4** induces neutral lipid and phospholipid remodeling that increases the availability of free AA (20:4) and reprograms PBMCs from LT to EET biosynthesis.

### 2.11 Depletion of neutral lipids impairs the capacity for 5-LOX product formation

We investigated whether the decrease in TG and CE levels observed after long-term treatment with **1, 3** and **4** drives the associated lipid mediator class switch from 5-LOX products to EETs in A23187-activated PBMCs. To this end, we suppressed CE biosynthesis with the selective sterol-*O*-acyltransferase (SOAT) inhibitor TMP-153^[51]^ or interfered with TG biosynthesis by inhibiting the diacylglycerol acyltransferase (DGAT)1 and 2 isoenzymes with A-922500^[52]^ and PF-06424439,^[53]^ respectively. Membrane integrity and the number of viable cells were not compromised under these experimental conditions (**Figure S22A**, Supporting Information). As expected, SOAT inhibition reduced cellular CE levels and DGAT-1/2 inhibition lowered the TG content within 48 h and together they decreased both TG and CE levels (**Figure 8A**) as well as the number, but not size, of lipid droplets, as shown by perilipin-2 staining (**Figure 8B**,**C**; **Figure S22B** and Figure S23). Interestingly, only the combined treatment with SOAT and DGAT1/2 inhibitors markedly attenuated 5-LOX product formation in A23187-activated PBMCs (**Figure 8**C,**D**) without obviously changing the subcellular localization of 5-LOX (**Figure S24**, Supporting Information), whereas single treatments had no effect and EET levels were only marginally increased (Figure 8C). Inhibition of both SOAT and DGAT1/2 did not markedly alter AA (20:4) release (**Figure 8E**) and, as observed for **3**, neither substantially affected 5-LOX expression (**Figure 8F**) and subcellular localization (Figure S24, Supporting Information) nor mitogen-activated protein kinase kinase (MEK)1/2 phosphorylation (Figure 8F and **Figure S25A**, Supporting Information), which proceeds 5-LOX nuclear translocation.^[4g, 54]^ However, SOAT together with DGAT1/2 inhibition tended to reduce the overall very low level of FLAP expression, which may contribute to the decrease in 5-LOX product formation (Figure 8H). Additionally, MEK1/2 expression was slightly decreased by dual SOAT and DGAT1/2 inhibition, whereas it was increased by **3**, resulting in elevated cellular p-MEK1/2 levels (Figure 8F and Figure S25A, Supporting Information), possibly as a cellular attempt to counteract the potent 5-LOX inhibition. Our data show that limiting the availability of neutral lipids (TGs and CE), as achieved by dual SOAT and DGAT1/2 inhibition, and observed for **3** and other meroterpenoids, suppresses 5-LOX product formation in innate immune cells. However, also additional mechanisms seem to contribute to the reduced capacity for 5-LOX product formation, given the superior potency of **3** (Figure 6E) compared to SOAT and DGAT1/2 inhibitors (Figure 8C), even when the meroterpenoid was removed prior to cell activation. Impaired FLAP expression can rather be excluded as a driving force since the effect was more pronounced for combined SOAT and DGAT1/2 inhibition than for treatment with **3**.

**Figure 8.**
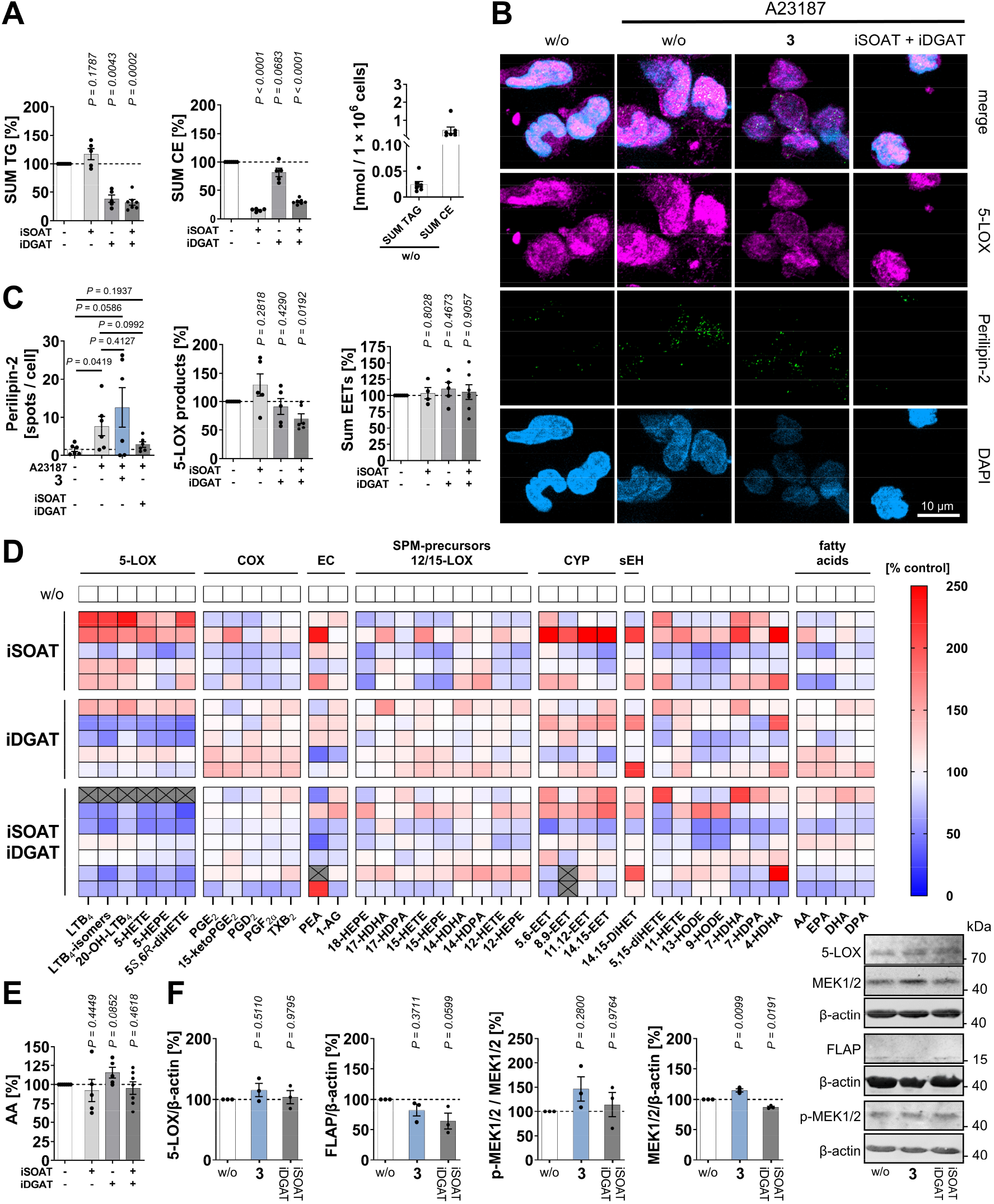
Combined SOAT and DGAT1/2 inhibition decreases neutral lipid levels and lipid droplet number along with suppression of 5-LOX product formation in PBMCs. A,F) For TG and CE analysis and expression studies, cells were treated with vehicle (DMSO, 0.1%), **3** (10 µM), TMP-153 (iSOAT; 500 nM), A-922500 (10 nM) + PF-06424439 (20 nM) (iDGAT), or TMP153 (100 nM) + A-922500 (10 nM) + PF-06424439 (20 nM) (iSOAT + iDGAT) for 48 h. A) Changes in TG and CE contents [per 1×10^6^ PBMCs] and comparison of TG and CE concentrations. B,C) For immunostaining, PBMCs were preincubated with vehicle (DMSO, 0.1%), **3** (10 µM) or iSOAT + iDGAT for 48 h before the medium was replaced and they were treated with calcium ionophore (A23187, 5 min). Immunofluorescence images are stained for perilipin-2 (green, exposure time 35 ms), 5-LOX (violet, exposure time 200 ms), and nuclei (DAPI, light blue, exposure time 10 ms). Images are representative of ≥ 70 single cells analyzed in n = 3 independent experiments. C) Number of perilipin-2-stained spots normalized to cell number. Figure S23, Supplementary Information summarizes zoomed versions of all images used for quantitation. C-E) For lipid mediator analysis, PBMCs were pretreated with vehicle (DMSO, 0.1%), iSOAT, iDGAT or iSOAT + iDGAT for 48 h, washed and activated with calcium ionophore (A23187, 10 min). D) Heatmap showing percentage changes in the concentration of lipid mediators relative to vehicle control for each independent experiment, with the crossed sections indicating significant outliers. E) AA (20:4) concentration. F) Effect of **3** and iSOAT + iDGAT on the expression of 5-LOX, FLAP and MEK1/2 and on MEK1/2 phosphorylation. Western blots are representative of n = 3 independent experiments. Uncropped blots are shown in Figure S25B, Supporting Information. Mean (D) or mean + SEM and single data (A,C,E,F) from n = 3 (B,C perilipin-2,F), n = 4-7 (A,C,D,E) independent experiments. *P* values given vs. vehicle control (A,C,E,F) or as indicated by bars (C); repeated measures one-way ANOVA + Dunnett *post hoc* tests (F), two-tailed paired Student *t* test of log data (A,C,E) or two-tailed unpaired Student *t* test (C perilipin-2).

## 3 Discussion

The epilipidome comprises structurally and functionally diverse lipids, including oxidized species, whose physiological functions on a global scale we are only beginning to understand. ^[55]^ Lipid mediators such as prostanoids, LTs, or EETs have been recognized early on to play a fundamental role in inflammatory diseases but also beyond,^[2b, 4e, 4h, 4i]^ and great efforts have been made to develop anti-inflammatory drug candidates against key enzymes in their biosynthesis or degradation, such as COX, 5-LOX, FLAP, microsomal prostaglandin E_2_ synthase, or soluble epoxide hydrolase,^[2b, 16b, 21c, 56]^ and also polypharmacological ligands have been successfully designed.^[4i, 21a, 21b, 24]^ With the ever-increasing number of bioactive lipids and the emerging functional links to cell metabolism and lipid pools that were previously considered purely structural, drug development faces significant challenges. Next-generation strategies will require input from OMICs technologies and network science to define favorable lipid signatures and to discover small molecules for targeted manipulation of local and systemic lipid profiles under disease conditions. The latter is of high importance given that the lipid network is dynamically and kinetically fluctuating in response to intrinsic and extrinsic factors, including the metabolic status of the cell and the organism. ^[57]^ It is now generally accepted that dysregulated cellular metabolism is a major driver of inflammation and that a better understanding of this under-explored interplay may provide access to more potent and safer anti-inflammatory drugs.^[58]^ Based on these considerations, we initiated a drug discovery program aimed at identifying novel biogenic scaffolds, which allow to favorably manipulate pre-defined lipid signatures in human innate immune cells. Our initial focus was placed on small molecules that suppress the biosynthesis of pro-inflammatory lipid mediators, while increasing the availability of anti-inflammatory or pro-resolving mediators. This approach was inspired from the current understanding of inflammation as a biphasic process of initiation and active resolution,^[1]^ the latter being essentially regulated by lipid mediators with pro-resolving activity.^[2d]^ While there is still some debate as to whether tri-hydroxylated SPMs in particular are produced in sufficient quantities to elicit a physiological response,^[15]^ there is also strong evidence from animal studies that excess SPMs (e.g., upon injection) are highly effective in alleviating inflammation,^[2d, 13]^ and initial clinical trials on SPM analogs^[59]^ or SPM precursors^[60]^ support anti-inflammatory and/or analgesic activity. As a result, the challenge for small molecule inflammation-resolving drugs is not only to increase SPM levels, but to increase them sufficiently to produce a pronounced and long-lasting effect. Following this newly established routine, we identified the meroterpenoid **1** as a potential candidate that potently suppresses 5-LOX product formation while increasing SPM biosynthesis. Based on this initial HIT, we first investigated whether structural modifications introduced by total synthesis^[25]^ could enhance this lipid mediator class switch. This strategy succeeded in identifying structural elements that allowed derivatives to outperform the parental compound and even increase SPM production in resting (unchallenged) cells, superior to previously described (non-toxic) small molecules that upregulate SPM biosynthesis.^[22a, 22f] [22b, 23a]^ Note that we compare fold-increases rather than absolute SPM levels between different studies, as metabololipidomics analysis still lacks sufficient inter-laboratory standardization,^[61]^ resulting in absolute values varying by more than a decade, even under comparable experimental settings.^[62]^ Second, we extended our investigations of the lipid mediator network to additional anti-inflammatory and immunomodulatory lipid classes, such as EETs, endocannabinoids, and bioactive sphingolipids, and found that they are also substantially upregulated, although differently and partially opposite between experimental systems. For example, compound **3** markedly increased EET levels in SACM-challenged human macrophages, but decreased them in peritoneal exudates of zymosan-injected mice.

Meroterpenoids have an overall anti-inflammatory activity, as shown in a mouse model of self-resolving inflammation, but it is difficult to estimate the contribution of individual lipid changes, and some adjustments may even be detrimental under certain disease conditions. While it is generally accepted that LTs promote,^[2b, 4g, 4i, 9]^ EETs suppress,^[63]^ and SPMs resolve inflammation,^[13]^ the directionality of the effect is less clear for prostaglandin E_2_,^[4i, 8]^ endocannabinoids,^[4d]^ and S1P.^[17a]^ All three play a central role in immune regulation and have well-documented anti-inflammatory functions, but can also promote the inflammatory reaction depending on the context.^[4b, 4d, 4i, 8, 17a, 64]^ This complex regulation and functionality of lipid mediator networks reflects the complex nature of inflammation and underscores the need to differentiate between inflammatory diseases and their specific treatment.

It is remarkable that small structural changes in meroterpenoids allow to tune the lipid profiles generated by PBMCs and macrophage subsets. Especially in the context of SPMs, where the (patho)physiological functions and signaling routes of individual subclasses are still obscure, ^[2a, 13, 15]^ such tools could be of high value for directional manipulation of endogenous SPM production. Specifically, meroterpenoids can shift SPM production in activated M2 macrophages to either PDs (**5**), PDs and MaRs (**2**), or to all major SPM classes (**3**). These preferences in shaping the lipid mediator profile do not readily correlate with differences in the molecular mechanism, as far as we have been able to uncover, and we speculate that they may determine preferential sites of subcellular accumulation or engage different kinetics. In fact, SPM production is a tightly coordinated, still incompletely understood process involving lipoxygenases and other enzymes that are localized in different subcellular compartments, some of which must undergo translocation for efficient SPM biosynthesis, at least under physiological conditions.^[4g, 12]^

Unraveling the molecular mechanisms of such extensive lipid alterations is hampered by the multitude of possible targets and the incompletely understood physiological regulation of lipid mediator networks.^[2b, 4e-g, 12, 15, 63]^ At this point, we can explain the suppression of leukotriene biosynthesis, interestingly achieved either by direct 5-LOX inhibition or by interference with FLAP, and gained a basic understanding of the molecular requirements for the induction of SPM biosynthesis, which seems to involve a moderate release of PUFAs from various sources (which boosts SPM biosynthesis but is otherwise optional) and a facilitated translocation of 15-LOX-1 to particulate compartments in the cytoplasm. The latter has been observed before and associated with efficient SPM biosynthesis,^[11-12, 29]^ but never been defined in terms of composition or function. Here, we show that meroterpenoids mobilize PUFAs from different lipid subclasses and favorably redirect AA (20:4) from neutral lipids to specific phospholipids over a longer time-scale. Less well understood are the subcellular compartments from which these PUFAs are derived and whether their release from specific lipid classes is functionally coupled to the biosynthesis of specific lipid mediators.

First hints in this direction came from network correlation analysis, where we generated a co-regulated lipid network based on data from PBMCs treated long-term with meroterpenoids, specifically with derivatives showing pronounced effects on the cellular lipid composition. Correlation of this network with the capacity of cells to produce lipid mediators, notably after removal of the meroterpenoids to exclude direct effects on lipid biosynthetic enzymes, revealed unexpected associations between LT respectively EET levels and major neutral and membrane lipid classes. Unexpected because this correlation was not limited to PUFA-containing species, but also included TGs, CEs, and phospholipids carrying saturated or monounsaturated fatty acids. We confirmed the functional relevance between TG and CE depletion and impaired 5-LOX product formation by selectively inhibiting SOAT to suppress CE biosynthesis and DGAT1 and DGAT2 to suppress TG biosynthesis. Inhibition of all three acyltransferases is required to downregulate the capacity for long-term 5-LOX product formation in PBMC. These results are of potential clinical relevance, since inhibitors of SOAT and DGAT produced mixed results in phase II and III clinical trials and SOAT inhibitors have entered the market in Japan and other countries for the treatment of hypercholesterolemia and atherosclerosis^[65]^, comprising an inflammatory component^[66]^ that has been associated with LT biosynthesis before.^[66a]^ Our findings are consistent with previous reports that manipulated cellular TG and CE levels and found a decrease in LT production, either by inhibiting DGAT1^[67]^ or by reducing PUFA release from lipid droplets by targeting adipose triglyceride lipase (ATGL)^[68]^ or lysosomal acid lipase (LAL).^[69]^. These enzymes, together with hormone-sensitive lipase, are all expressed in human PBMCs^[70]^ and release PUFAs from neutral lipids and phospholipids contained in lipid droplets,^[50]^ which have been identified as sites of LT formation.^[71]^ Lipid droplets are dynamic lipid storage organelles that are connected to most other cellular organelles through membrane contact sites and are at the center of lipid and energy homeostasis.^[49]^ Interestingly, lipid droplets host key enzymes in lipid mediator biosynthesis^[50, 71b]^, including 5-LOX^[50]^, and are depleted upon treatment with SOAT and DGAT inhibitors under our experimental conditions. Therefore, it is tempting to speculate that the depletion of neutral lipids in lipid droplets either limits the substrate access to 5-LOX (and possibly to specific other isoenzymes in lipid mediator biosynthesis) or interferes with the subcellular distribution of 5-LOX, which in turn could affect the functional coupling with FLAP or other lipoxygenases, thereby reducing 5-LOX product formation.

## 4 Conclusion

Low-grade inflammation is a central driver of chronic inflammatory and metabolic diseases and needs to be tackled with efficient and safe therapeutic approaches. OMICs technologies and network science are opening new opportunities to discover small molecules that reset aberrant metabolism on a global scale. By applying functional metabololipidomics and focusing on lipid mediator class switches in innate immune cells that either suppress or resolve inflammation and contribute to immunoregulation, we have discovered biogenic meroterpenoid scaffolds that favorably manipulate the lipid mediator profile by multiple mechanisms both in innate immune cells *in vitro* and murine peritonitis *in vivo*. The mode of action ranges from the direct inhibition of pro-inflammatory factors in lipid mediator metabolism (5-LOX, FLAP, MAGL, p-IκBα), to control of subcellular localization (15-LOX-1) and substrate supply (PUFAs), to reprogramming of cellular lipid composition for sustained regulation of the lipid mediator-biosynthetic capacity. Our study illustrates the power of lipidomics-guided drug development for the discovery of anti-inflammatory small molecule leads and advances our understanding of the emerging link between lipid diversity, cell metabolism, and low-grade inflammation.

## 5 Experimental Section

### Test compounds, lipids and standards

Compounds **1** to **9** were synthesized as described^[25]^, with the purity being determined by 1H-NMR to be ≥ 95%. Test compounds and small molecule inhibitors were dissolved in DMSO and stored in the dark at -20 °C (test compound dilutions and inhibitors) or -80 °C (test compound stock solutions) under argon, minimizing freeze-thaw cycles. Lipid mediators, fatty acids, and standards were obtained from Cayman Chemicals (Ann Arbor, MI) and stored at -80 °C upon receipt. Lipids were diluted in methanol/water (1:1) (fatty acids, oxylipins, and endocannabinoids) or methanol (phospholipids and neutral lipids), and aliquots were stored in the dark at -80 °C under argon.

### Analysis of cell numbers, viability, and mitochondrial dehydrogenase activity

Membrane integrity, total and viable cell counts were determined using a Vi-CELL Series Cell Counter (Beckman Coulter, Krefeld, Germany; software: Vi-Cell XR Cell Viability Analyzer, version 2.06.3) after trypan blue staining.^[6]^

Cellular dehydrogenase activity was analyzed by the conversion of 3-(4,5-dimethylthiazol-2-yl)-2,5-diphenyltetrazolium bromide (MTT) (Sigma Aldrich, St. Louis, MO) as described.^[22e, 72]^ Briefly, human PBMCs (2 × 10^5^/well of a 96-well plate) were treated with vehicle (DMSO, 0.5%), test compounds, or staurosporine (1 µM, pan-kinase inhibitor, Sigma Aldrich) and incubated for 24 h or 48 h (37 °C; 5% CO_2_) in RPMI 1640 medium (100 µL) supplemented with 5% FCS, 2 mmol/L *L*-glutamine, 100 U/mL penicillin, and 100 µg/mL streptomycin. MTT (5 mg/mL in PBS pH 7.4; 20 µL) was added and the incubation continued for another 3 h (37 °C; 5% CO_2_) until blue formazan crystals were clearly visible. After solubilization of the formazan with SDS lysis buffer (10% SDS in 20 mM HCl; 100 µL) (> 16 h) under shacking, the absorbance was measured at 570 nm using a SpectraMax iD3 Microplate Reader (Molecular Devices, San José, CA).

### Isolation of PBMCs from human blood products

The Central Institute for Blood Transfusion and Immunology of the University Hospital Innsbruck, Tirol Kliniken GmbH (Austria) provided leukoreduction system chambers (LRSCs) derived from platelet donors (18-65 years) after informed consent that residual blood can be used for scientific purposes. Platelet donors underwent a physical examination by trained medical personnel and fulfilled the criteria of the Austrian Blood Donation Regulation (BGBI. II Nr. 217/2022). Human PBMCs were freshly isolated from LRSCs, as previously described.^[22d, 73]^ Briefly, PBMCs were obtained by isopycnic density gradient centrifugation using Histopaque®-1077 (Sigma-Aldrich, St. Louis, MO, USA) (400 × g, 20 min, room temperature (RT)) and purified by hypotonic lysis of erythrocytes (water) and two consecutive washing steps with PBS pH 7.4 (270 × g, 5 min, RT).

### Monocyte differentiation and polarization into M1- and M2-type macrophages

Freshly isolated PBMCs consisting predominantly of CD14^+^/CD16^+^ and CD14^+^/CD16^-^ monocytes (46.4 ± 4.9% as determined by flow cytometric analysis), were stimulated with 20 ng/mL GM-CSF (HiSS Diagnostics GmbH, Freiburg, Germany) or M-CSF (HiSS Diagnostics GmbH) for 6 days in macrophage medium (RPMI 1640 supplemented with 10% FCS, 2 mmol/L *L*-glutamine, 100 U/mL penicillin, and 100 µg/mL streptomycin) to obtain monocyte-derived M_0_ macrophages.

Polarization of M_0_ macrophages for 48 h in macrophage medium supplemented with 100 ng/mL lipopolysaccharide (LPS; *Escherichia coli* O127:B8, Sigma Aldrich) and 20 ng/mL interferon-γ (Peprotech, Hamburg, Germany) yielded M1-type macrophages, whereas 20 ng/mL interleukin-4 (Peprotech, Hamburg, Germany) was added to generate M2-type macrophages.^[22d, 22e]^

### Treatment of human innate immune cells for lipid mediator analysis

Human PBMCs (5 × 10^6^ cells/mL in 1 mL PBS pH 7.4 plus 1 mM CaCl_2_) were pretreated with vehicle (DMSO; 0.1%) or test compounds for 10 min (37 °C) before cells were stimulated by addition of the Ca^2+^-ionophore A23187 (Cayman Chemical; 2.5 µM, 10 min) alone or in combination with AA (20:4) (Cayman Chemical; 20 µM). Monocyte-derived M1 and M2 macrophages (2 mL PBS pH 7.4 plus 1 mM CaCl_2_) were challenged with SACM (1.0%, 3 h) after 15 min preincubation or 48 h treatment during polarization with vehicle (DMSO; 0.1%) or test compounds (37 °C and 5% CO_2_). SACM was obtained by culturing *Staphylococcus aureus* (LS1 strain) in Brain Heart Infusion (BHI) medium for 18 h followed by sterile filtration of the supernatant (3,400 × g, 10 min, RT) through a Rotilabo^®^-syringe filter (PVDF, 0.22 µm, Roth, Karlsruhe, Germany).^[73-74]^ Alternatively, M2 cells were incubated with vehicle (DMSO, 0.1%) or test compounds for 195 min without further stimulation to investigate lipid mediator profiles in resting macrophages.

To investigate long-term effects on lipid mediator profiles, freshly isolated human PBMCs (1 × 10^7^ cells/mL in 4 mL RPMI 1640 medium supplemented with 5% FCS, 2 mmol/L *L*-glutamine, 100 U/mL penicillin and 100 µg/mL streptomycin) were incubated with vehicle (DMSO, 0.1%), test compounds, the SOAT inhibitor TMP-153 (500 nM), the DGAT1 inhibitor A-922500 (10 nM) plus the DGAT2 inhibitor PF-06424439 (20 nM), or TMP153 (100 nM) plus A-922500 (10 nM) and PF-06424439 (20 nM) for 48 h. Cells were harvested by sequential washing with PBS pH 7.4 containing 5 mM EDTA (ice-cold; 1 mL) and PBS pH 7.4 (ice-cold; 1mL). The number of viable cells was determined using a Vi-CELL Series Cell Counter as described above. The cells were then pelleted (270 × g, 7 min, 4 °C), washed with PBS pH 7.4 (ice-cold; 1 mL), resuspended in PBS pH 7.4 plus 1 mM CaCl_2_ (ice-cold; 1 mL) and stimulated with the Ca^2+^-ionophore A23187 (Cayman Chemical; 2.5 µM, 10 min) at 37 °C.

Lipid mediator formation was stopped with ice-cold methanol (2 mL for PBMCs and 3.5 mL for macrophages) containing deuterium-labeled internal standards (Cayman Chemical, Ann Arbor, MI), i.e., 200 pg 5*S*-hydroxy-6*E*,8*Z*,11*Z*,14*Z*-eicosatetraenoic-5,6,8,9,11,12,14,15-d8 acid, 5*S*,12*R*-dihydroxy-6*Z*,8*E*,10*E*,14*Z*-eicosatetraenoic-6,7,14,15-d4 acid, 9-oxo-11α,15*S*-dihydroxy-prosta-5*Z*,13*E*-dien-1-oic-3,3,4,4-d4 acid, 5*S*,6*R*,15*S*-trihydroxyicosa-7*E*,9*E*,11*Z*,13*E*-tetraenoic-19,19,20,20,20-d5 acid, and 7*S*,16*R*,17*S*-trihydroxy-4*Z*,8*E*,10*Z*,12*E*,14*E*,19*Z*-21,21’,22,22,22-d5 docosahexaenoic acid, and 2000 pg 5*Z*,8*Z*,11*Z*,14*Z*-eicosatetraenoic-5,6,8,9,11,12,14,15-d8 acid. For Figure 5B-C,E,H,I, Figure 6D-E, Figure 7B,D and Figure 8C-E; Figure S12B-F, Figure S13 and Figure S21, Supporting information additionally 200 pg (±)8(9)-epoxy-5*Z*,8*Z*,14*Z*-eicosatrienoic-16,16,17,17,18,18,19,19,20,20,20-d11 acid and for Figure 8C-E additionally 2000 pg *N*-(2-hydroxyethyl)-5*Z*,8*Z*,11*Z*,14*Z*-eicosatetraenamide-5,6,8,9,11,12,14,15-d8 were added.

### Determination of cell-free 15-lipoxygenase activity

M2 macrophages (2 × 10^6^ cells) were resuspended in PBS pH 7.4 plus 1 mM EDTA and homogenized (3-times 15 s; power: 35% of 125 W; Q125 Sonicator, QSonica, Newtown, CT). Complete homogenization was visually confirmed by light microscopy. Following 15 min preincubation with vehicle (DMSO, 0.1%) or test compounds (on ice), the reaction was initiated by adding 2 mM CaCl_2_ and 20 µM AA (20:4). The reaction was terminated after 15 min at 37 °C by addition of 1 mL ice-cold methanol containing 200 pg 5*S*-hydroxy-6E,8Z,11Z,14Z-eicosatetraenoic-5,6,8,9,11,12,14,15-d8 acid as internal standard.

### Solid-phase extraction and UPLC-MS/MS-based profiling of lipid mediators

To allow for protein precipitation, samples were stored at -20 °C for at least 1 h prior to centrifugation (750 × g, 10 min, 4 °C). Samples (PBMCs: 3 mL; M1/M2 macrophages: 5.5 mL; exudate: 2.8 mL; plasma: 2.4 mL) were premixed with acidified water (PBMCs, exudate and plasma: 8 mL; M1/M2 macrophages: 9 mL; pH 3.5) and loaded onto conditioned (6 mL methanol) and equilibrated (2 mL water) solid phase extraction cartridges (Sep-Pak® Vac 6cc 500 mg/6 mL C-18; Waters, Milford, MA). The cartridges were then washed with water (6 mL) and hexane (6 mL, 4 °C) before lipid mediators were eluted with methyl formate (6 mL). After evaporation of the organic solvent (TurboVap LV; Biotage, Uppsala, Sweden), the lipid mediators were redissolved in methanol/water (1:1), centrifuged (750 × g, 10 min, 4 °C and twice 21,100 × g, 4 °C, 5 min), and subjected to UPLC-MS/MS analysis.^[22d,73]^

The chromatographic separation of lipid mediators was performed on an Acquity UPLC BEH C18 column (130Å, 1.7 µm, 2.1 × 100 mm, Waters) at 55 °C column temperature using an ExionLC AD UHPLC system (Sciex, Framingham, MA), with mobile phase A (methanol, 0.01% acetic acid) and mobile phase B (water/methanol, 90/10, 0.01% acetic acid) at a flow rate of 0.35 mL/min. For the analysis of oxylipins, the gradient was ramped at 0.35 mL/min from 35.6% to 84.4% A within 12.5 min followed by 5 min of isocratic elution (97.8% A) (Figure 1 and 2; Figure S1E-H and Figure S2-6, Supporting Information).^[22d]^ Alternatively, oxylipins and endocannabinoids were separated using a gradient that increased linearly from 35.6% to 84.4% A within 12.5 min and then to 87.0% A within 2.5 min followed by 3 min of isocratic elution at 97.8% A.^[73]^

Oxylipins and free PUFAs were analyzed in negative and endocannabinoids in positive ion mode by scheduled multiple reaction monitoring (MRM) and polarity switching using a QTRAP 6500^+^ Mass Spectrometer (Sciex) equipped with an IonDrive Turbo V Ion Source and a TurboIonSpray probe for electrospray ionization (Sciex). The curtain, sheath and auxiliary gas pressures were set to 40 psi, the collision gas to medium, the heated capillary temperature to 500 °C and the ion spray voltage to - 4000 V and 4000 V, respectively. The MRM transitions used for quantitation and their corresponding parameters are listed in **Table 2**: retention time (rt), declustering potential (DP), entrance potential (EP), collision energy (CE), collision cell exit potential (CXP), and MRM detection window. Lipid concentrations refer to an 11- or 15-point standard curve and were normalized to a subclass-specific deuterated internal standard and to cell number or sample volume. Mass spectra were acquired and processed using Analyst 1.7.1 or 1.7.2 and 1.6.3 (Sciex).

**Table 2.**
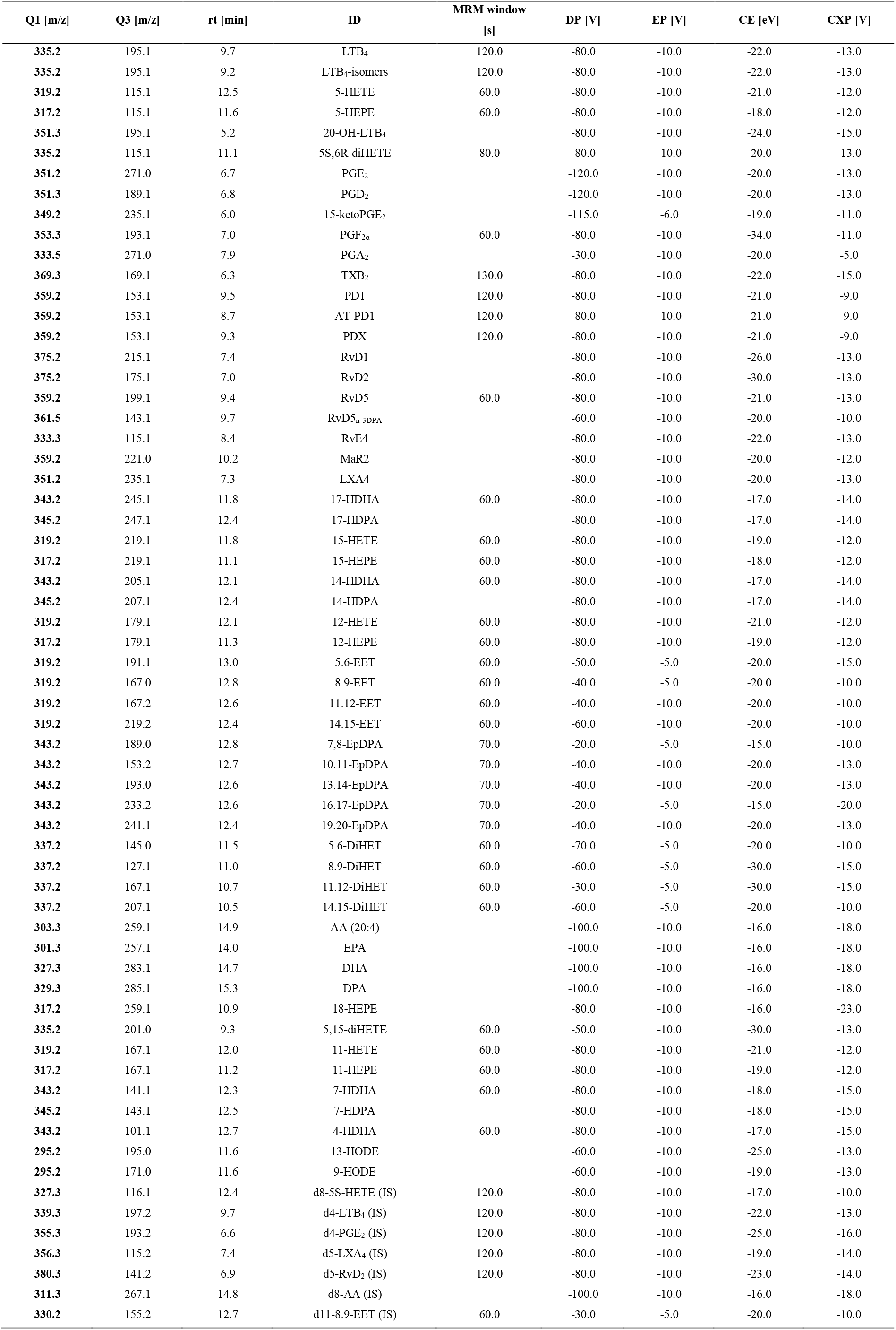
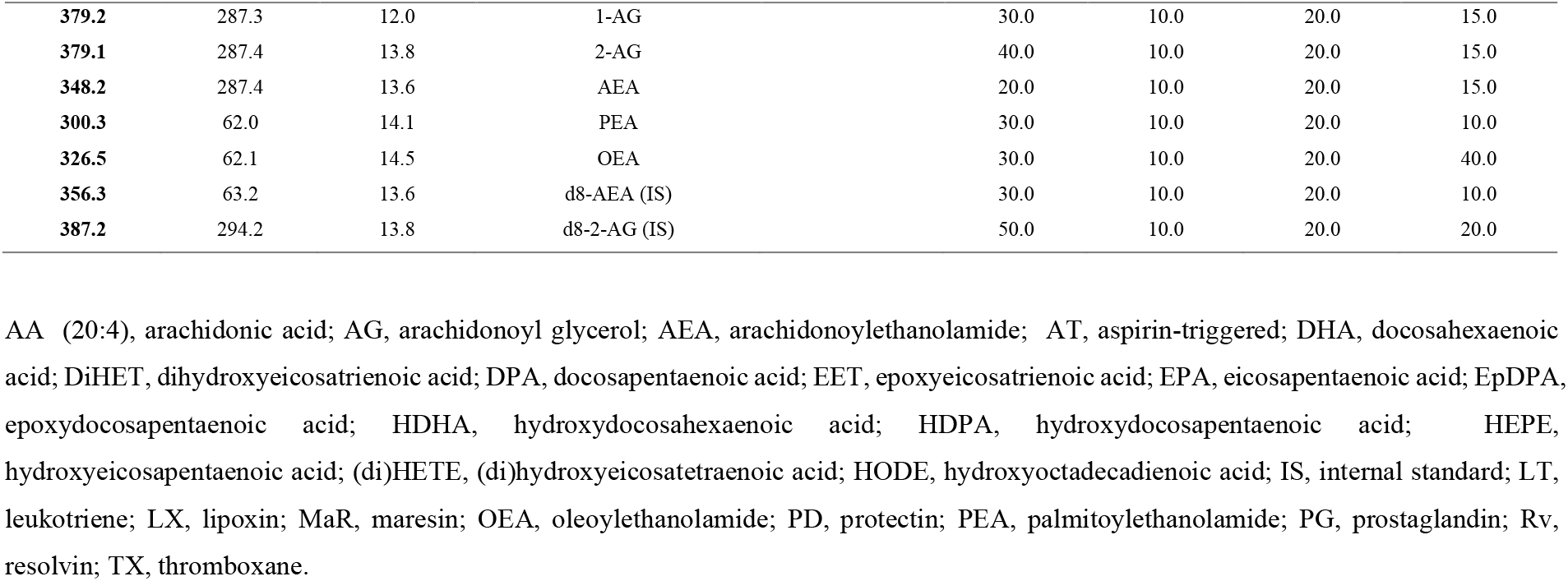
Quantitative MRM transitions of lipid mediators and endocannabinoids.

External standards (Cayman Chemical): 5*S*,12*R*-dihydroxy-6*Z*,8*E*,10*E*,14Z-eicosatetraenoic acid; (±)5-hydroxy-6*E*,8*Z*,11*Z*,14*Z*-eicosatetraenoic acid, (±)-5-hydroxy-6*E*,8*Z*,11*Z*,14*Z*,17*Z*-eicosapentaenoic acid; 5*S*,12*R*,20-trihydroxy-6*Z*,8*E*,10*E*,14*Z*-eicosatetraenoic acid; 5*S*,6*R*-dihydroxy-7*E*,9*E*,11*Z*,14*Z*-eicosatetraenoic acid; 9α,15*S*-dihydroxy-11-oxo-prosta-5*Z*,13*E*-dien-1-oic acid; 9-oxo-11α,15*S*-dihydroxy-prosta-5*Z*,13*E*-dien-1-oic acid; 9,15-dioxo-11α-hydroxy-prosta-5*Z*,13*E*-dien-1-oic acid; 9α,11α,15*S*-trihydroxy-prosta-5*Z*,13*E*-dien-1-oic acid; 9-oxo-15*S*-hydroxy-prosta-5*Z*,10,13*E*-trien-1-oic acid; 9α,11,15*S*-trihydroxythromba-5*Z*,13*E*-dien-1-oic acid; 10*R*,17*S*-dihydroxy-4*Z*,7*Z*,11*E*,13*E*,15*Z*,19*Z*-docosahexaenoic acid; 10*R*,17*R*-dihydroxy-4*Z*,7*Z*,11*E*,13*E*,15*Z*,19*Z*-docosahexaenoic acid; 10(*S*),17(*S*)-dihydroxy-4*Z*,7*Z*,11*E*,13*Z*,15*E*,19*Z*-docosahexaenoic acid; 7*S*,8*R*,17*S*-trihydroxy-4*Z*,9*E*,11*E*,13*Z*,15*E*,19*Z*-docosahexaenoic acid; 7*S*,16*R*,17*S*-trihydroxy-4*Z*,8*E*,10*Z*,12*E*,14*E*,19*Z*-docosahexaenoic acid; 7*S*,17*S*-dihydroxy-4*Z*,8*E*,10*Z*,13*Z*,15*E*,19*Z*-docosahexaenoic acid; 7,17-dihydroxy-8*E*,10*Z*,13*Z*,15*E*,19*Z*-docosapentaenoic acid; 5*S*,15*S*-dihydroxy-6*E*,8*Z*,11*Z*,13*E*,17*Z*-eicosapentaenoic acid; 13*R*,14*S*-dihydroxy-4*Z*,7*Z*,9*E*,11*E*,16*Z*,19*Z*-docosahexaenoic acid; 5*S*,6*R*,15*S*-trihydroxy-7*E*,9*E*,11*Z*,13*E*-eicosatetraenoic acid; (±)17-hydroxy-4*Z*,7*Z*,10*Z*,13*Z*,15*E*,19*Z*-docosahexaenoic acid; 14*S*-hydroxy-4*Z*,7*Z*,10*Z*,12*E*,16*Z*,19*Z*-docosahexaenoic acid; (±)7-hydroxy-4*Z*,8*E*,10*Z*,13*Z*,16*Z*,19*Z*-docosahexaenoic acid; (±)4-hydroxy-5*E*,7*Z*,10*Z*,13*Z*,16*Z*,19*Z*-docosahexaenoic acid; (±)-18-hydroxy-5*Z*,8*Z*,11*Z*,14*Z*,16*E*-eicosapentaenoic acid; 15*S*-hydroxy-5*Z*,8*Z*,11*Z*,13*E*,17*Z*-eicosapentaenoic acid; 12*S*-hydroxy-5*Z*,8*Z*,10*E*,14*Z*,17*Z*-eicosapentaenoic acid; (±)-11-hydroxy-5*Z*,8*Z*,12*E*,14*Z*,17*Z*-eicosapentaenoic acid; 15*S*-hydroxy-5*Z*,8*Z*,11*Z*,13*E*-eicosatetraenoic acid; 12*S*-hydroxy-5*Z*,8*Z*,10*E*,14*Z*-eicosatetraenoic acid; 11*S*-hydroxy-5*Z*,8*Z*,12*E*,14*Z*-eicosatetraenoic acid; 5*S*,15*S*-dihydroxy-6*E*,8*Z*,10*Z*,13*E*-eicosatetraenoic acid; (±)-9-hydroxy-10*E*,12*Z*-octadecadienoic acid; (±)-13-hydroxy-9*Z*,11*E*-octadecadienoic acid; (±)5,6-epoxy-8*Z*,11*Z*,14*Z*-eicosatrienoic acid; (±)8,9-epoxy-5*Z*,11*Z*,14*Z*-eicosatrienoic acid; (±)11,(12)-epoxy-5*Z*,8*Z*,14*Z*-eicosatrienoic acid; (±)14(15)-epoxy-5*Z*,8*Z*,11*Z*-eicosatrienoic acid; (±)-(4*Z*)-6-[3-(2*Z*,5*Z*,8*Z*,11*Z*)-2,5,8,11-tetradecatetraen-1-yl-2-oxiranyl]-4-hexenoic acid; (±)-(4*Z*,7*Z*)-9-[3-(2*Z*,5*Z*,8*Z*)-2,5,8-undecatrien-1-yl-2-oxiranyl]-4,7-nonadienoic acid; (±)-(4*Z*,7*Z*,10*Z*)-12-[3-(2Z,5Z)-2,5-octadien-1-yl-2-oxiranyl]-4,7,10-dodecatrienoic acid; (±)16,17-epoxy-4*Z*,7*Z*,10*Z*,13*Z*,19*Z*-docosapentaenoic acid; (±)19,20-epoxy-4*Z*,7*Z*,10*Z*,13*Z*,16*Z*-docosapentaenoic acid; (±)5,6-dihydroxy-8*Z*,11*Z*,14*Z*-eicosatrienoic acid; (±)8,9-dihydroxy-5*Z*,11*Z*,14*Z*-eicosatrienoic acid;(±)11,12-dihydroxy-5*Z*,8*Z*,14*Z*-eicosatrienoic acid; (±)14,15-dihydroxy-5*Z*,8*Z*,11*Z*-eicosatrienoic acid; 5*Z*,8*Z*,11*Z*,14*Z*-eicosatetraenoic acid; 5*Z*,8*Z*,11*Z*,14*Z*,17*Z*-eicosapentaenoic acid, 4*Z*,7*Z*,10*Z*,13*Z*,16*Z*,19*Z*-docosahexaenoic acid; 7*Z*,10*Z*,13*Z*,16*Z*,19*Z*-docosapentaenoic acid; 5*Z*,8*Z*,11*Z*,14*Z*-eicosatetraenoic acid,1-glyceryl ester; 5*Z*,8*Z*,11*Z*,14*Z*-eicosatetraenoic acid, 2-glyceryl ester; *N*-(2-hydroxyethyl)-5*Z*,8*Z*,11*Z*,14*Z*-eicosatetraenamide; *N*-(2-hydroxyethyl)-hexadecanamide; *N*-(2-hydroxyethyl)-9*Z*-octadecenamide; 9-oxo-15*S*-hydroxy-prosta-8(12),13*E*-dien-1-oic acid.

### Determination of human recombinant 5-lipoxygenase activity, including studies on reversibility, nuisance inhibition

Human recombinant 5-LOX (Cayman Chemical, 10 U) was pretreated with vehicle (DMSO, 0.1%) or test compounds for 10 min on ice (1 mL PBS pH 7.4, 1 mM EDTA, and 1 mM ATP), and the enzymatic reaction was initiated by addition of AA (20:4) (20 µM; Cayman Chemical) and CaCl_2_ (2 mM).^[22d, 22e]^ Triton X-100 (Thermo Fisher Scientific, Waltham, MA; 0.01%) was added to the reaction buffer to expose nuisance inhibitors. To discriminate between reversible and irreversible 5-LOX inhibitors, the reaction mixture (100 µL) was diluted 10-fold (in PBS pH 7.4, 1 mM EDTA, 1 mM ATP, and either vehicle or test compounds) before the addition of AA (20:4) (20 µM; Cayman Chemical) and CaCl_2_ (2 mM).^[31]^ After 10 min (37 °C), the reaction was stopped (ice-cold methanol; 1 mL), and the internal standard PGB_1_ (200 ng; Cayman Chemical) was added.

Lipid mediators were extracted by solid phase extraction using Clean-Up C-18 Endcapped SPE cartridges (100 mg, 10 mL, UCT, Bristol, PA), conditioned with methanol (1 mL; twice), and equilibrated with water (1 mL). The acidified samples (530 µL PBS plus 60 mM HCl) were centrifuged (750 × g, 10 min, 4 °C), and the supernatants were loaded onto the cartridges, subsequently washed with water (1 mL) and methanol/water (75/25, 1 mL), and the lipid mediators were eluted with methanol (300 µL). The eluates were diluted by adding 120 µL water, centrifuged (21,100 × g, 10 min, 4 °C), and subjected to LC-PDA analysis.^[22d, 22e]^

Chromatographic separation of 5-LOX-derived lipid mediators (all-*trans* isomers of LTB_4_ and 5-HETE, 12-HETE, and 15-HETE) was performed on a Kinetex C-18 LC-column (100 Å, 1.3 μm, 2.1 × 50 mm, Phenomenex, Torrance, CA) using a Nexera X2 UHPLC system (Shimadzu, Kyoto, Japan) operated at 40 °C and a flow rate of 0.45 mL/min using solvent A (50% methanol/50% water/0.05% trifluoroacetic acid) and solvent B (100% methanol/0.05% trifluoroacetic acid) with an initial mobile phase composition of 14% B and 86% A. After 2 min of isocratic elution, the proportion of mobile phase B stepwise increased to 46% (2 min) and then to 90% B (another 2 min). LTB_4_ isomers, 5-HETE, 12-HETE, 15-HETE, and PGB_1_ were quantified using a photodiode array detector (SPD-M20A, Shimadzu) at 280 nm (LTB_4_ isomers and PGB_1_) or 235 nm (HETEs). Chromatograms were acquired and processed using LabSolutions (version 5.97, Shimadzu), and lipid quantities were calculated by one-point internal calibration using PGB_1_.^[22d, 22e]^

### Quantitative analysis of cytokine expression

Freshly isolated human PBMCs (1.4 × 10^6^ cells/mL in 1 mL RPMI 1640 medium supplemented with 5% FCS, 2 mmol/L *L*-glutamine, 100 U/mL penicillin and 100 µg/mL streptomycin) were pretreated (30 min) with vehicle (DMSO, 0.1%), test compounds, or dexamethasone (10 µM) and challenged with LPS (*Escherichia coli* O127:B8; Sigma Aldrich) (10 ng/mL) for 4 h (TNF-α, IL-8) or 18 h (IL-1β, IL-1ra, IL-6, MCP-1, IL-10, IL-12 (p70), IL-23).

Supernatants were collected after centrifugation (21,000 × g, 5 min, 4 °C), and cytokine/chemokine (TNF-α, IL-1β, IL-6, IL-8, MCP-1, IL-10) levels were analyzed using in-house established ELISA systems based on DuoSet ELISA Development kits (Bio-Techne, Minneapolis, MN) according to the manufacturer’s instructions. Alternatively, cytokines (IL-1ra, IL-12 (p70), IL-23) were analyzed using a Bio-Plex 200 System with Bio-Plex Pro Human Cytokine Singleplex Sets, Bio-Plex Pro Reagent Kit III with Flat Bottom Plate, and Bio-Plex Pro Human Cytokine Screening Panel Standards (BIO-RAD, Hercules, CA). Cytokine and chemokine concentrations refer to 8-point standard curves.

Cytokine (TNF-α, IL-1β and IL-10) expression in peritoneal exudates was determined by ELISA using mouse DuoSet ELISA Development kits (Bio-Techne) as outlined above.

### Extraction of membrane and neutral lipids and meroterpenoids

Phospholipids, neutral lipids, sphingolipids, FFA, and compounds **2** and **3** were extracted from cell pellets (PBMCs), peritoneal exudates (150 µL), or mouse plasma (50 µL) by sequential addition of PBS pH 7.4, methanol, chloroform, and saline with a final ratio of 14:34:35:17.^[6, 75]^ The organic phase was evaporated using an Eppendorf Concentrator Plus System (Eppendorf, Hamburg, Germany; high vapor pressure application mode). The resulting lipid film was dissolved in methanol, the sample diluted and centrifuged (21,100 × g, 4 °C, 5 min; twice), and the supernatant subjected to UPLC-MS/MS analysis. Internal standards (Sigma-Aldrich): i. PBMCs (48 h treatment with meroterpenoids): 1,2-dimyristoyl-*sn*-glycero-3-phosphatidylcholine; 1,2-dimyristoyl-*sn*-glycero-3-phosphatidylethanolamine; 1,2-dimyristoyl-*sn*-glycero-3-phosphatidylglycerol; 1,2-myristoyl-*sn*-glycero-3-phosphoserine; 1,2-dioctanoyl-*sn*-glycero-3-phospho-(1’-myo-inositol), (15,15,16,16,17,17,18,18,18-d9)oleic acid; cholest-5-en-3ß-yl heptadecanoate; 1,2,3-trimyristoyl-*sn*-glycerol: 1,2-dimyristoyl-*sn*-glycerol, 1’,3’-bis[1,2-dimyristoyl-*sn*-glycero-3-phospho]-glycerol; *N*-heptadecanoyl-*D*-erythro-sphingosine; and *N*-heptadecanoyl-*D*-erythro-sphingosylphosphorylcholine. ii. DGAT and/or SOAT inhibition (48 h): 1,3-dipentadecanoyl-2-oleyol(d7)-glycerol and 25,26,26,26,27,27,27-d7-cholest-5-en-3β-ol (9Z-octadecenoate). iii. Mouse peritonitis: *D*-erythro-sphingosine-d7, *N*-heptadecanoyl-*D*-erythro-sphingosine, *D*-glucosyl-β-1,1’-*N*-heptadecanoyl-*D*-erythro-sphingosine, *N*-lauroyl-ceramide-1-phosphate, *N*-heptadecanoyl-*D*-erythro-sphingosylphosphorylcholine, and *D*-erythro-sphingosine-d7-1-phosphate.

### Quantitative analysis of neutral and membrane lipids and meroterpenoids by UPLC-MS/MS

Lipids (except for S1P and cholesterol) were separated on an Acquity UPLC BEH C8 column (130 Å, 1.7 μm, 2.1×100 mm, Waters, Milford, MA) using an ExionLC AD UHPLC system (Sciex).^[6, 75a]^ The chromatographic separation of PC, PE, PI, PG, TG, DG, CE, So, sphinganine (Sa), (dh)Cer, hexosylceramide (HexCer), C1P, (dh)SM, and FFA was performed at 45 °C and a flow rate of 0.75 mL/min. For the analysis of PS, the oven temperature was increased to 65 °C and the flow rate to 0.85 mL/min, and CL was separated at a flow rate of 0.60 mL/min and 55 °C. Glycerophospholipids, FFA, and sphingolipids were separated using acetonitrile/water, 95/5, 2 mM ammonium acetate as a mobile phase A and water/acetonitrile, 90/10, 2 mM ammonium acetate as mobile phase B. For the separation of TG, DG, and CE, the mobile phase B was 100% isopropanol. CL were separated using methanol with 2 mM ammonium acetate as mobile phase A and water with 2 mM ammonium acetate as mobile phase B.

For the separation of PC, PE, PI, PG, PS, and FFA, the gradient was ramped from 75% to 85% A within 5 min and to 100% A within another 2 min, followed by 2 min of isocratic elution, which was prolonged to 13 min for the analysis of sphingolipids. TG, DG, and CE were separated using a mobile phase composition of A/B = 90/10, which was ramped to A/B = 70/30 within 6 min, followed by 4 min of isocratic elution.^[6]^ For the elution of CL, the proportion of mobile phase A was increased from 85% to 98% over 8 min, followed by isocratic elution (1 min).

Eluted lipids were detected using a QTRAP 6500^+^ mass spectrometer (Sciex) equipped with an IonDrive Turbo V Ion Source and a TurboIonSpray probe for electrospray ionization. PC ([M+OAc]^-^ to fatty acid anions),^[76]^ other glycerophospholipids ([M-H]^-^ to fatty acid anions),^[76]^ and CL ([M-2H]^2-^ to fatty acid anions) were detected by MRM in the negative ion mode. FFA were quantified by single ion monitoring in the negative ion mode.^[6]^ TG, DG, and CE were analyzed by MRM ([M+NH_4_]^+^ to [M-fatty acid anion]^+^) in the positive ion mode.^[6, 77]^ Transitions from [M+H]^+^ to [M+H-H_2_O]^+^ (So, Sa, dhCer), to m/z = 184.1 ([dh]SM), and to m/z = 264.4 (Cer, HexCer, C1P) were selected for the quantitation of sphingolipids by scheduled MRM.^[75a]^ The source and compound-specific parameters in positive and negative ion mode are listed in **Table 3**.

**Table 3.**
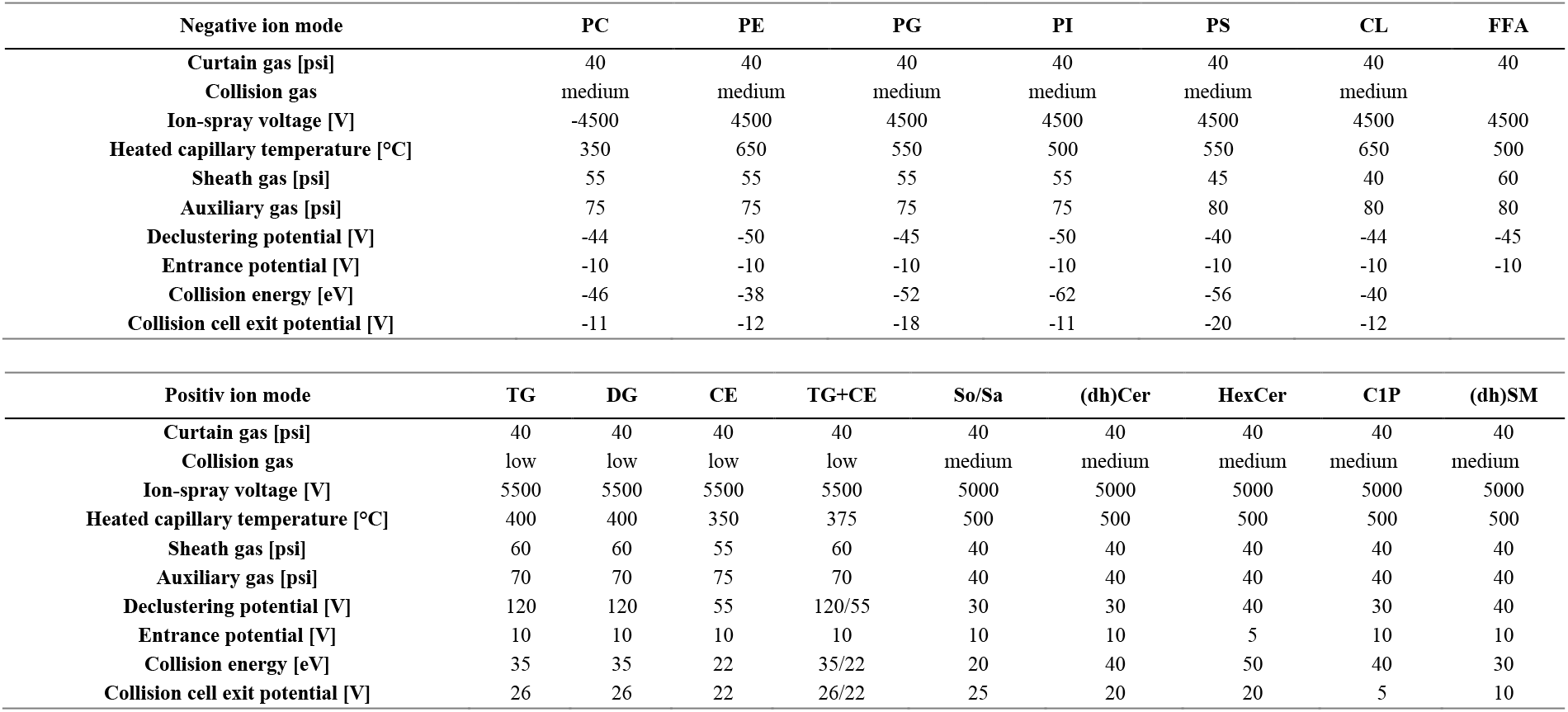
QTRAP 6500^+^ parameters for targeted lipidomics.

Deviating from the procedures described above, S1P, cholesterol, and compounds **2** and **3** were separated on an Acquity UPLC CSH C18 column (130Å, 1.7 μm, 2.1 × 50 mm, Waters) at 55 °C and a flow rate of 0.55 mL/min. A mixture of water/acetonitrile/isopropanol (32/20/48) with 0.1% formic acid was used as mobile phase for the isocratic elution of S1P, **2**, and **3**. For the elution of cholesterol, the initial conditions of 40% mobile phase A (water/acetonitrile (80/20) with 0.1% formic acid) and 60% mobile phase B (isopropanol/acetonitrile (80/20) with 0.1% formic acid) were held for 3 min before the proportion of B was increased linearly to 70% within 2 min and then further to 100% within 0.4 min, followed by isocratic elution for 1.6 min.

S1P, cholesterol, and compounds **2** and **3** were detected in the positive ion mode using a QTRAP 6500^+^ mass spectrometer (Sciex) equipped with an IonDrive Turbo V Ion Source and a TurboIonSpray probe for electrospray ionization. Transitions from m/z 380.3 → 264.2 (S1P; [M+H]^+^ to [M+H-H_3_PO_4_-H_2_O]^+^), m/z 359.2 → 169.3 (**2**), m/z 345.2 → 153.0 (**3**), and m/z 387.4 ([M+H]^+^) to 281.0 ([M-C_6_H_15_O-H_2_]^+^) (cholesterol) were selected for quantitation using the instrument parameters listed in **Table 4**.

**Table 4.** QTRAP 6500^+^ parameters for the analysis of S1P, cholesterol, 2 and 3.

| Positiv mode | S1P | Cholesterol | **2** | **3** |
| --- | --- | --- | --- | --- |
| Curtain gas [psi] | 40 | 40 | 40 | 40 |
| Collision gas | low | low | low | low |
| Ion-spray voltage [V] | 5500 | 5500 | 4500 | 5500 |
| Heated capillary temperature [°C] | 550 | 500 | 550 | 550 |
| Sheath gas [psi] | 60 | 45 | 60 | 60 |
| Auxiliary gas [psi] | 30 | 45 | 30 | 30 |
| Declustering potential [V] | 40 | 20 | 30 | 40 |
| Entrance potential [V] | 10 | 5 | 10 | 10 |
| Collision energy [eV] | 20 | 10 | 25 | 30 |
| Collision cell exit potential [V] | 20 | 20 | 10 | 15 |

Absolute and relative amounts of individual PC, PE, PG, PI, PS, CE, FFA, DG and sphingolipid species were calculated from the mean of the analyzed transitions. Signals from transitions indicating fatty acids occurring at multiple positions were first divided by the number of positions where these fatty acids occur. Quantification of TG and CL was instead based on the most abundant and specific transition corrected for repeatedly occurring fatty acids as described above. Glycerophospholipids, neutral lipids, FFA, and sphingolipids were quantified by one-point calibration based on a subclass-specific internal standard and normalized to cell number or sample volume. Sphingolipid signals from murine plasma and exudates (except for S1P) were normalized to a subclass-specific internal standard, and concentrations were calculated based on hyperbolic 11-point external calibrations using subclass-specific standards. Concentrations of **2** and **3** in mouse plasma and peritoneal exudates were calculated from a linear 10-point standard curve, normalized to an internal standard (*D*-erythro-sphingosine-d7-1-phosphate), the recovery of **3** for liquid-liquid extraction, and the sample volume.

Relative intensities (i.e., proportions) of lipids were calculated by summarizing the absolute amounts of PC, PE, PG, PI, PS, CL, TG, DG, CE, FFA, and sphingolipid species (= 100%), respectively and expressing the amount of each lipid species as a percentage of this sum. The cellular proportion of lipids containing a specific fatty acid (e.g., 20:4) was calculated by summarizing the proportions of individual lipids containing the respective fatty acid (after correction for repeatedly occurring fatty acids) and expressing this sum as a percentage of total lipids (= 100%). Mass spectra were acquired and processed using Analyst 1.7.1 or 1.7.2 and 1.6.3 (Sciex).

### Zymosan-induced peritonitis in mice

Male CD-1 mice (33-39 g, eight weeks old, Charles River Laboratories, Calco, Italy) were provided with standard rodent chow and water and allowed to acclimate for four days in a controlled environment with a 12-h light/12-h dark schedule at a constant temperature of 21 ± 2 °C. All experiments were performed during the light phase, and mice were randomly assigned to different experimental groups. Mice received vehicle (2% DMSO in saline; 0.5 ml), **2** (10 mg/kg), and **3** (10 mg/kg) intraperitoneally (i.p.) 30 min prior to injection of zymosan (Sigma-Aldrich; 2 mg/ml in saline, 0.5 ml, i.p.). Mice were sacrificed (by CO_2_ inhalation) after 4 h or 18 h, and peritoneal exudates and plasma were collected.^[78]^ Cell numbers in the exudates were determined after vital trypan blue staining.^[78]^ Lipid mediators (in 100 µL plasma and 500 µL exudate), sphingolipids (in 50 µL plasma and 150 µL exudate), and compounds **2** and **3** (in 50 µL plasma and 150 µL exudate) were extracted and quantified as described above.

### Determination of the activity of endocannabinoid degrading enzymes

The activity of human recombinant FAAH (#10005196) and MAGL (#705192) was determined using commercially available inhibitor screening assay kits (Cayman Chemical) according to the manufacturers’ instructions.

### Immunofluorescence microscopy

Lipoxygenases and perilipin-2 were visualized in immune cells according to a procedure adapted from Jordan et al.^[29]^ M2 macrophages (8 × 10^5^) in RPMI 1640 medium supplemented with 10% FCS, 2 mmol/L *L*-glutamine, 100 U/mL penicillin, and 100 µg/mL streptomycin were seeded onto glass coverslips (Thermo Fisher Scientific) coated with poly-*D*-lysine hydrobromide (Sigma Aldrich; 30 min, 37 °C, 5% CO_2_). After 30 min, the medium was replaced with PBS pH 7.4 plus 1 mM CaCl_2_, and the attached cells were treated with vehicle (DMSO, 0.1%), SACM (0.5%), **2** (10 µM), or **3** (10 µM) for 90 min (37 °C, 5% CO_2_). Cells were fixed with 4% paraformaldehyde in PBS pH 7.4 (Sigma Aldrich, 20 min, RT), permeabilized with acetone (3 min, 4 °C), and treated with 0.1% Triton X-100 in PBS pH 7.4 plus 1 mM CaCl_2_ (Thermo Fisher Scientific, 10 min, RT). After blocking with 10% Normal Goat Serum in PBS pH 7.4 with 0.1% sodium azide (Thermo Fisher Scientific) for 30 min (RT), the coverslips were incubated overnight (4 °C) with a mouse monoclonal anti-15-LOX-1 antibody (1:100, # ab119774, Lot: GR3322776-8; Abcam, Cambridge, UK), washed repeatedly, and stained with Alexa Fluor™ 555 goat anti-mouse IgG (1:500, # A21422, Lot: 10143952; Invitrogen, Carlsbad, CA) for 30 min (RT).

PBMCs (3.65 × 10^6^) in RPMI 1640 medium supplemented with 5% FCS, 2 mmol/L *L*-glutamine, 100 U/mL penicillin, and 100 µg/mL streptomycin were seeded onto poly-*D*-lysine-coated glass coverslips, incubated for 30 min (37 °C, 5% CO_2_), and treated with vehicle (DMSO, 0.1%), **3**, or the indicated inhibitors for 48 h. The culture medium was replaced with PBS pH 7.4 plus 1 mM CaCl_2_, and the cells were challenged with the calcium ionophore A23187 (5 min, 37 °C). After fixation (4% paraformaldehyde in PBS pH 7.4, Sigma Aldrich; 20 min, RT) and permeabilization of the cells (0.1% Triton X-100 in PBS pH 7.4 plus 1 mM CaCl_2_, Thermo Fisher Scientific; 15 min, RT), samples were blocked with 10% Normal Goat Serum in PBS pH 7.4 with 0.1% sodium azide (Thermo Fisher Scientific) for 30 min (RT) and incubated overnight (4 °C) with mouse anti-5-LOX (1:1000; # 610695; Lot: 9067817; BD Biosciences, Franklin Lakes, NJ) and rabbit anti-perilipin-2 (1:100; # ab108323, Lot: GR3259785-40; Abcam) antibodies. 5-LOX was stained with Alexa Fluor™ 555 goat anti-mouse IgG (1:500, # A21422, Lot: 10143952; Invitrogen), and perilipin-2 was stained with Alexa Fluor™ 488 goat anti-rabbit IgG (1:500, # 11034, Lot: 10729174; Invitrogen) for 30 min (RT).

Samples were mounted with ProLong™ Gold Antifade Mountant with DAPI (Thermo Fisher Scientific) and visualized using a Zeiss AxioObserver Z1 microscope (Carl Zeiss, Jena, Germany) operated with Zen 2.6 (Carl Zeiss) and equipped with a Plan-Apochomat 40x/1.4 Oil DIC (UV) VIS-IR M27 objective (Carl Zeiss) and an Axiocam 702 (Carl Zeiss). The exposure time was kept constant for all acquired images. Maximum intensity projections of Z stacks (8-10 slices, Z = 1 µm) (Figure 3H, Figure 8B and Figure S23, Supporting Information) or single slices (Z = 1 µm) (Figure S24, Supporting Information) were exported with ZEN blue 3.7 (Carl Zeiss). For merged images, the DAPI channel was pseudo-colored light blue, Alexa 555 was pseudo-colored yellow (15-LOX-1) or violet (5-LOX), and Alexa 488 was pseudo-colored green (perilipin-2). 15-LOX-1 translocation and the number of lipid bodies were quantitatively assessed using Fiji^[79]^ by counting cellular particles stained for 15-LOX-1 (Threshold: 0-4000; Particle size: 0-infinity; Circularity: 0-1) and perilipin-2 (Threshold: n1 600-65535, n2 500-600, n3 325-65535; Particle size: 0-infinity; Circularity: 0-1), respectively.

### Determination of protein expression and phosphorylation by SDS-PAGE and Western blotting

Freshly isolated PBMCs (3.84 × 10^6^) were serum-starved overnight in RPMI 1640 medium supplemented with 2 mmol/L *L*-glutamine, 100 U/mL penicillin, and 100 µg/mL streptomycin and then treated with vehicle (DMSO, 0.1%) or test compounds in the presence of 2% FCS at 37 °C and 5% CO_2_. After 30 min, cells were stimulated with 10 ng/mL LPS (*Escherichia coli* O127:B8; Sigma Aldrich) for 15 min or 1 h to analyze the expression of IκBα, p-IκBα, and NF-κB. Cells were placed on ice, washed with ice-cold PBS pH 7.4 (twice, 1 mL), mixed with 100 µL lysis buffer (20 mM Tris-HCl pH 7.4, 150 mM NaCl, 2 mM EDTA, 1% Triton X-100, 5 mM sodium fluoride, 10 μg/mL leupeptin, 60 μg/mL soybean trypsin inhibitor, 2.7 mM sodium vanadate, 2.5 mM sodium pyrophosphate, and 1 mM phenylmethanesulfonyl fluoride), and sonicated (2 × 5 s, on ice, power: 35% of 125 W; Q125 Sonicator, QSonica, Newtown, CT).

For the analysis of 5-LOX, FLAP, and MEK1/2 expression and MEK1/2 phosphorylation, PBMCs (1 × 10^7^) were resuspended in RPMI 1640 medium supplemented with 5% FCS, 2 mmol/L *L*-glutamine, 100 U/mL penicillin, and 100 µg/mL streptomycin and preincubated with vehicle (DMSO, 0.1%) or test compounds at 37 °C for 48 h. Cells were harvested by successive rinses with PBS pH 7.4 plus 5 mM EDTA (ice-cold; 1 mL) and PBS pH 7.4 (ice-cold; 1 mL), washed twice with ice-cold PBS pH 7.4, and centrifuged (2000 × g, 7 min, 4 °C). The pellet was resuspended in 100 µL lysis buffer (20 mM Tris-HCl pH 7.4, 150 mM NaCl, 2 mM EDTA, 1% Triton X-100, 5 mM sodium fluoride, 10 μg/mL leupeptin, 60 μg/mL soybean trypsin inhibitor, 1 mM sodium vanadate, 2.5 mM sodium pyrophosphate, and 1 mM phenylmethanesulfonyl fluoride) and sonicated (2 × 5 s, on ice, power: 35% of 125 W; Q125 Sonicator, QSonica).

Cell lysates were centrifuged (1200 × g, 5 min, 4 °C), total protein levels were determined using a DC protein assay kit (Bio-Rad Laboratories GmbH, Munich, Germany), and equal protein concentrations were adjusted. Samples were mixed with 5× SDS/PAGE sample loading buffer (125 mM Tris-HCl pH 6.5, 25% sucrose, 5% SDS, 0.25% bromophenol blue, and 5% β-mercaptoethanol) and heated at 95 °C for 5 min. Aliquots (6-30 µg protein depending on the target protein) were resolved on SDS-PAGE gels (10%), and proteins were transferred to Cytiva Amersham™ Protran™ 0.45 µm NC nitrocellulose blotting membranes (GE Healthcare, Munich, Germany). After blocking with 5% bovine serum albumin (Roth, Karlsruhe, Germany) or 5% skim milk (Sigma Aldrich) in TBS pH 7.4 plus 1% Tween-20 for 1 h (RT), the membranes were incubated with primary antibodies at 4 °C overnight. The following primary antibodies were used: mouse anti-IκBα (1:800; # 4814S; Cell Signaling, Danvers, MA), rabbit anti-phospho-IκBα (1:1000; # 2859; Cell Signaling), rabbit anti-NF-κB (1:500; # 4764; Cell Signaling), mouse anti-5-LOX (1:1000; # 610695; Lot: 9067817; BD Biosciences), rabbit anti-MEK1/2 (1:1000; # 9122; Cell Signaling), rabbit anti-phospho-MEK1/2 (1:1000; #9121; Cell Signaling), goat anti-FLAP (1:1000; # ab53536; Lot: GR67416-18; Abcam), rabbit anti-β-actin (1:1000; #4970; Lot: 19; Cell Signaling), and mouse anti-β-actin (1:1000; # 3700; Lot: 21; Cell Signaling). Membranes were washed three times with TBS pH 7.4 plus 1% Tween-20 and incubated with fluorophore-conjugated secondary antibodies for 1 h (RT). Secondary antibodies used: DyLight® 800 anti-mouse IgG (1:10000; # SA5-10176; Lot: VE2995014; Thermo Fisher Scientific), DyLight® 800 anti-rabbit IgG (1:10000; # SA5-10036; Lot: VC2960806; Thermo Fisher Scientific), DyLight® 680 anti-mouse IgG (1:10000; # 35519; Lot: VB298075; Thermo Fisher Scientific), DyLight® 680 anti-rabbit IgG (1:10000; # 35569; Lot: VB302113; Thermo Fisher Scientific), and Alexa Fluor® 680 anti-goat IgG (1:8333; #ab175776; Lot: GR3459994-1; Abcam).

Fluorescence signals were detected using a Fusion FX7 Edge Imaging System (spectra light capsules: C680, C780; emission filters: F-750, F-850; VILBER Lourmat, Collegien, France). Densitometric analysis was performed using the Bio-1D imaging software (version: 15.08c (VILBER Lourmat) applying a rolling ball background subtraction, and protein levels were normalized to β-actin or total kinase. Uncropped blots are shown in Figure S11 and Figure S25B, Supporting Information.

### Statistics and network analysis

Data are expressed as mean, mean ± SEM, or mean ± SEM and single data from *n* independent experiments or animals (zymosan-induced peritonitis). For data with similar variance between groups, Shapiro-Wilk tests were applied to test for normal distribution, and outliers were detected using Grubb’s tests, except when n = 3. *P* values < 0.05 were considered statistically significant. Non-transformed or logarithmized data were used for statistical analysis. Two-tailed (two-sided α level of 0.05) paired or unpaired Student *t* tests were used for pairwise comparisons, and one-way ANOVAs, two-way ANOVAs, or mixed-effects models (REML) were used to compare independent or correlated samples, followed by Dunnett *post hoc* tests. Negative log_10_(q values) for volcano plots were calculated using two-tailed multiple (un)paired Student *t* tests with correction for multiple comparisons using a two-stage step-up procedure by Benjamini, Krieger, and Yekutieli. Data were analyzed using Microsoft Excel Version 2302 (Microsoft 365 apps for enterprise, Microsoft, Redmond, WA), and statistics were evaluated using GraphPad Prism 9.5 or newer (GraphPad Software, San Diego, CA). Principal component analysis was performed using Origin 2022b SR1 (OriginLab, Northampton, MA).

The correlation-based network was constructed using the MetScape 3.1.3 plugin for Cytoscape 3.9.1 (Cytoscape Consortium).^[80]^ Pearson correlation coefficients were calculated from the mean percentage changes in absolute lipid amounts (1 and 10 µM). Each node represents a single lipid species and edges represent correlation factors ≥ 0.7 (Layout: Edge-weighted Spring Embedded Layout based on correlation from matrix). Correlations between individual lipid species and lipid mediator subgroups (5-LOX products, EETs) or AA (20:4) were calculated (Pearson correlation) in Microsoft Excel Version 2302 (Microsoft 365 apps for enterprise, Microsoft). Positive correlations ≥ 0.7 are depicted in red and negative correlations ≤ -0.7 are depicted in blue.

## Supporting Information

Supporting Information is available from the Wiley Online Library.

## Conflict of Interest

The authors declare no conflict of interest.

## Authors Contributions

Lorenz Waltl – Methodology, Investigation, Visualization, Writing - Original Draft, Writing - Review & Editing. Klaus Speck - Methodology, Investigation, Analysis. Raphael Wildermuth - Methodology, Investigation. Franz-Lucas Haut - Methodology, Investigation. Stephan Permann - Methodology, Investigation, Analysis. Danilo D’Avino - Methodology, Investigation. Ida Cerqua – Methodology, Investigation. Anita Siller – Resources. Harald Schennach – Resources. Antonietta Rossi - Methodology, Investigation, Supervision, Funding acquisition. Thomas Magauer – Resources, Writing - Review & Editing, Supervision, Funding acquisition. Andreas Koeberle: Project administration, Conceptualization, Data Curation, Writing - Original Draft, Writing - Review & Editing, Supervision, Funding acquisition.

## Ethics Approval Statement

Studies on PBMC and macrophages were approved by the ethical commission of the Medical University Innsbruck (no. 1041/2020 from June 19^th^, 2020) and performed with informed consent of the platelet donors. The experimental procedures for studies related to zymosan-induced peritonitis in mice followed the guidelines outlined in the Italian (DL 26/2014) and European (Directive 2010/63/EU) regulations on the ethical use and protection of animals for scientific purposes, complied with the ARRIVE guidelines for reporting *in vivo* experiments, and were approved by the Italian Ministry.

### Acknowledgements

The authors thank Gabriell Knoll for technical assistance in performing experimental methods and data analysis.

## Funding Statement

Research activities of A.K. related to the subject of this article were funded in part by the Austrian Science Fund (FWF) (I 4968, P 36299). T.M. acknowledges the European Research Council under the European Union’s Horizon 2020 research and innovation program (grant agreement No 714049, No 101000060) and the Center for Molecular Biosciences (CMBI). A.R. acknowledges Ministero dell’Università e della Ricerca (MUR) PRIN2022 PNRR (P2022CNPH8). For the purpose of open access, the authors have applied a CC BY public copyright license to any Author Accepted Manuscript version arising from this submission.

## Notes

### Competing Interest Statement

The authors have declared no competing interest.

