## Supporting information for "Reorganization of innate immune cell lipid profiles by bioinspired meroterpenoids to limit inflammation"

Waltl et al.

### Supplementary figures

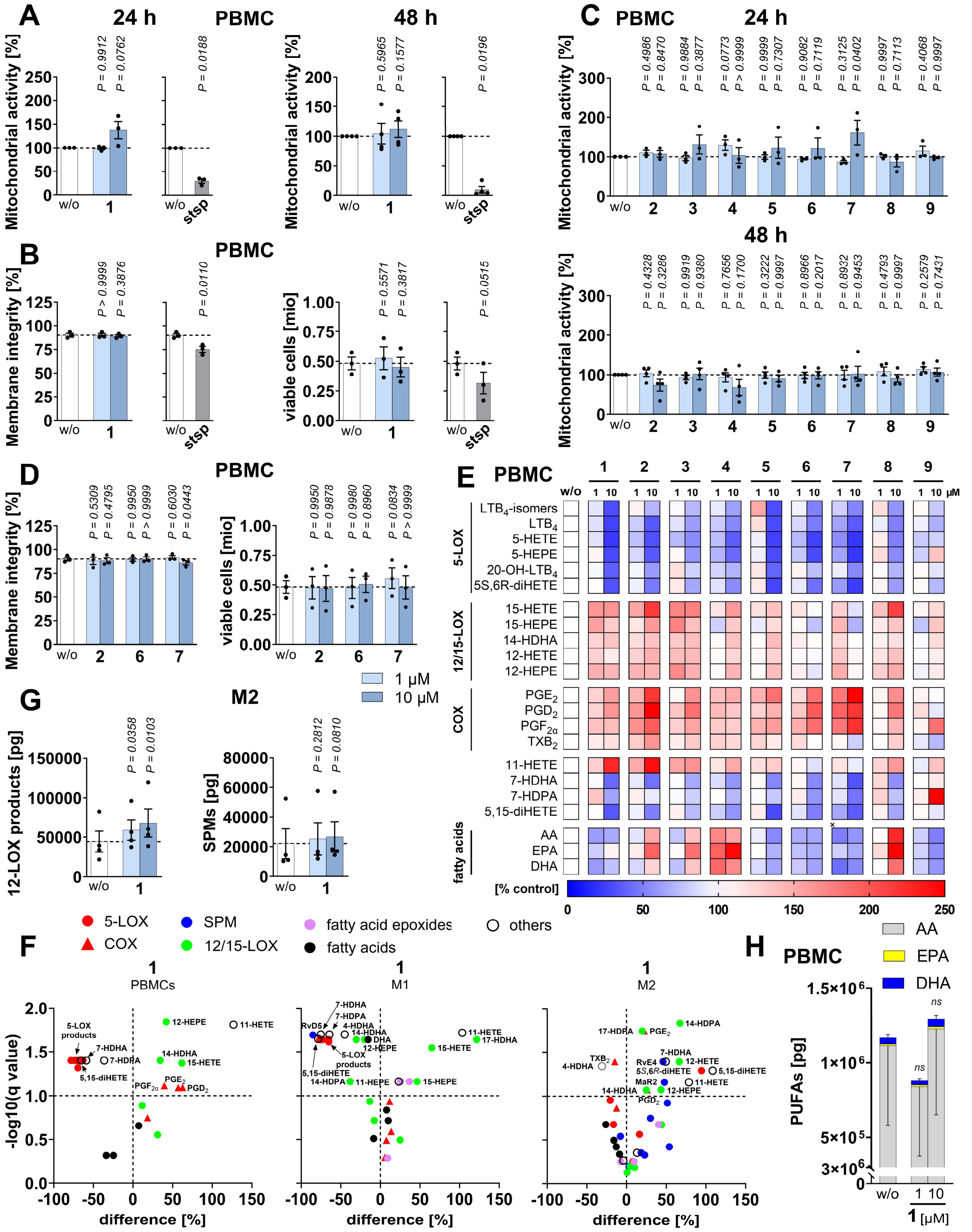

**Figure S1.** Meroterpenoids alter the lipid mediator profile of activated innate immune cells without inducing cytotoxicity. A,C) Changes in cellular dehydrogenase activity of PBMCs treated with vehicle (DMSO, 0.1%) or meroterpenoids for 24 h or 48 h (MTT assay). The vehicle control shown in C is identical to panel A. B,D) Membrane integrity of PBMCs treated with vehicle (DMSO, 0.1%) or meroterpenoids. E,F,H) PBMCs were preincubated (10 min) with vehicle (DMSO, 0.1%) or meroterpenoids and then challenged with A23187 (10 min). E) Heatmap showing percentage changes in the concentration of lipid mediators relative to vehicle control. F,G) M1 or M2 macrophages were preincubated (15 min) with vehicle (DMSO, 0.1%) or **1** (10 µM, if not indicated otherwise). F) 12-LOX products and SPMs [per $2\times10^6$ M2]. G) Volcano plots showing the mean percentage difference relative to vehicle control and the negative $\log_{10}$(q value) calculated vs. vehicle control; two-tailed multiple paired student *t* tests with correction for multiple comparisons (false discovery rate 10%). H) Concentration of the free PUFAs AA (20:4), eicosapentaenoic acid (EPA), and docosahexaenoic acid (DHA) [per $5 \times 10^6$ PBMCs]. Mean (E,F) or mean + SEM (H) and single data (A-D,G) from n = 3 (A 24 h,B,C 24 h,D), n = 3-4 (E,F,H), n = 4 (A 48 h,C 48 h,G) independent experiments. *P* values given vs. vehicle control (A-D, F-H); repeated measures one-way ANOVA of log data (A-D,G) + Dunnett *post hoc* tests or mixed-effects model (REML) of log data (H) + Dunnett *post hoc* tests. Staurosporine ('stsp', 1 µM) was used as cytotoxic control, two-tailed paired Student *t* test.

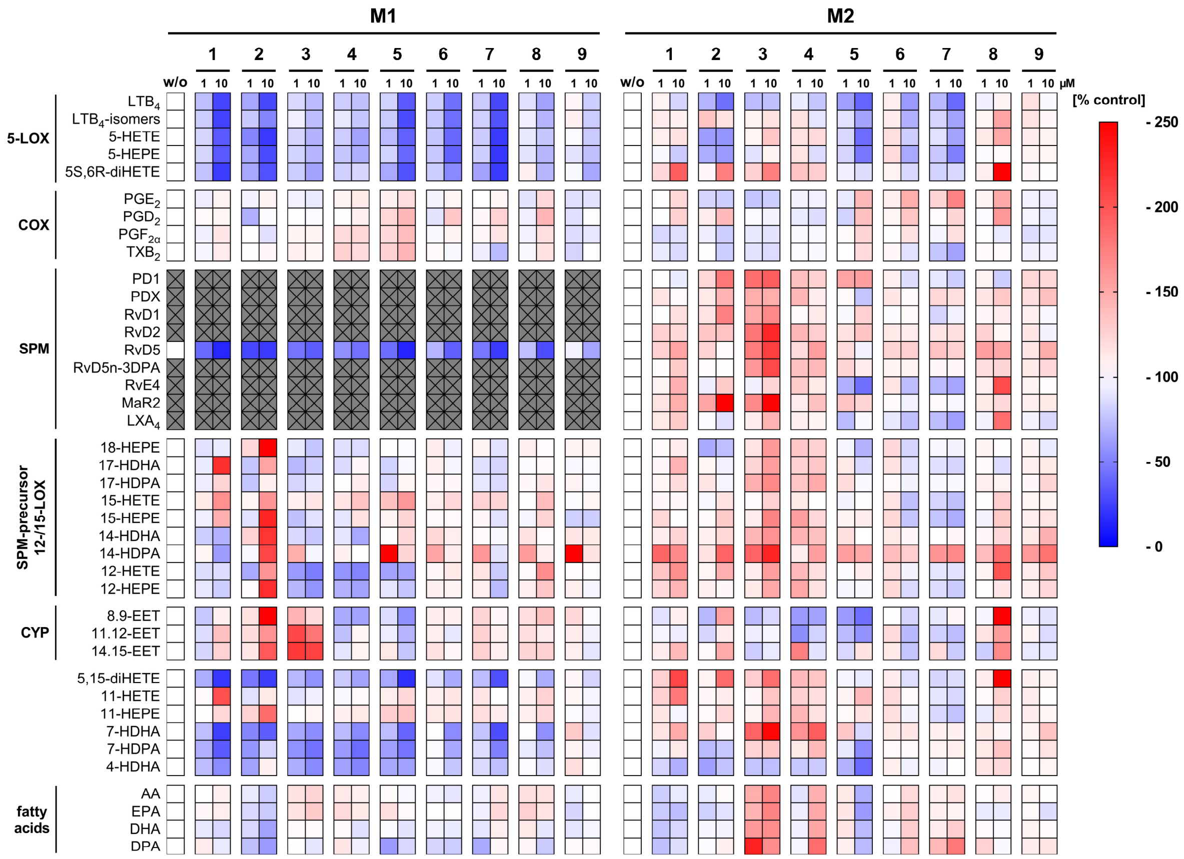


**Figure S2.** Effect of meroterpenoids on the lipid mediator profile of activated macrophages. M1 or M2 macrophages were preincubated (15 min) with vehicle (DMSO, 0.1%) or test compounds and activated with SACM (180 min). Heatmap showing percentage changes in the concentration of lipid mediators relative to vehicle control. Mean from n = 3-4 independent experiments.

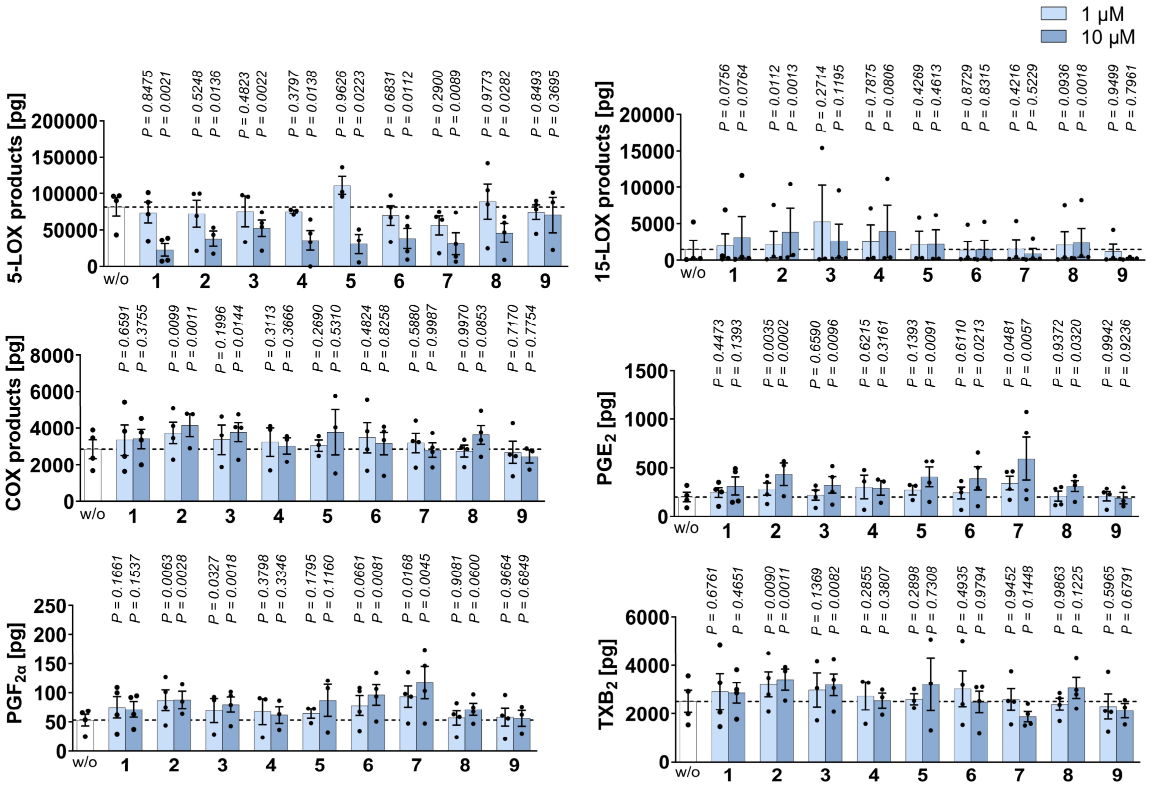


**Figure S3**. Effect of meroterpenoids on the biosynthesis of lipid mediator classes and selected species in activated innate immune cells. PBMCs were preincubated (10 min) with vehicle (DMSO, 0.1%) or **1–9** and then challenged with A23187 (10 min). 5-LOX products, 15-LOX products. COX products, and selected prostanoids [per 5 × $10^6$ PBMCs]. Data for the vehicle control and **1** (5-LOX and 15-LOX) are identical to Figure 1A. Mean + SEM and single data from n = 3-4 independent experiments. *P* values given vs. vehicle control; repeated measures one-way ANOVA of log data or mixed-effects model (REML) of log data + Dunnett post hoc tests.

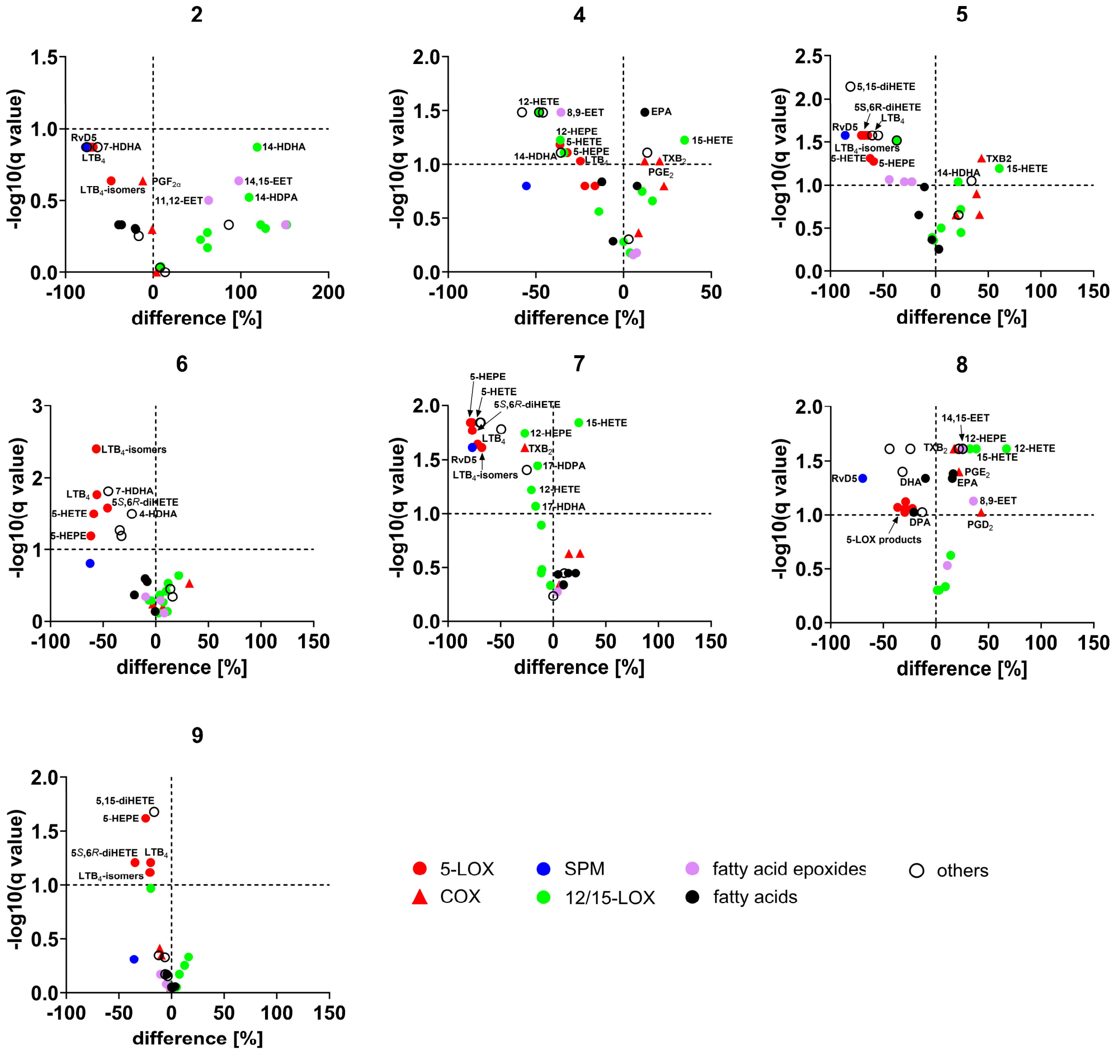


**Figure S4.** Extent and statistical significance of the effect of **2**, **4-9** on the levels of lipid mediators produced by activated M1 macrophages. Cells were preincubated (15 min) with vehicle (DMSO, 0.1%) or test compounds (10 μM) and activated with SACM (180 min). Volcano plot showing the mean percentage difference relative to vehicle control and the negative $\log_{10}$(q value) calculated vs. vehicle control from n= 3-4 independent experiments; two-tailed multiple paired Student *t* tests with correction for multiple comparisons (false discovery rate 10%).

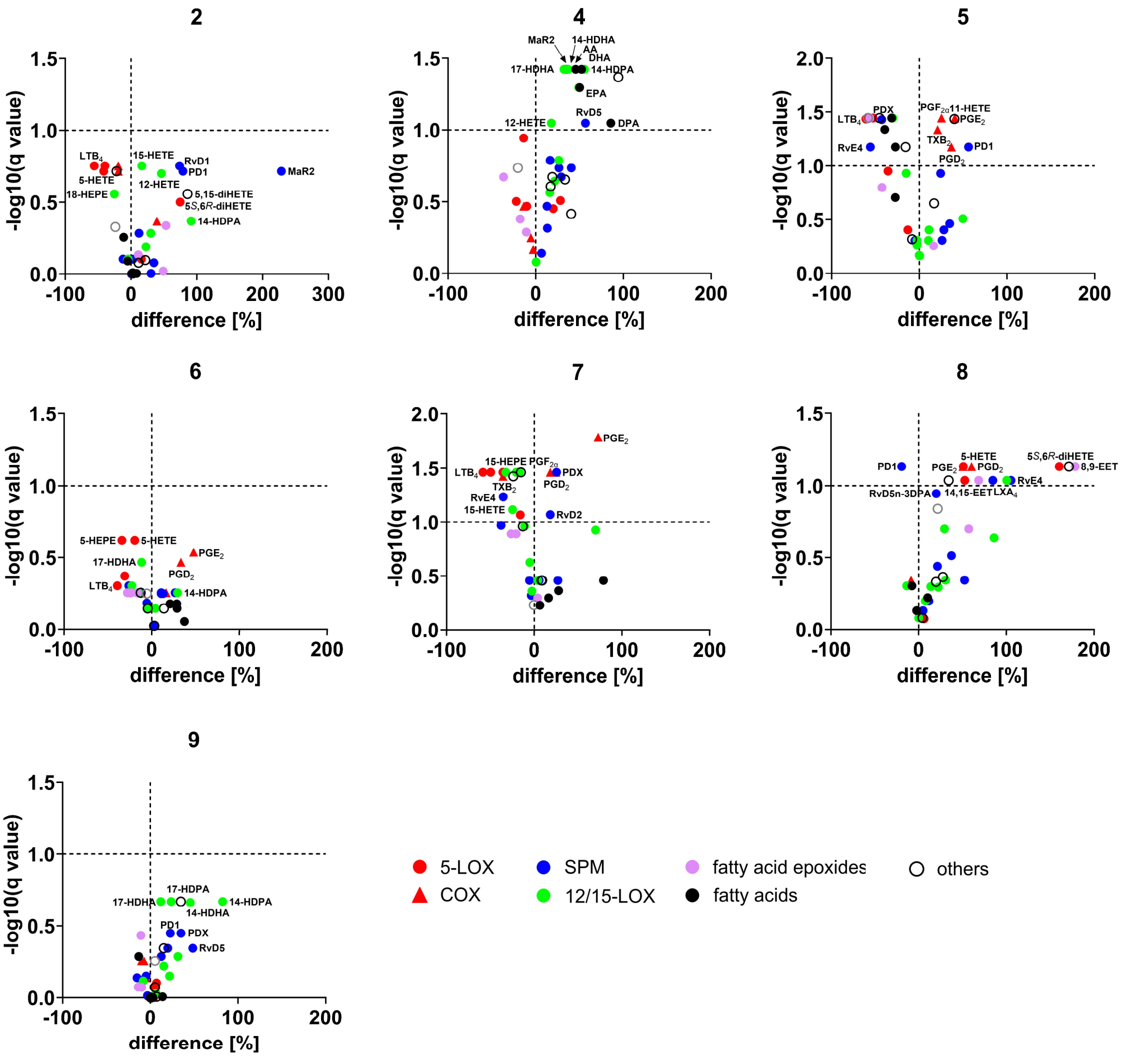


**Figure S5.** Extent and statistical significance of the effect of **2**, **4-9** on the levels of lipid mediators produced by activated M2 macrophages. Cells were preincubated (15 min) with vehicle (DMSO, 0.1%) or test compounds (10 μM) and activated with SACM (180 min). Volcano plot showing the mean percentage difference relative to vehicle control and the negative $\log_{10}$(q value) calculated vs. vehicle control from n= 3-4 independent experiments; two-tailed multiple paired Student *t* tests with correction for multiple comparisons (false discovery rate 10%).

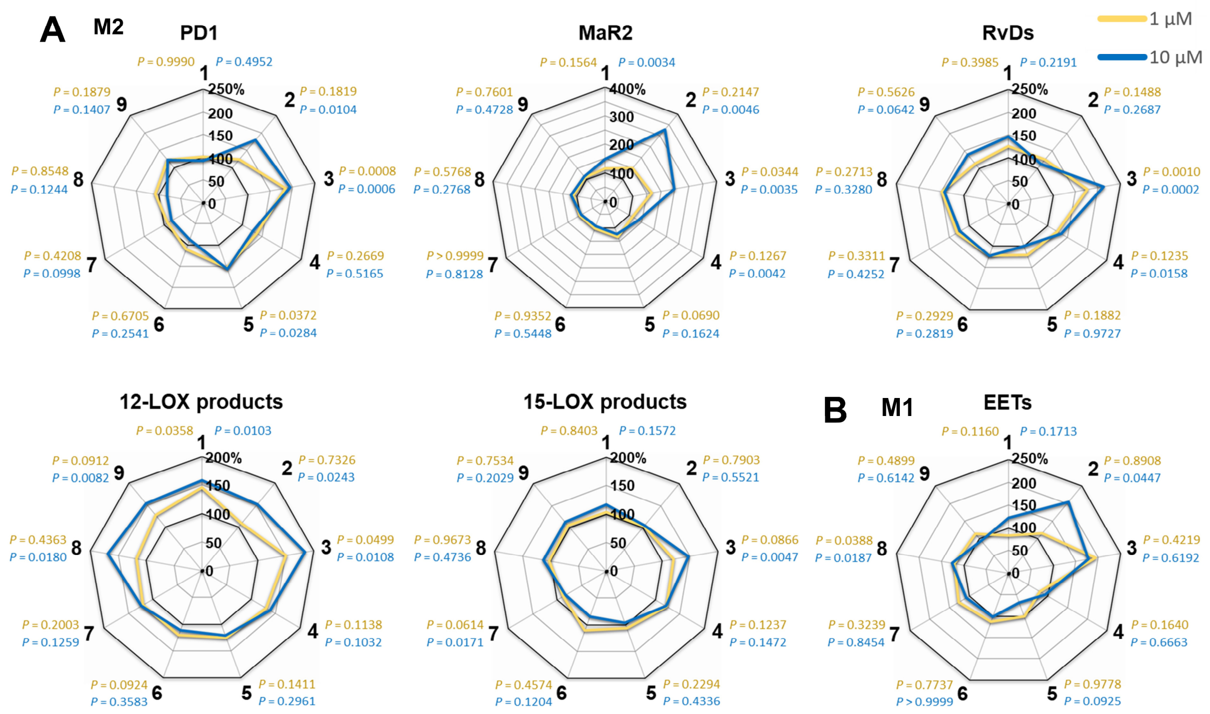


**Figure S6**. Effect of meroterpenoids on the biosynthesis of lipid mediator classes and selected species in activated macrophages. M1 or M2 macrophages were preincubated (15 min) with vehicle (DMSO, 0.1%) or test compounds and activated with SACM (180 min). A PD1, MaR2, RvDs, 12- and 15-LOX products produced by M2 macrophages. Data for **1** are identical to Figure 1B-D. B) EETs produced by M1 macrophages. Data is identical to Figure 2E. Mean from n = 3-4 independent experiments. *P* values given vs. vehicle control; repeated measures one-way ANOVA of log data or mixed-effects model (REML) of log data + Dunnett post hoc tests.

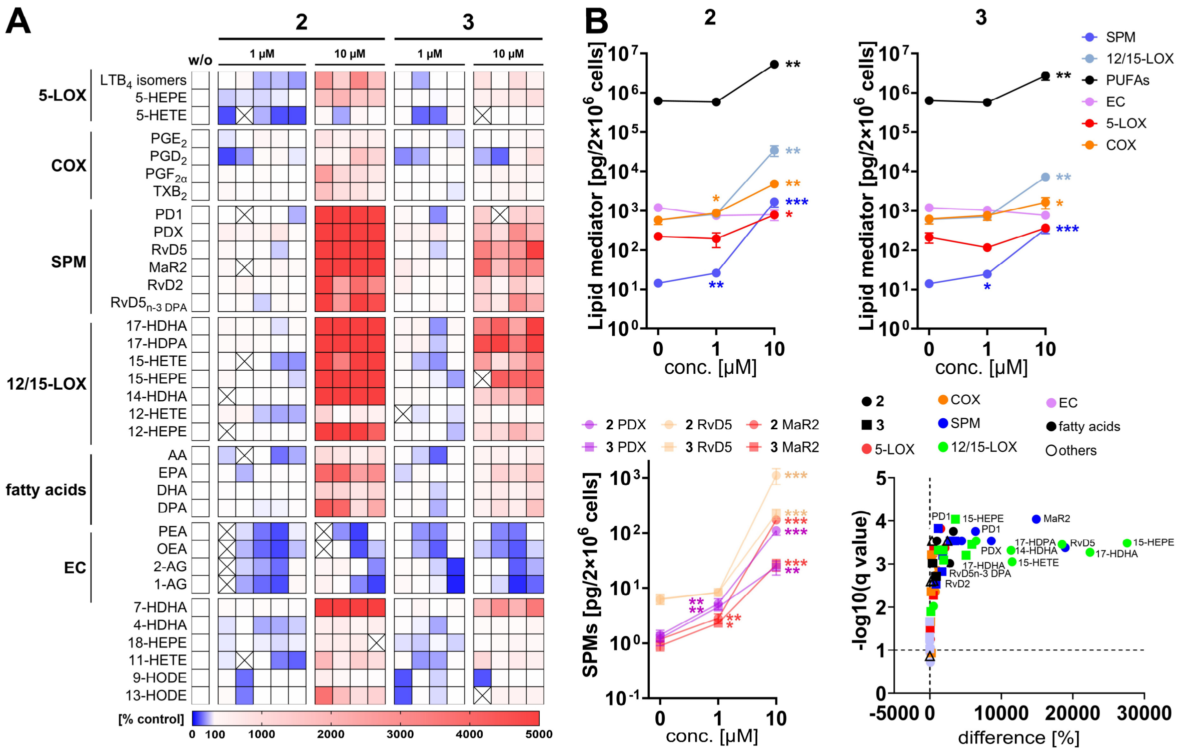


**Figure S7.** Effect of **2** and **3** on the lipid mediator profile of resting M2 macrophages. Cells were preincubated with vehicle (DMSO, 0.1%), **2** or **3** (10 µM unless stated otherwise) for 195 min**.** A) Heatmap showing percentage changes in lipid mediator levels relative to vehicle control for each independent experiment, with the crossed sections indicating significant outliers. B) Line charts showing the levels of lipid mediator subclasses or of individual SPMs and a volcano plot showing the mean percentage difference relative to vehicle control and the negative $\log_{10}$(q value) calculated vs. vehicle control; two-tailed multiple paired Student *t* tests with correction for multiple comparisons (false discovery rate 10%). Mean (A,B) or mean + SEM (B) from n = 3-5 independent experiment*s*. **P* < 0.05, ***P* < 0.01, ****P* < 0.001 vs. vehicle control (B); two-tailed paired Student *t* test of log data (B).

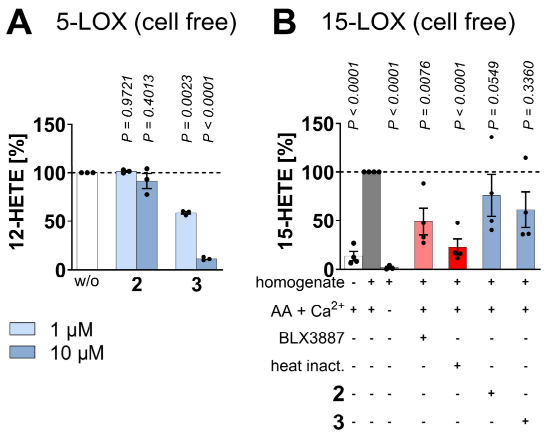


**Figure S8.** Effect of **2** and **3** on positional side reactions of human 5-LOX and 15-LOX activity. A) Human recombinant 5-LOX was pretreated (10 min) with vehicle (DMSO, 0.1%) or test compounds, and product formation was initiated with AA (20:4) and $CaCl_2$ (10 min). Conversion of AA (20:4) to 12-HETE by 5-LOX. B) 15-HETE formation (indicative of 15-LOX activity) in M2 homogenates that were preincubated with vehicle (DMSO, 0.1%) or test compounds (10 µM, 15 min) and then treated with AA (20:4) and $CaCl_2$ (15 min). Mean + SEM and single data from n = 3 (A), n = 3-4 (B) independent experiments. *P* values given vs. vehicle control; repeated measures one-way ANOVA of log data (A) or mixed-effects model (REML) of log data (B) + Dunnett post hoc tests.

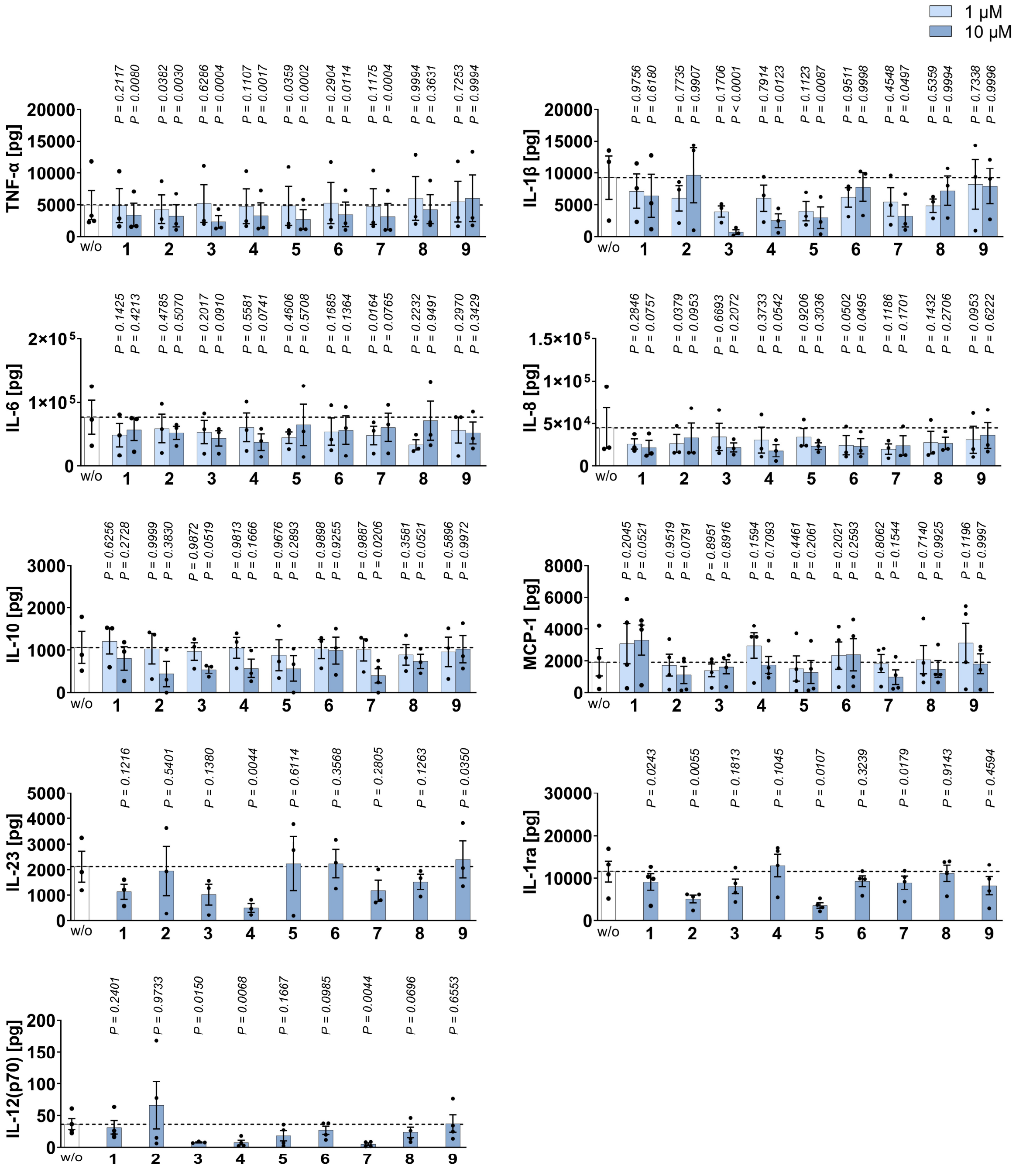


**Figure S9.** Effect of meroterpenoids on cytokine and chemokine expression of PBMCs. Cells were pretreated (30 min) with vehicle (DMSO, 0.1 %) or test compounds and then stimulated with LPS for 4 h (TNF-α, IL-8), 18 h (IL-1β, IL-6, IL-10, MCP-1, IL-12 (p70), IL-23, IL-1ra). Cytokine and chemokine levels are given in pg per $1.4\times10^{6}$ PBMCs. Data for TNF-α and IL-1β are identical to Figure 4C. Mean + SEM and single data from n = 3-4 independent experiments. *P* values given vs. vehicle control; repeated measures one-way ANOVA of log data (IL-β, IL-6, IL-8, IL-10, MCP-1) or mixed-effects model (REML) of log data (TNF-α) + Dunnett *post hoc* tests or two-tailed paired student *t* tests of log data (IL-23, IL-1ra, IL-12(p70).

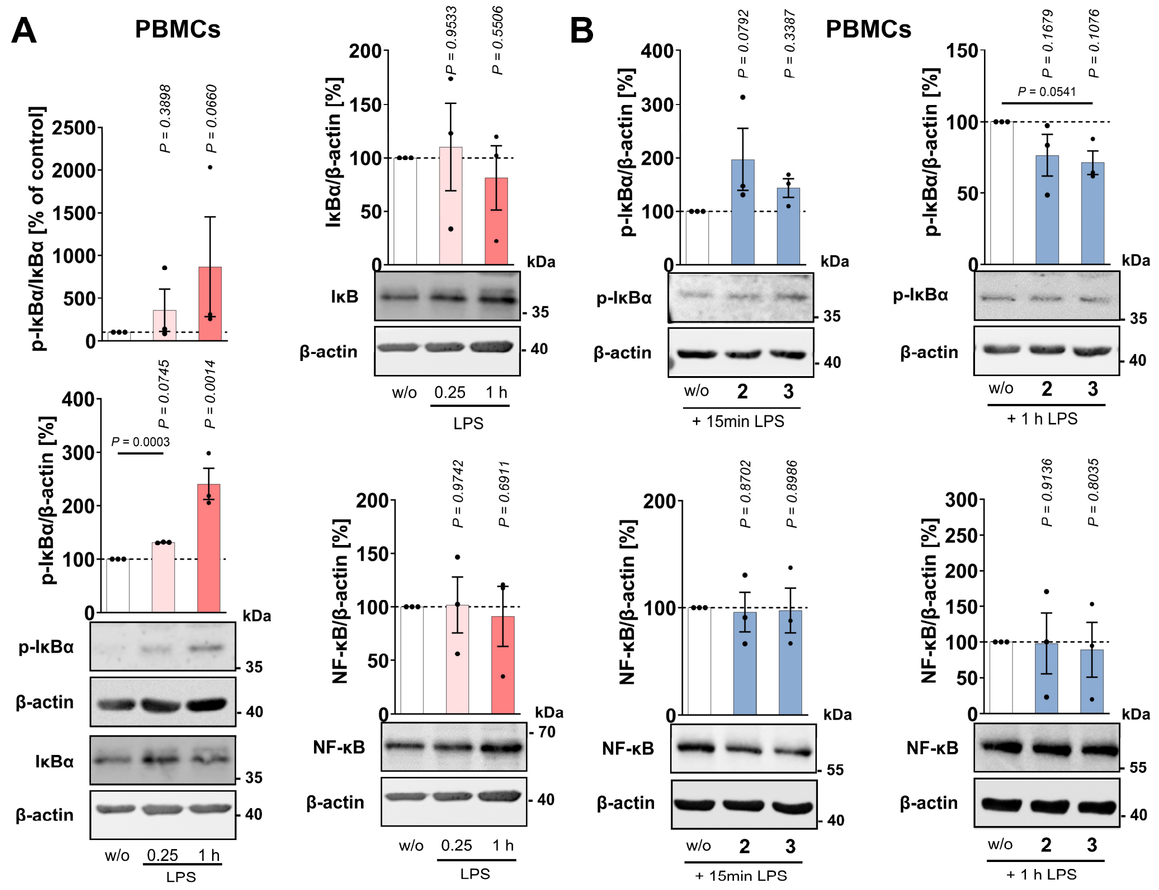


**Figure S10.** Effect of LPS, **2** and **3** on IκBα phosphorylation and IκBα and NF-κB expression. PBMCs were pretreated (30 min) with vehicle (DMSO, 0.1 %) or test compounds (10 µM) and then stimulated with LPS for 15 min or 1 h. A) Changes in IκBα phosphorylation and IκBα and NF-κB protein levels. Western blots are representative of three independent experiments. Uncropped blots are shown in Figure S11C, Supporting Information. B) Changes in p-IκB and NF-κB protein levels. Blots shown are identical to those shown in Figure 4B. Uncropped blots are shown in Figure S11A,B, Supporting Information. Mean + SEM and single data from n = 3 independent experiments. *P* values given vs. vehicle control; repeated measures one-way ANOVA (B) of log data (A) + Dunnett post hoc tests and two-tailed paired student *t* test for pairwise comparisons as indicated by bars (A,B).

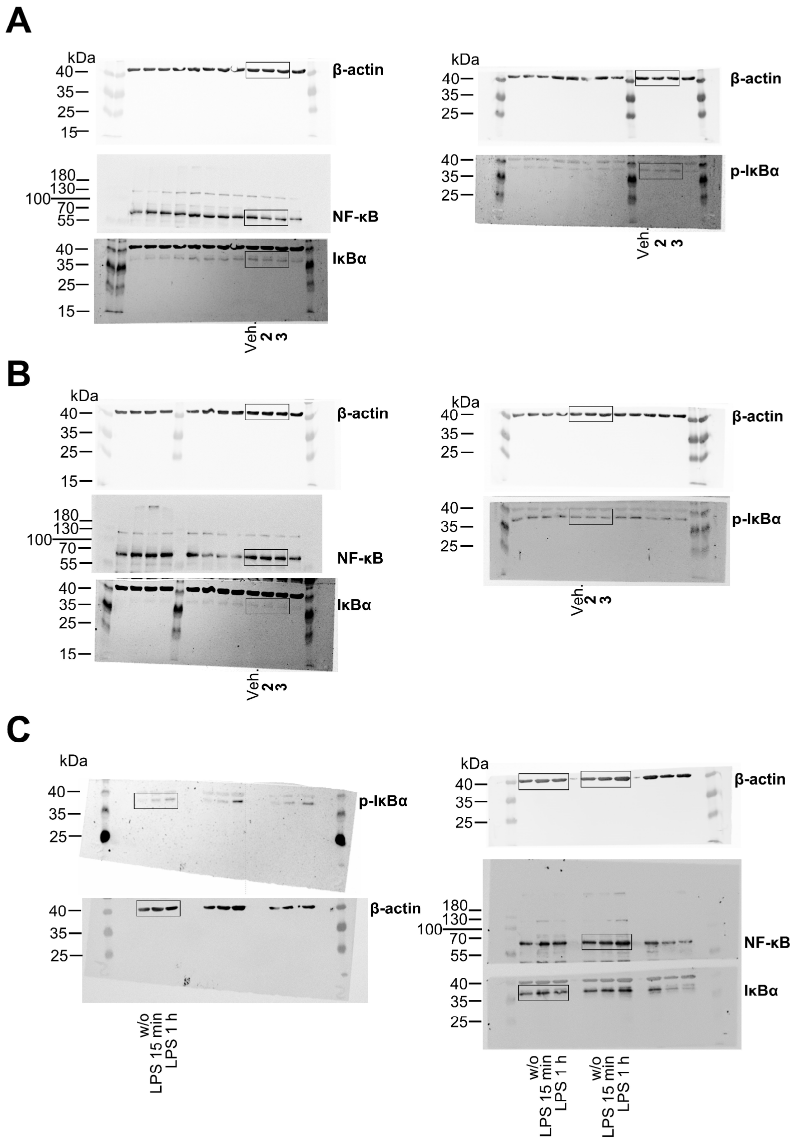


**Figure S11.** Uncropped versions of the blots shown in Figure 4B and Figure S10, Supporting Information. A,B) Effect of vehicle, **2** and **3** on NFκB, IκB, p-IκB, and β-actin protein levels in PBMCs stimulated with LPS for 15 min (A) or 1 h (B). C) Time-dependent effect of LPS on the levels of the above mentioned (phospho)proteins.

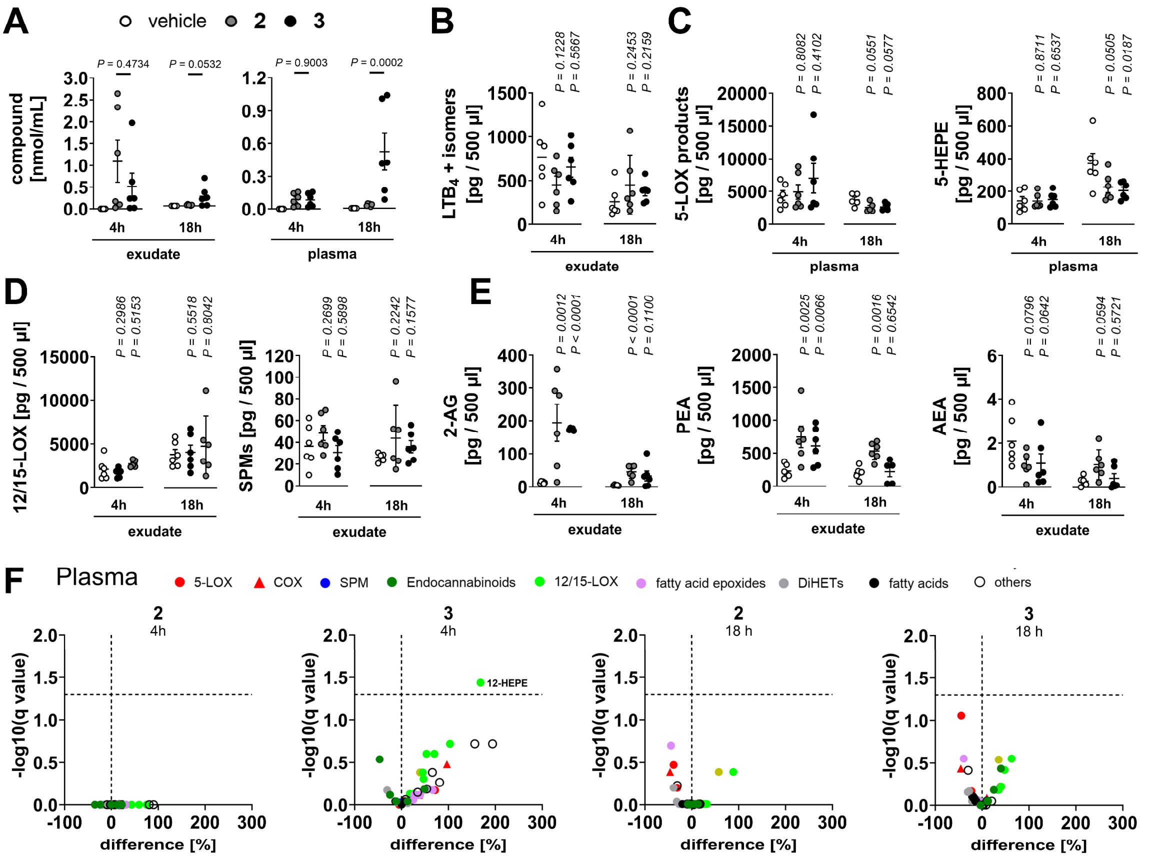


**Figure S12.** Exudate and plasma concentrations of **2** and **3** in murine peritonitis and effect on systemic and local lipid mediator profiles. Mice received vehicle (DMSO), **2** or **3** (10 mg/kg, i.p.) 30 min prior to zymosan injection and were sacrificed after 4 h or 18 h for the analysis of administered compound and lipid mediators in plasma and exudate. A) Concentrations of **2** or **3**. B) Total $LTB_4$ plus $LTB_4$ isomer levels in exudates. C) 5-LOX products and 5-HETE in plasma. D) 12/15-LOX products and SPMs in exudates. E) Selected endocannabinoids in exudates. F) Volcano plots showing the mean percentage difference relative to vehicle control and the negative $\log_{10}$(q value) calculated vs. vehicle control; two-tailed multiple paired Student *t* tests with correction for multiple comparisons (false discovery rate 5%). Mean (F) or mean + SEM and single data (A-E) from n = 5-6 (C,E,F), n = 6 (A,B,D). *P* values given vs. vehicle control; two-tailed unpaired student *t* test (B,C,D) of log data (A,E).

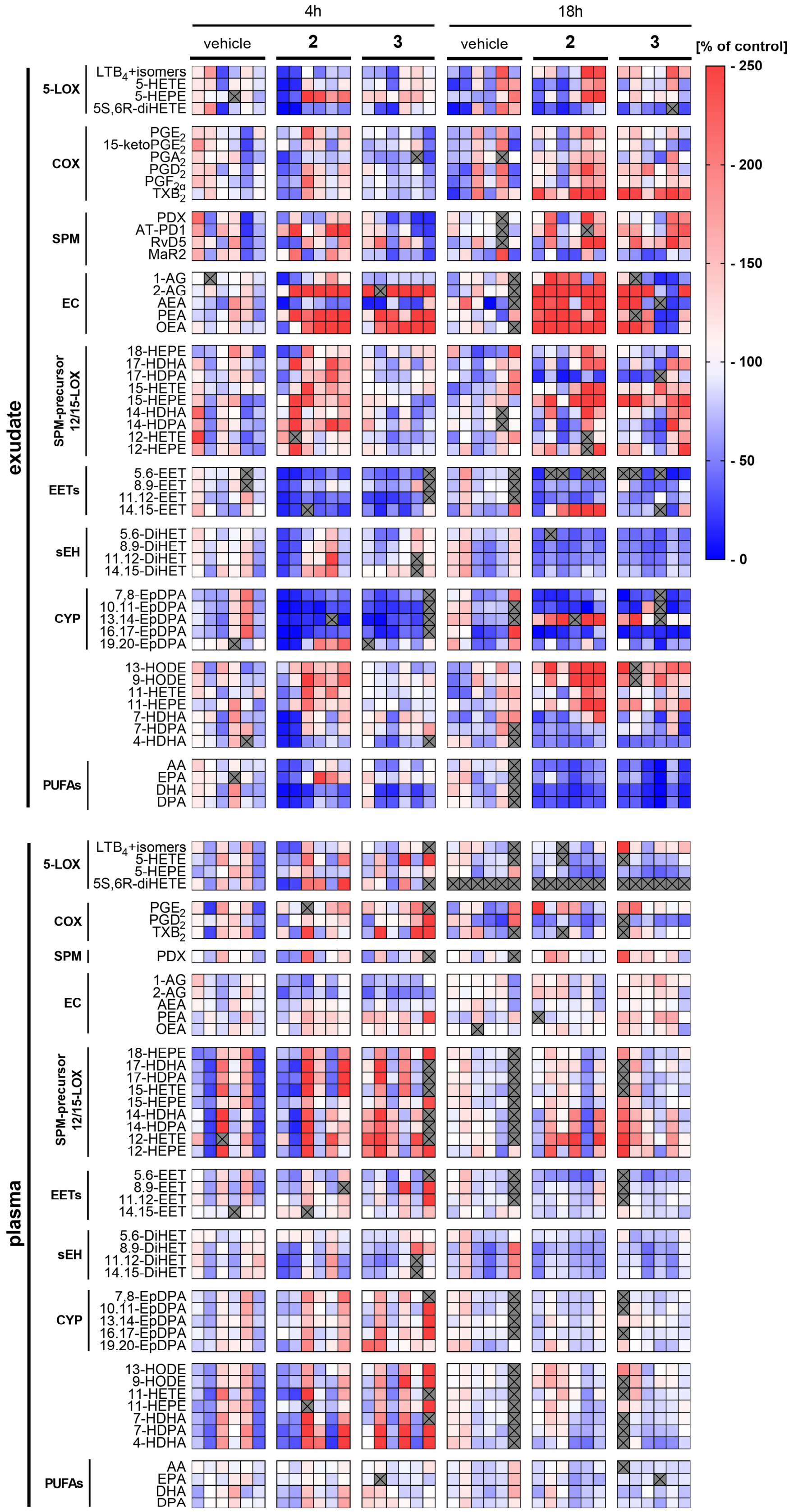


**Figure S13.** Detailed lipid mediator profile of murine exudates and plasma from zymosan-challenged mice upon administration of **2** or **3**. Mice received vehicle (DMSO), **2** or **3** (10 mg/kg, i.p.) 30 min prior to zymosan injection and were sacrificed after 4 h or 18 h for the analysis of lipid mediators in plasma and exudate**.** Heatmap showing percentage changes in lipid mediator levels relative to vehicle control for each animal (n = 2-6), with the crossed sections indicating significant outliers or non-detected species.

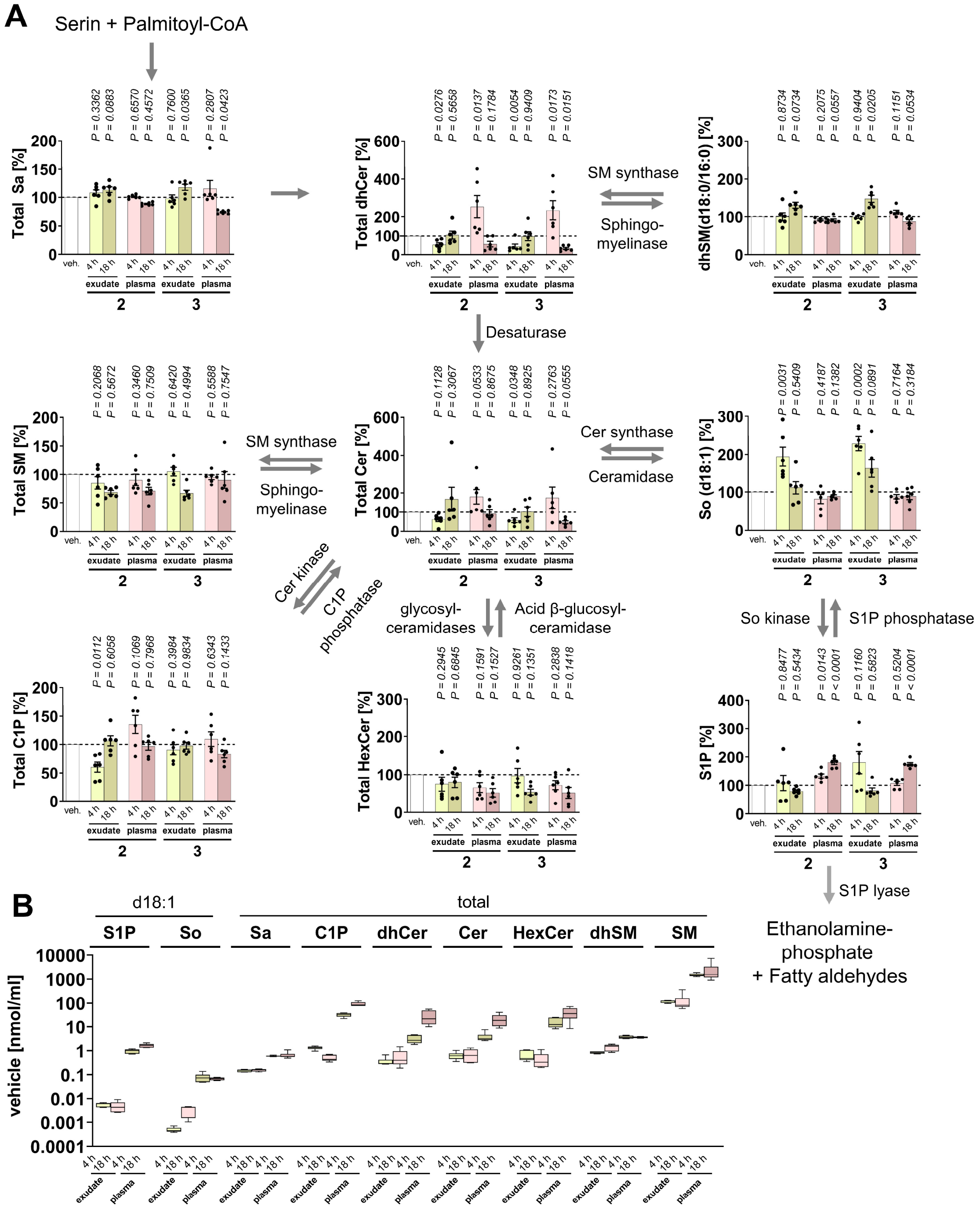


**Figure S14.** Effect of **2** and **3** on the exudate and plasma sphingolipidome in murine peritonitis. Mice received vehicle (DMSO), **2** or **3** (10 mg/kg, i.p.) 30 min prior to zymosan injection and were sacrificed after 4 h or 18 h for the analysis of sphingolipids in plasma and exudate**.** A) Percentage changes in the levels of sphingolipid subclasses relative to vehicle control and visualization of interconverting metabolic pathways and enzymes. Data for S1P are identical to Figure 5G. Abbreviations: Ceramide-1-phosphate (C1P), (dihydro)ceramide ((dh)Cer), (dihydro)sphingomyelin ((dh)SM), hexosylceramides (HexCer), sphingosine-1-phosphate (S1P), sphinganine (Sa), sphingosine (So). *P* values given vs. vehicle control; two-tailed unpaired Student *t* test of log data. B) Comparison of absolute concentrations of sphingolipid classes in vehicle control. Mean + SEM (B) and single data (A) from n = 5-6 mice.

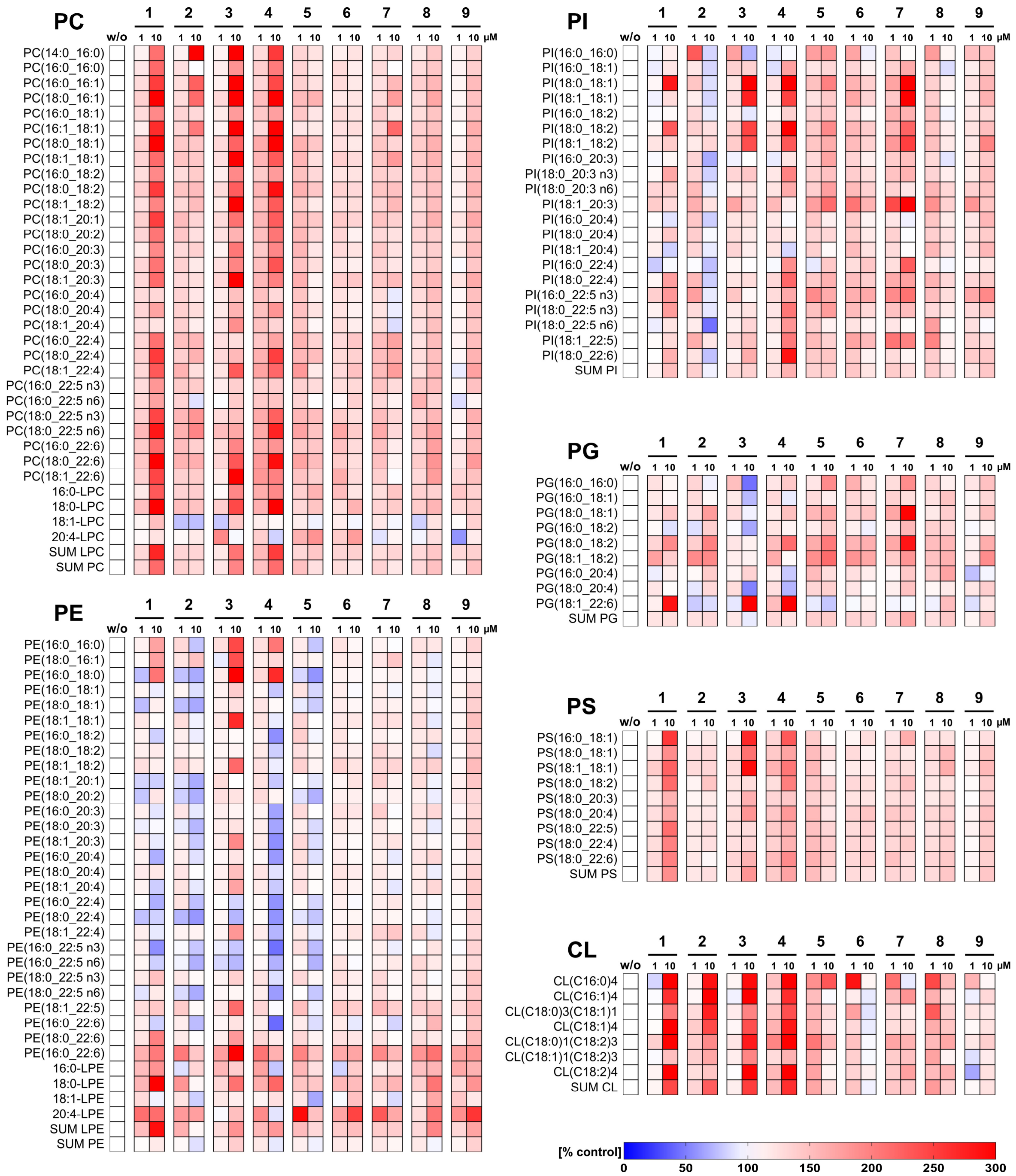


**Figure S15.** Effect of meroterpenoids on the cellular glycerophospholipidome of PBMCs. Cells were incubated with vehicle (DMSO, 0.1%) or test compounds for 48 h. Heatmaps showing percentage changes in the concentration of glycerophospholipid species and the subclass (‘SUM’) relative to vehicle control. Data is given as mean from n = 3 independent experiments. Data for subclasses are identical to Figure 6B.

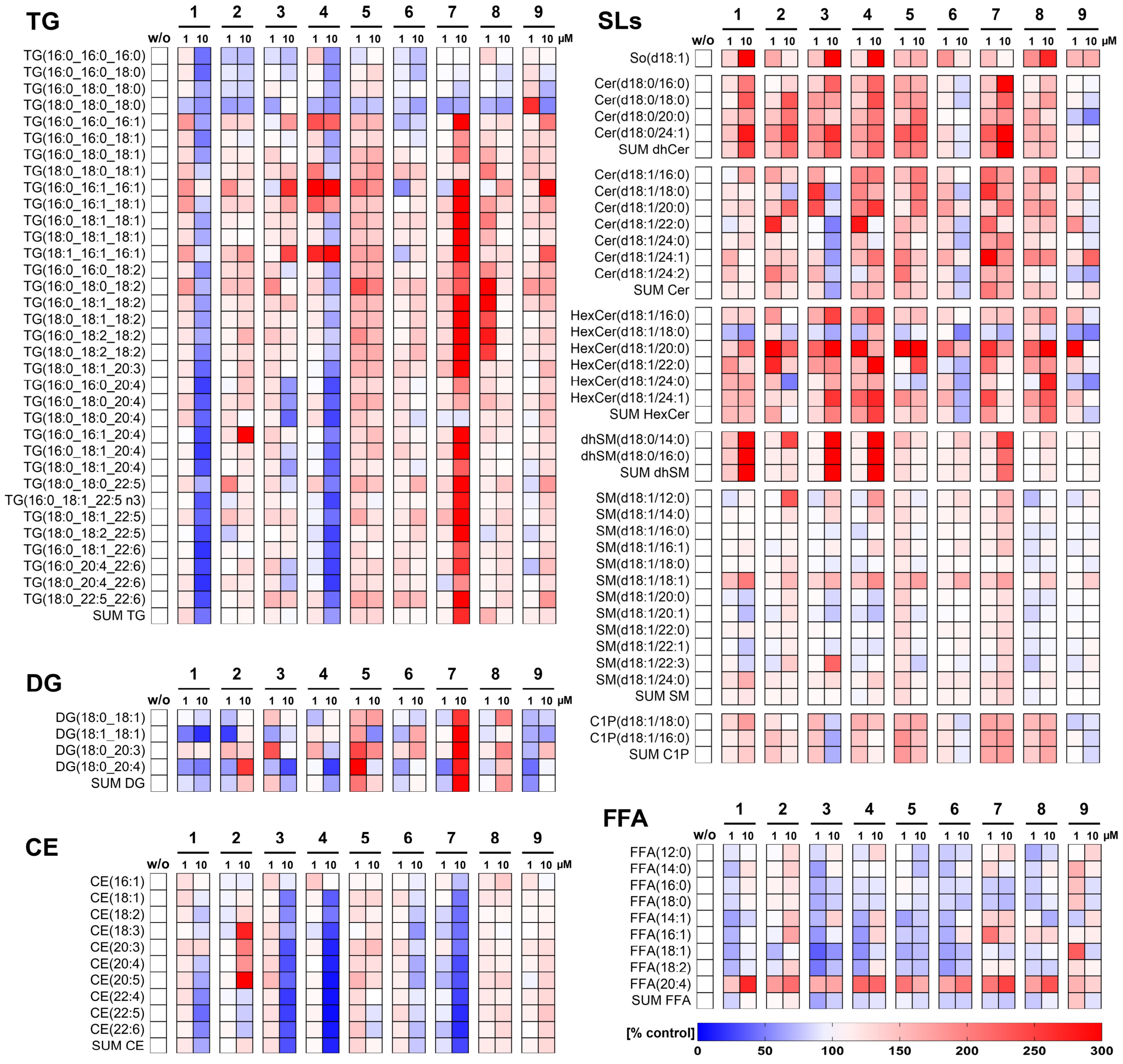

**Figure S16.** Effect of meroterpenoids on the composition of neutral lipids, sphingolipids (SLs), and FFAs in PBMCs. Cells were incubated with vehicle (DMSO, 0.1%) or test compounds for 48 h. Heatmaps showing percentage changes in the concentration of lipid species and the subclass ('SUM') relative to vehicle control. Data is given as mean from n = 3 independent experiments. Data for subclasses are identical to Figure 6B.

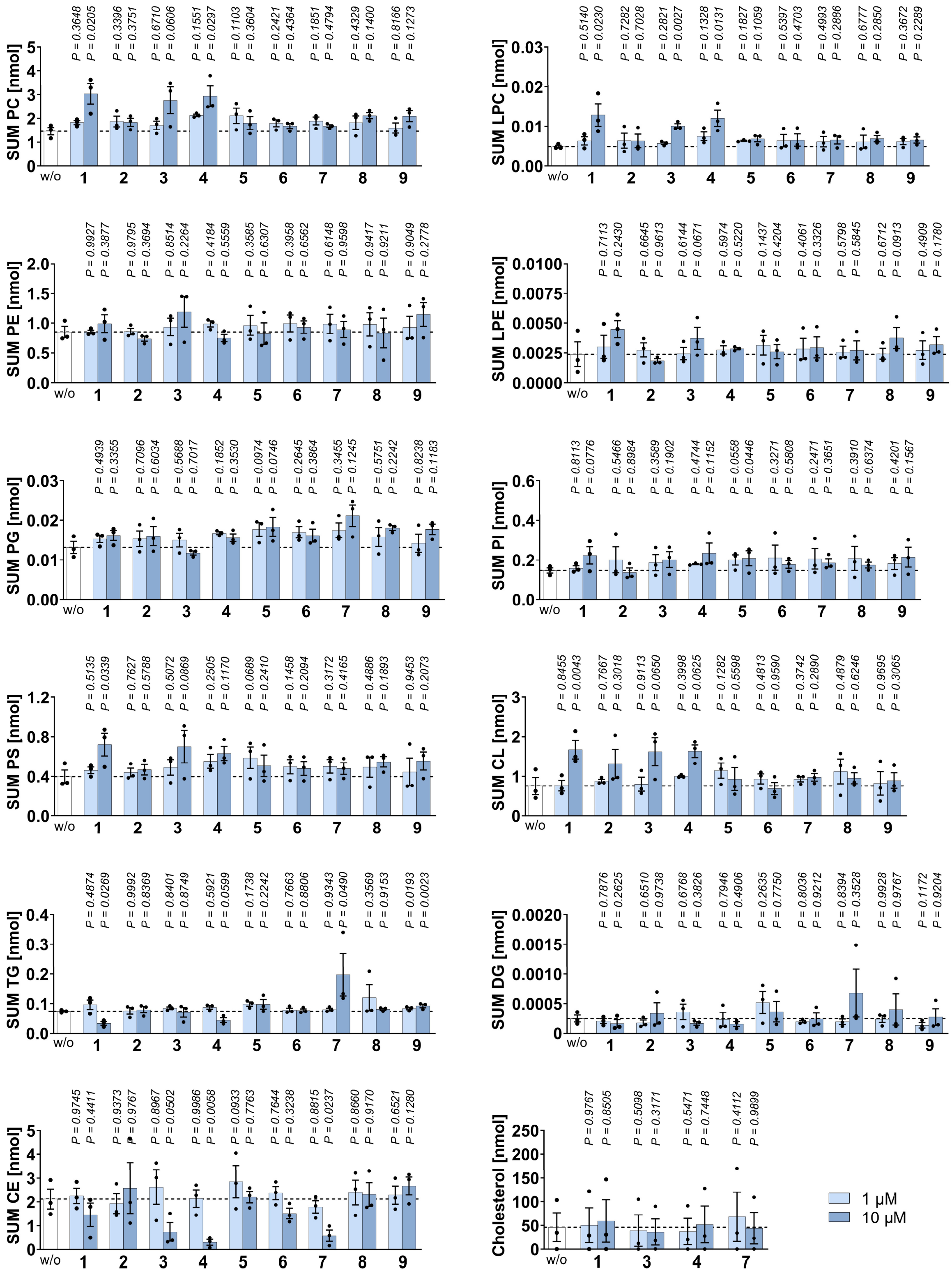

**Figure S17.** Absolute amounts of glycerophospholipid and neutral lipid subclasses in PBMCs treated with meroterpenoids. Cells were incubated with vehicle (DMSO, 0.1%) or test compounds for 48 h. The amounts of lipid classes are shown in nmol per 1×10$^6$ cells. Data are identical to Figure 6B (all lipids) and Figure 7E (LPC, LPE). Mean + SEM and single data from n = 3 independent experiments. *P* values given vs. vehicle control; repeated measures one-way ANOVA of log data + Dunnett *post hoc* tests.

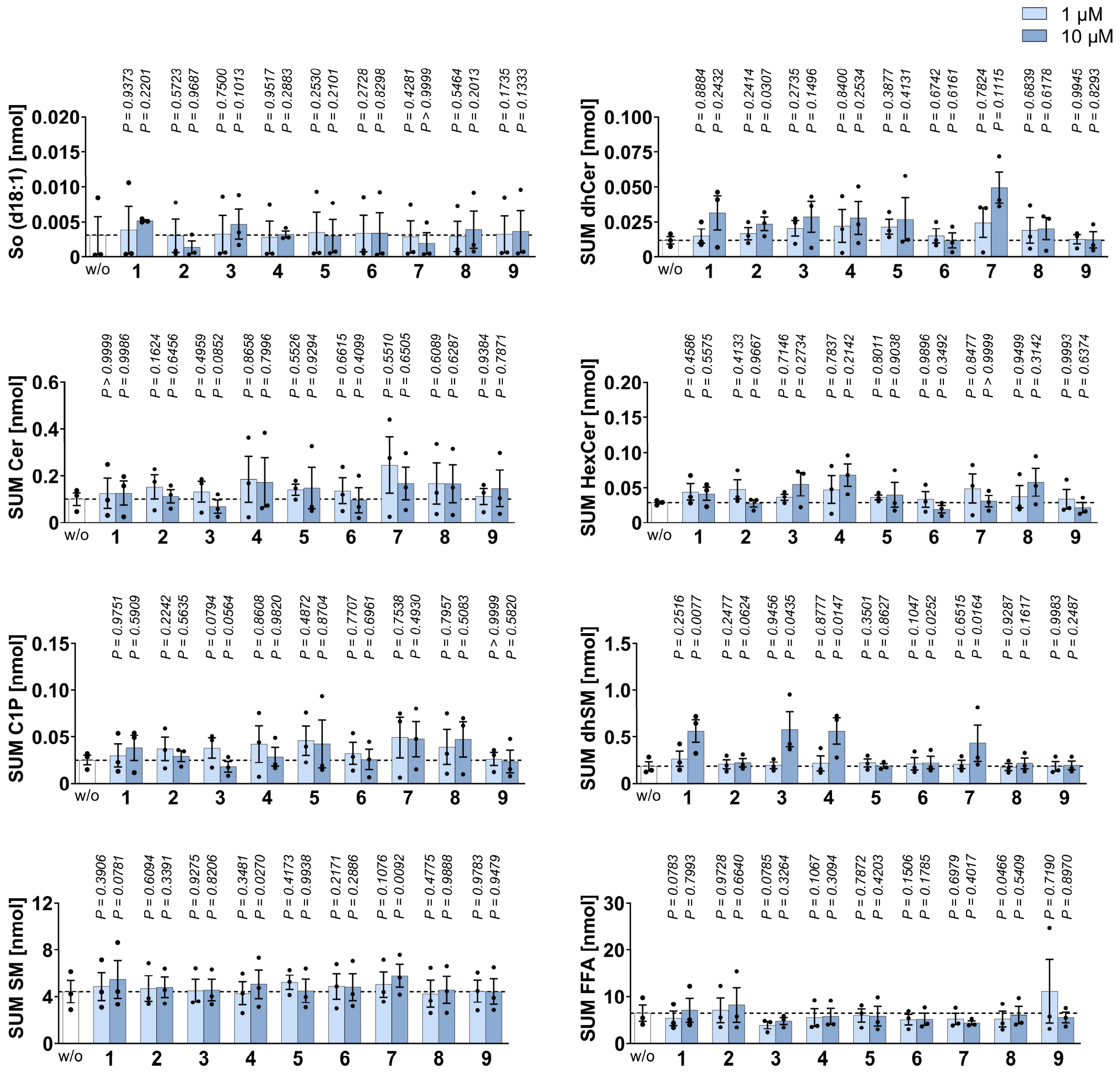


**Figure S18.** Absolute amounts of sphingolipid subclasses and FFAs in PBMCs treated with meroterpenoids. Cells were incubated with vehicle (DMSO, 0.1%) or test compounds for 48 h. The amounts of lipid classes are shown in nmol per $1\times10^6$ cells. Data are identical to Figure 6B (all lipids). Mean + SEM and single data from n = 3 independent experiments. *P* values given vs. vehicle control; repeated measures one-way ANOVA of log data + Dunnett *post hoc* tests.

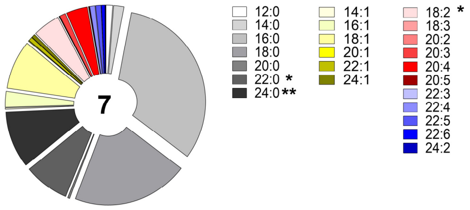


**Figure S19.** Proportion of fatty acids across all lipid species analyzed in **7**-treated PBMCs. Cells were incubated with vehicle (DMSO, 0.1%) or **7** (10 µM) for 48 h. The fatty acid composition of the vehicle control is shown in Figure 6C. Mean from n = 3 independent experiments. *P* values given vs. vehicle control; two-tailed paired Student *t* test.

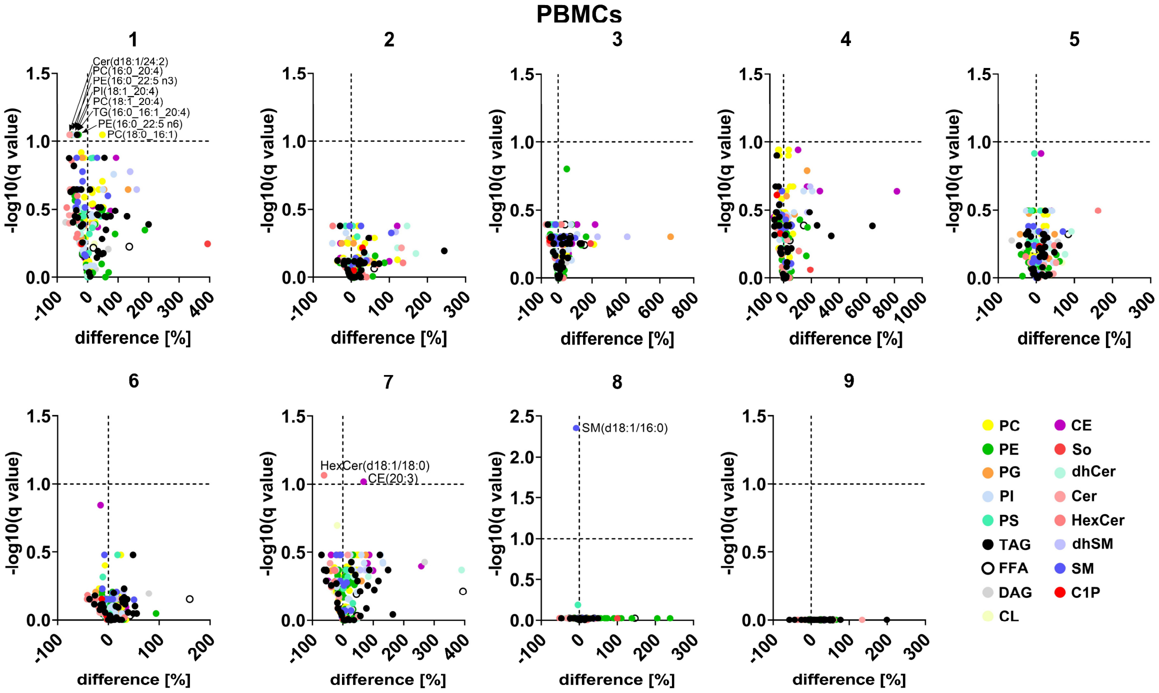


**Figure S20.** Extent and statistical significance of the effect of meroterpenoids on the relative lipid composition of PBMCs. Cells were incubated with vehicle (DMSO, 0.1%) or test compounds (10 µM) for 48 h. Volcano plot showing the mean percentage difference relative to vehicle control and the negative $\log_{10}$(q value) calculated vs. vehicle control from n = 3 independent experiments; two-tailed multiple paired Student *t* tests with correction for multiple comparisons (false discovery rate 10%).

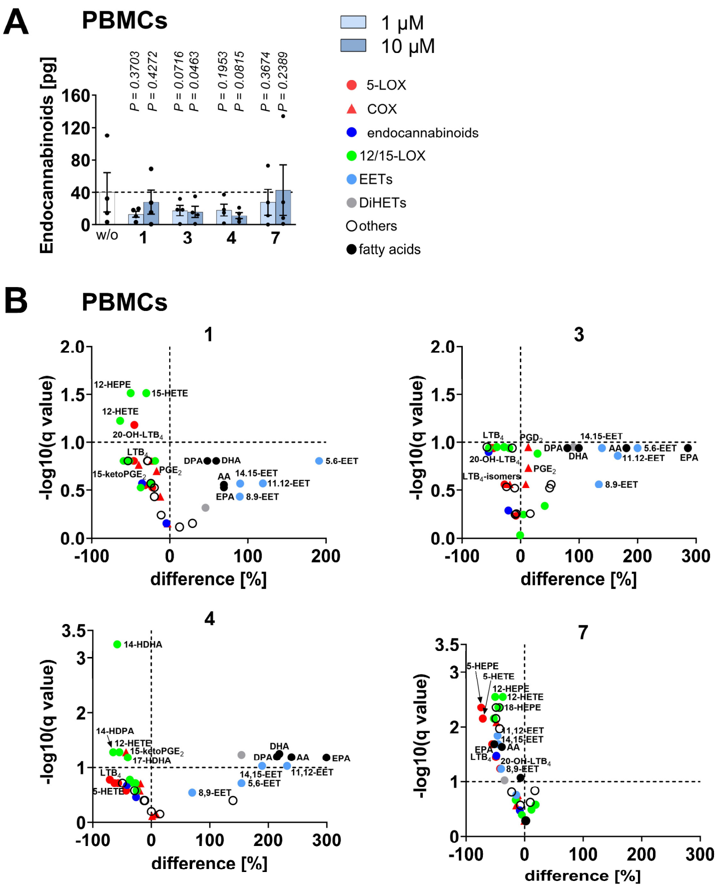

**Figure S21.** Changes in the capacity for lipid mediator biosynthesis following long-term exposure of PBMCs to **1**, **3**, **4** or **7**. Cells were pretreated (48 h) with vehicle (DMSO, 0.1%) or test compounds (10 µM unless otherwise noted), washed and activated with calcium ionophore (A23187, 10 min). A) Endocannabinoid levels in pg per $1 \times 10^6$ PBMCs. B) Volcano plot showing the mean percentage difference relative to vehicle control and the negative $\log_{10}$(q value) calculated vs. vehicle control; two-tailed multiple paired Student *t* tests with correction for multiple comparisons (false discovery rate 10%). Mean (B) or mean + SEM and single data (A) from n = 3-4 (B) or n = 4 (A) independent experiments. *P* values given vs. vehicle control; repeated measures one-way ANOVA of log data or mixed-effects model (REML) of log data + Dunnett *post hoc* tests (A).

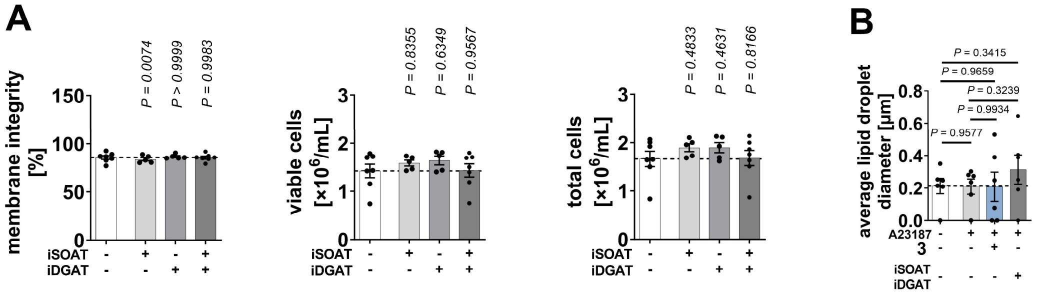


**Figure S22.** Combined SOAT and/or DGAT1/2 inhibition is compatible with PBMC survival and hardly affects lipid droplet diameter. A,B) PBMCs were treated with vehicle (DMSO, 0.1%), **3** (10 µM), TMP-153 (iSOAT, 500 nM), A-922500 (10 nM) + PF-06424439 (20 nM) (iDGAT), or TMP-153 (100 nM) + A-922500 (10 nM) + PF-06424439 (20 nM) (iSOAT + iDGAT) for 48 h. A) Effect on membrane integrity and viable and total cell number. Mean + SEM and single data from n = 5-7 independent experiments. B) PBMCs were subsequently treated with calcium ionophore (A23187, 5 min), and perilipin-2 was analyzed by immunofluorescence microscopy. The average lipid droplet diameter was calculated by dividing the total area of perilipin-2 staining by the number of individual perilipin-2 spots for each image. Two representative images (shown in Figure S23, Supporting Information) were analyzed per treatment group in n = 3 independent experiments. *P* values given vs. vehicle control (A) or as indicated by bars (B); mixed-effects model (REML) + Dunnett *post hoc* tests (A) or two-tailed unpaired Student *t* test for pairwise comparisons (B).

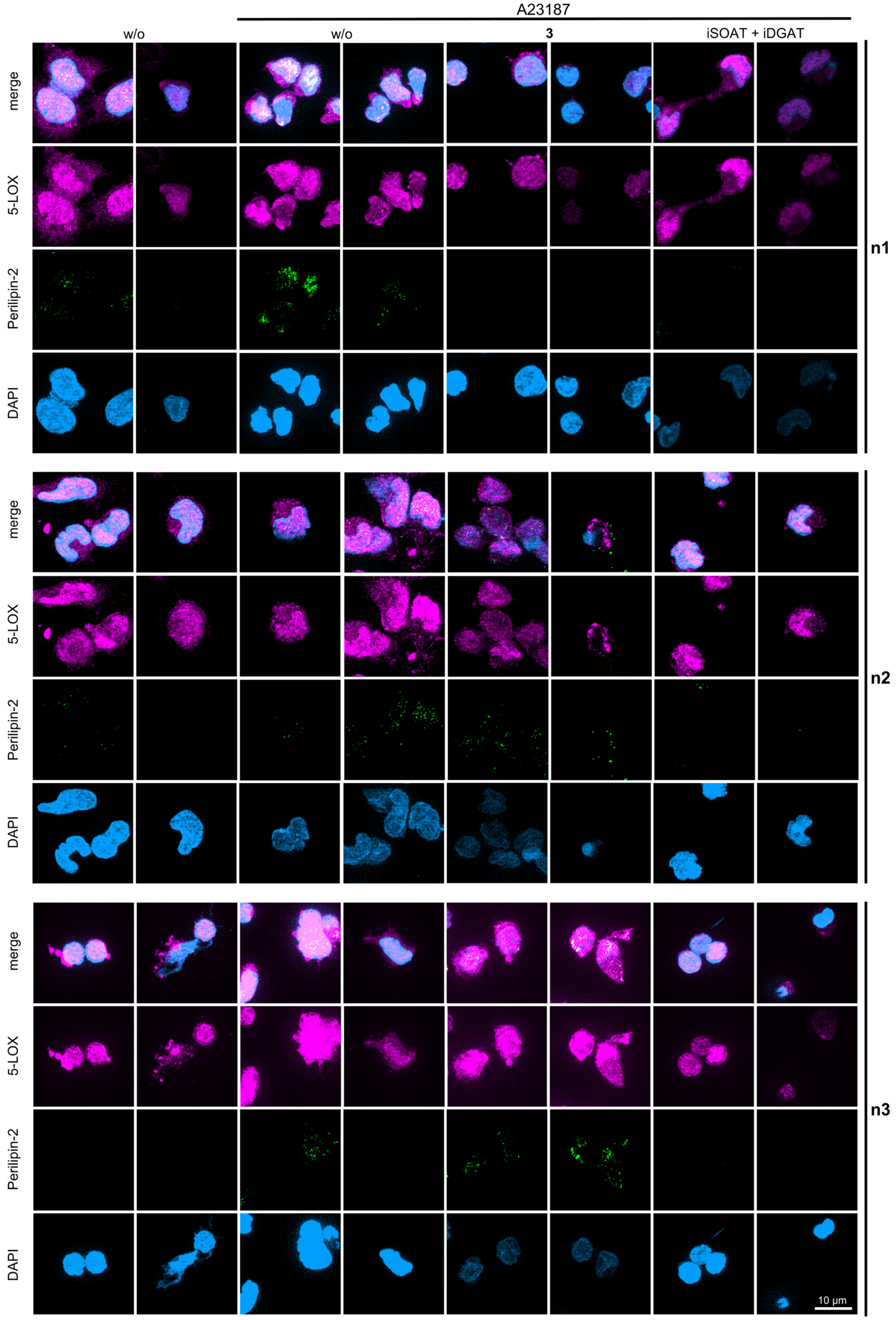
A23187
w/o
w/o
3
iSOAT + iDGAT
merge
5-LOX
Perilipin-2
DAPI
n1
n2
n3
10 µm

**Figure S23.** Effect of **3** and combined SOAT and DGAT inhibition on the abundance and subcellular distribution of perilipin-2 and 5-LOX in activated PBMCs. Cells were preincubated with vehicle (DMSO, 0.1%), **3** (10 µM) or TMP-153 (100 nM) + A-922500 (10 nM) + PF-06424439 (20 nM) (iSOAT + iDGAT) for 48 h before medium was exchanged and cells were treated with calcium ionophore (A23187, 5 min). Immunofluorescence images are stained for perilipin-2 (green, exposure time 35 ms), 5-LOX (violet, exposure time 200 ms), and nuclei (DAPI, light blue, exposure time 10 ms). Two representative images per treatment group are shown for n = 3 independent experiments.

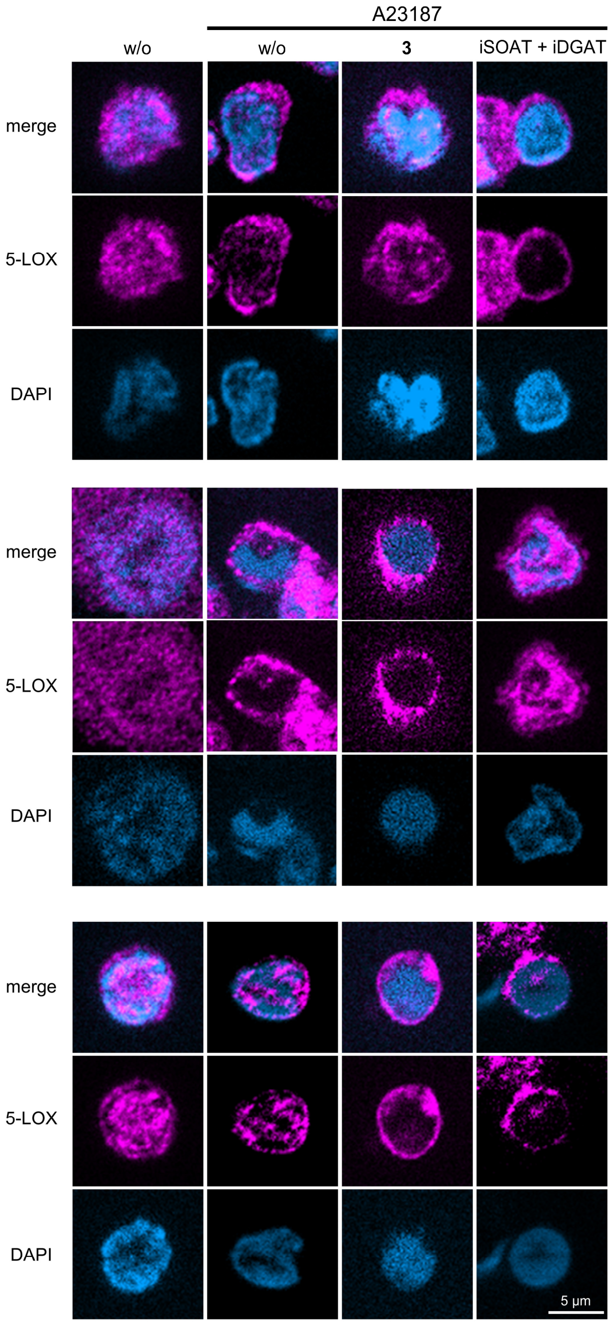


**Figure S24.** 5-LOX translocation in A23187-activated PBMCs is neither affected by the meroterpenoid **3** nor dual SOAT/DGAT inhibition. Cells were preincubated with vehicle (DMSO, 0.1%), **3** (10 µM) or TMP-153 (100 nM) + A-922500 (10 nM) + PF-06424439 (20 nM) (iSOAT + iDGAT) for 48 h before medium was exchanged and cells were treated with calcium ionophore (A23187, 5 min). Immunofluorescence images are stained for 5-LOX (violet, exposure time 200 ms) and nuclei (DAPI, light blue, exposure time 10 ms). Selected cells from single slices (Z = 1 µM) of n = 3 independent experiments are shown.

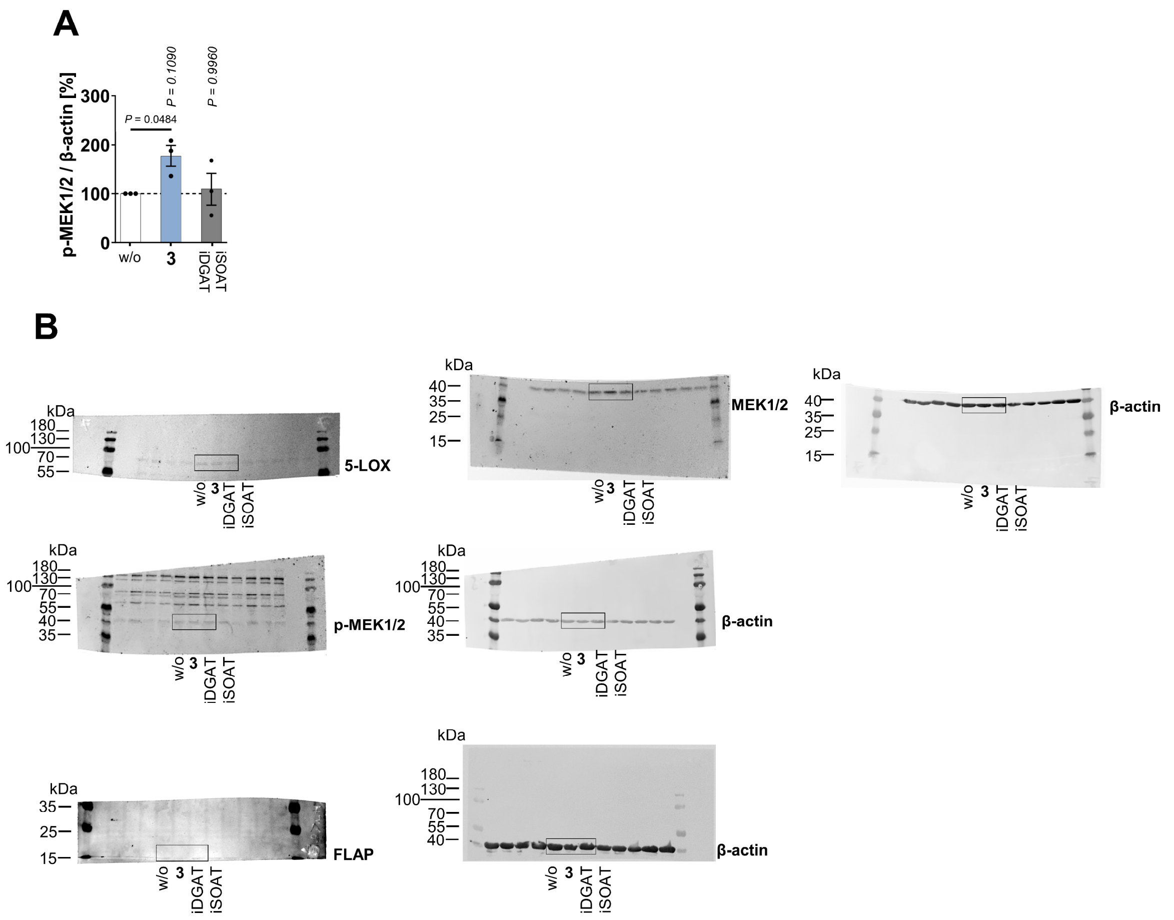


**Figure S25.** Effect of meroterpenoid **3** and combined SOAT and DGAT inhibition on the expression of key enzymes and regulatory factors in 5-LOX product biosynthesis. A,B) PBMCs were preincubated with vehicle (DMSO, 0.1%), **3** (10 µM), or TMP-153 (100 nM) + A-922500 (10 nM) + PF-06424439 (20 nM) (iSOAT + iDGAT) for 48 h. A) Effect of **3** and iSOAT + iDGAT on the expression of p-MEK1/2. Mean + SEM and single data from n = 3 independent experiments. *P* values given vs. vehicle control or as indicated by bars; repeated measures one-way ANOVA and two-tailed paired student *t* test for pairwise comparisons. B) Uncropped versions of the blots shown in Figure 8F.
